# A spatial map in the somatosensory cortex

**DOI:** 10.1101/473090

**Authors:** Xiaoyang Long, Sheng-Jia Zhang

## Abstract

Spatially selective firing in the forms of place cells, grid cells, boundary vector/border cells and head direction cells are the basic building blocks of a canonical spatial navigation system centered on the hippocampal-entorhinal complex. While head direction cells can be found throughout the brain, spatial tuning outside the hippocampal formation are often non-specific or conjunctive to other representations such as a reward. Although the precise mechanism of spatially selective activities is not understood, various studies show sensory inputs (particularly vision) heavily modulate spatial representation in the hippocampal-entorhinal circuit. To better understand the contribution from other sensory inputs in shaping spatial representation in the brain, we recorded from the primary somatosensory cortex in foraging rats. To our surprise, we were able to identify the full complement of spatial activity patterns reported in the hippocampal-entorhinal network, namely, place cells, head direction cells, boundary vector/border cells, grid cells and conjunctive cells. These newly identified somatosensory spatial cell types form a spatial map outside the hippocampal formation and support the hypothesis that location information is necessary for body representation in the somatosensory cortex, and may be analogous to spatially tuned representations in the motor cortex relating to the movement of body parts. Our findings are transformative in our understanding of how spatial information is used and utilized in the brain, as well as functional operations of the somatosensory cortex in the context of rehabilitation with brain-machine interfaces.

## Introduction

Spatial representation in the brain has been largely associated with the hippocampus and the entorhinal cortex ^1, 2^. Place cells ^3^ were originally discovered in the hippocampus, with boundary vector/border cells ^4, 5^, grid cells ^6^ and conjunctive cells ^7^ discovered later in the entorhinal cortex. Unlike the location-specific spatial cells in the hippocampal-entorhinal circuit, head direction cells ^8^ can be found in areas ranging from the diencephalon to thalamic, striatal and cortical regions ^9^.

Despite the focus on hippocampal-entorhinal contributions to spatial representation, location-dependent firing conjunctive to other sensory/cognitive (e.g. reward) information has been reported in sensory areas ^10–12^ and cortical regions heavily interconnected with the hippocampal-entorhinal system such as the septum, subicular complex, cingulate cortex, parietal cortex and retrosplenial cortex ^13–20^. In rodents, the only extra-parahippocampal region exhibiting apparent location-dependent firing restricted to space alone is the anterior claustrum, where stable, visually-anchored place cell- and border cell-like activities can be recorded ^21^. A preliminary report also suggests the dorsal geniculate nucleus may contain place cells similar to those found in the hippocampus ^22^. In humans, hexadirectional grid-like signals have been observed in many brain regions outside the hippocampal formation including the posterior cingulate cortex, the medial prefrontal cortex, the retrosplenial cortex, the medial parietal cortex and the frontal cortex ^23–26^ while grid-cell-like neuronal representations were identified with intracranial electroencephalography recordings on presurgical epilepsy patients and fMRI studies in the human entorhinal cortex ^27–29^. Computational modelling also predicted the existence of grid cell-like neurons in all cortical regions throughout the neocortex ^30^. These results therefore indicate that spatial tuning is much more distributed into multiple cortical domains beyond the classical hippocampal formation than we previously thought.

The exact physiological mechanism of spatial selectivity in the hippocampal-entorhinal system is still incompletely understood. Lesions to either area severely affect, but do not abolish spatially selective activities in the other ^31, 32^. Past studies have shown manipulation of visual cues ^33, 34^, olfaction ^35^ and vibrissae inputs ^36^ all impact on hippocampal place cell activity, yet none is necessary. Recent theoretical work has suggested the somatosensory area may contain higher-order presentations of the body itself beyond elementary sensory information, of which location presentation is integral in a “body simulation” model of somatosensation ^37^. While there are place cell-like activities in the wheelchair-seated monkey sensorimotor cortices ^38^, it is unclear if these spatial representations are comparable to their freely moving rodent equivalents. Based on the assumption that somatosensation should play a major role in sensory inputs shaping parahippocampal spatial activity ^39, 40^ and spatial presentation has been theorized to be crucial for body representation ^37^, we sought to identify spatially selective activities in the primary somatosensory cortex (S1). Remarkably, we were able to detect all spatial cell types reported in the hippocampal-entorhinal system ^1, 2^ within the S1. We show these spatial cell types within this newly discovered somatosensory spatial representation system share similar properties to their parahippocampal counterparts in terms of stability and response to environmental manipulations ^41^.

## Results

### Extracellular recording from the somatosensory cortex

We obtained neurophysiological recordings in freely behaving rats by using tetrodes mounted on movable microdrives. We recorded 2025 putative single units from eight implanted rats across 287 recording sessions. Recordings were obtained mainly from layers IV to VI within the S1 area (Fig. 1a and Fig. S1), as confirmed by the histological reconstruction. The tetrodes were generally lowered in steps of either 25 μm or 50 μm per day. The recording sessions lasted between 10 to 30 minutes to ensure sufficient coverage of the whole arena, which was facilitated by foraging food pellets intermittently thrown into the arena. Spike sorting was performed manually offline using TINT (Axona, St. Albans, U.K.). The separation of the clusters and waveforms was quantitatively measured with an isolation distance and the L-ratio ^42^ (Fig. S2).

**Fig. 1.**
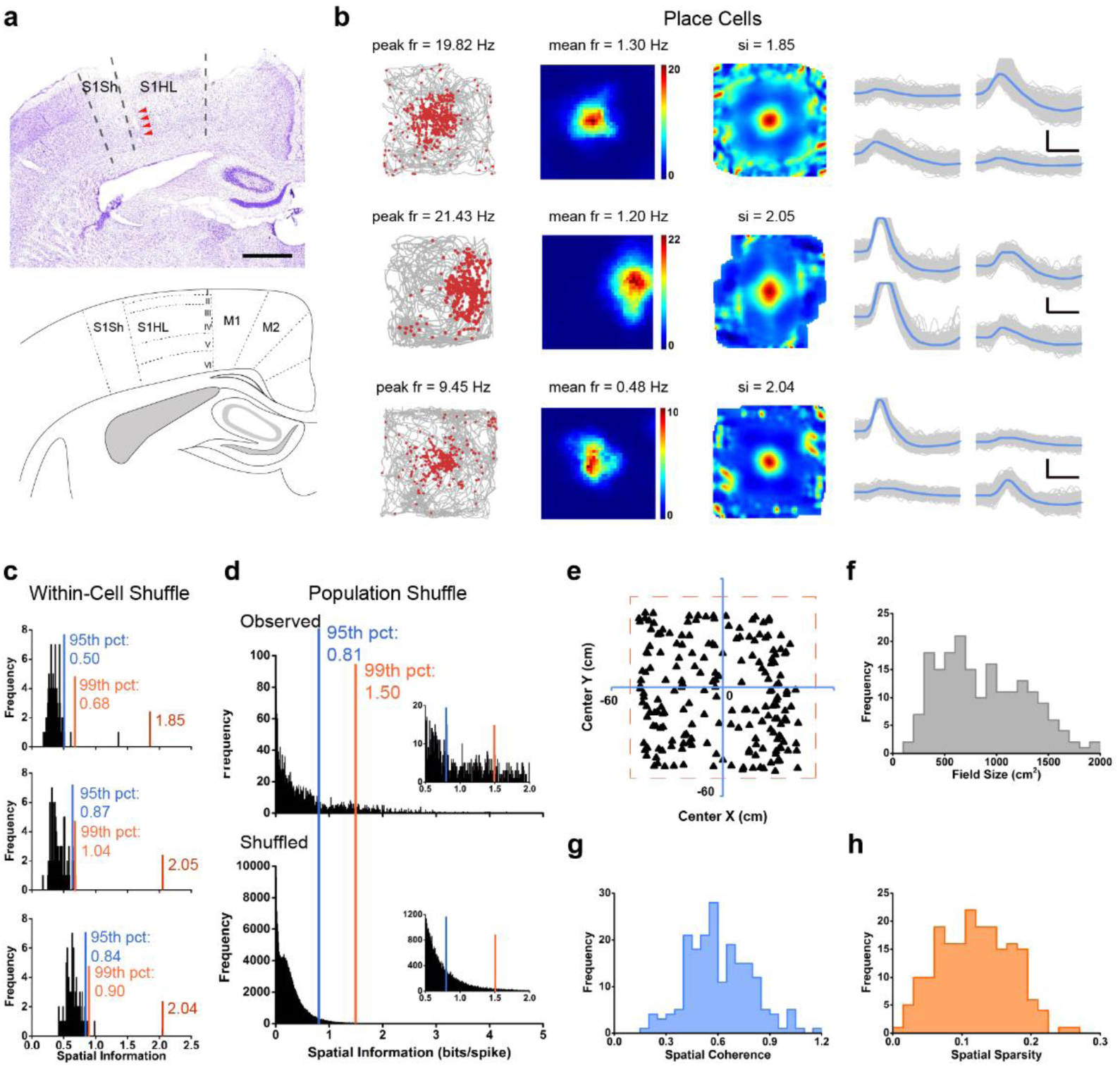
Place cells in the somatosensory cortex. **(a)** A Nissl-stained coronal section (upper) showing tetrode tracking trajectory (arrowheads) through all of six layers across the rat primary somatosensory cortex. Broken dashes depict the boundaries of the hindlimb (S1HL) and shoulder (S1Sh) regions of the primary somatosensory cortex (S1). The bottom panel shows the schematic delimitation of the different layers and sub-regions of the primary somatosensory cortex. Scale bar, 1 mm. (**b)** Trajectory (grey line) with superimposed spike locations (red dots) (left column); heat maps of spatial firing rate (middle left column) and autocorrelation (middle right column) are color-coded with dark blue indicating minimal firing rate and dark red indicating maximal firing rate. The scale of the autocorrelation maps is twice that of the spatial firing rate maps. Peak firing rate (fr), mean firing rate (fr) and spatial information (si) for each representative cell are labelled at the top of the panels. Spike waveforms on four electrodes are shown on the right column. Scale bar, 150 µV, 300 µs. (**c)** Distribution of within-cell shuffled spatial information for three representative somatosensory place cells. The orange and blue stippled lines mark the 99^th^ and the 95^th^ percentile significance level of each randomly shuffled distribution, respectively. The red line indicates the observed spatial information. (**d)** Distribution of spatial information for all isolated somatosensory units. The top panel shows the distribution for observed spatial information. The bottom panel shows the distribution for randomly shuffled data from the same sample. The orange and blue stippled lines mark the 99^th^ and the 95^th^ percentile significance level of each randomly shuffled distribution, respectively. A zoomed panel shows the magnification of the specified area marked by the red dashed rectangle. (**e**) The uniformly areal distribution of various place firing fields relative to the field center. (**f-h**) The population histograms of place firing field sizes (**f**), spatial coherence (**g**) and spatial sparsity (**h**) for all of 195 identified somatosensory place cells.

We applied the same analysis described in previous work ^7, 17, 43^ on 2025 somatosensory cells and 392 met the criteria for various types (e.g. place, head direction, boundary vector/border and grid cells) of spatial tuning (Fig. S3). The number of each functionally distinct somatosensory spatial cell type from individual implanted animals was summarized in the supplementary Table S1. No significant differences were observed among spatial information, mean vector length and grid score across different animals (ANOVA test, *P* = 0.38, 0.08 and 0.18, respectively, Figs. S4a, b and d). Both border score and grid spacing remained similar across interindividual animals except that they are slightly different in one of eight animals (ANOVA test, * *P* < 0.05, Figs. S4c-d). The distribution of different spatial cell types across different animals suggested, at least between layers IV and VI, there was a tendency for head direction cells to avoid layer VI and the presence of spatially tuned somatosensory cells was sparse in superficial layer V (Figs. S3b-f). Place cells appeared to predominate deeper layers (Figs. S3a-c).

### Place cells in the somatosensory cortex

As illustrated in Fig. 1, somatosensory cells possessing place cell-like properties were the most abundant spatial cell type encountered (Figs. S3a-c). Spatially selective firing patterns similar to those originally described for hippocampal place cells ^3^ can be observed (Fig. 1b and Fig. S5). These S1 place cells were classified if their spatial information (SI) score ^44^ exceeds a stringent 99^th^ percentile threshold (SI > 1.50 cut-off) from the population-based distribution of shuffled rate maps (Fig. 1d). 195 out of the 2025 S1 neurons (9.63%) met criterion and thus are referred to as “place cells” from here on. These place cells also passed the spatial threshold for the within-cell shuffling (Fig. 1c and Fig. S6a), which is less stringent than the population shuffling. The within-cell shuffling outcome is also true of other types of somatosensory spatial cells (Figs. S6b-d). Accordingly, we used the population shuffling for defining all different types of somatosensory spatial cells throughout this study. Furthermore, we applied the maximum-likelihood approach ^45, 46^ to evaluate inhomogeneous sampling biases and found that somatosensory place cells showed a robust locational effect, which was not altered by the correction algorithm (Figs. S7a-c). The average SI in bits per spike of all 195 classified S1 place cells was 2.19 ± 0.04 (mean ± s.e.m., all data are shown as mean ± s.e.m. unless otherwise indicated). This percentage was significantly higher than expected by random selection from the entire shuffled population (Fig. 1d; *Z* = 39.03, *P* < 0.001; binomial test with expected *P_0_* of 0.01 among large samples).

We noted that if a 95^th^ percentile criterion was applied, 511 out of the 2025 S1 neurons (24.00%) would be classified as place cells (average SI of 1.59 ± 0.03; cut-off = .81). This percentage was substantially higher than expected with a random selection from the entire shuffled population (Fig. 1d; *Z* = 109.61, *P* < 0.001; binomial test with expected *P_0_* of 0.01 among large samples). We found that the percentage of identified S1 place cells is lower within the somatosensory cortex than that of hippocampal place cells ^47^. The average peak firing rate and the mean firing rate of identified somatosensory location-specific place cells were 10.32 ± 0.37 Hz and 0.70 ± 0.04 Hz (Fig. S8a), respectively. The histograms of peak-to-peak amplitude and peak-to-trough spike width were shown in Fig. S8e. Like a mixture of a single place field and multiple place fields identified from the hippocampal place cells ^48^, we found that the proportion of multi-field place cells was 25.64%. The average somatosensory spatial firing field size was 891.82 ± 32.65 cm^2^ (Fig. 1f). To quantify the smoothness of recorded somatosensory place fields, we calculated the spatial coherence by computing the mean correlation between the firing rate of each bin and the averaged firing rate of the 8 neighboring bins from unsmoothed spatial firing rate maps ^34^. For 195 recorded somatosensory place cells, the average spatial coherence was 0.61 ± 0.01 (Fig. 1g). Furthermore, we also calculated spatial sparsity to evaluate the extent of spatial information in each spike discharged by S1 place cells ^44^. The spatial sparsity measure of S1 place cells was 0.12 ± 0.01 (Fig. 1h). Besides, S1 place cells can be found at any position within the arena for the uniformity testing based on the Friedman-Rafsky’s MST test (Fig. 1e, Friedman-Rafsky’s MST test, *P* = 0.42, two-tailed *z*-test) as in the hippocampus ^49^.

Place cell activity was defined not only by its location-specific firing but also by the stability of its spatial representation. To this end, we next evaluated the spatial stability of somatosensory place fields by computing the spatial correlations between single intra-trial behavioral sessions. The average spatial correlation for S1 place cells was 0.64 ± 0.01 (Fig. S9d) between the two halves of a recording session (Figs. S9a-c), which remain stable in extended recording sessions up to 30 min (Figs. S10a-c). Hippocampal place cell activities were known to be heavily modulated by visual cues ^33, 34^, and were known to rotate with the positioning of visual cues ^34^. We performed similar experiments to test if S1 place cell activities were modulated by visual cues by moving it 90° from its original position and back. Remarkably, the relative position of S1 place fields did indeed rotate with the placement of the visual cue and remained stable after the cue was moved back to its original position (Figs. S11a-c). Furthermore, as hippocampal place codes were reorganized to remap under different geometric environments ^50, 51^, we also found that S1 place fields remapped between a square box and a circular box in the same recording room with changed firing rate and firing location (Figs. S12a-c).

To measure the animal’s running speed within and outside the firing fields of S1 place cells, we calculated the average “in-field” and “out-field” running speed (> 2.5 cm/s) for 195 identified place cells in S1. We quantified the average running speed across all the spatial bins within the firing field of the place cell (“in-field” running speed) as well as outside the firing field of the place cell (“out-field” running speed) (Fig. S13a). We found that there was no significant difference between the average “in-field” and “out-field” running speed (14.58 ± 0.25 cm/s vs 15.00 ± 0.18 cm/s, *n* = 195, *P* = 0.10, two-tailed paired *t*-test, Fig. S13b). To evaluate the effect of the animal’s running speed on spatial responses, we calculated the spatial information scores of S1 place cells for running speed above 2.5 cm/s and below 2.5 cm/s, respectively. We found that the place fields during inactive mobility are less prevalent with lower spatial information score than those for active exploring (Figs. S14a-c). Similar results were observed for S1 border cells (Figs. S27a-c) and S1 grid cells (Figs. S30a-c). As for S1 head direction cells, it appeared that head directional responses were less influenced with immobility than other three S1 spatial cell types (Figs. S21a-c).

### Head direction cells in the somatosensory cortex

Head direction cells discharge only when the animal’s head is at a particular angle respective to the environment ^8^. To classify the preferred firing angle of each cell, the mean vector length was used to compute head direction tuning ^7^. Cells were categorized as head direction cells if the mean vector length exceeded the 99^th^ percentile of the total distribution of the shuffling procedure from the entire pool of putative single cells (Fig. 2c). 80 cells (3.95% of all recorded cells) met the criterion, showing significant modulation with the animal’s head direction (Fig. 2a and Fig. S15), with the average mean vector length found to be 0.53 ± 0.01 and the mean firing rate to be 0.96 ± 0.11 (Fig. S8b). The histograms of peak-to-peak amplitude and peak-to-trough spike width were displayed in Fig. S8f. Besides, the within-cell shuffling for those three representative somatosensory head direction cells generated similar results to the population shuffling (Fig. 2b). This percentage is significantly higher than expected by random selection from the entire shuffled population (Fig. 2c; *Z* = 13.34, *P* < 0.001; binomial test with expected *P_0_* of 0.01 among large samples). Notably, 352/2025 identified S1 neurons (17.38%) fulfilled the lower 95^th^ percentile criterion with the threshold of mean vector length being 0.26. To correct for the possible effects of direction and location for identified head direction cells, we applied the maximum-likelihood correction approach ^45, 46^ and found that the somatosensory head directional responses remain unaltered from the corrected directional firing rate histograms (Figs. S16a-c).

**Fig. 2.**
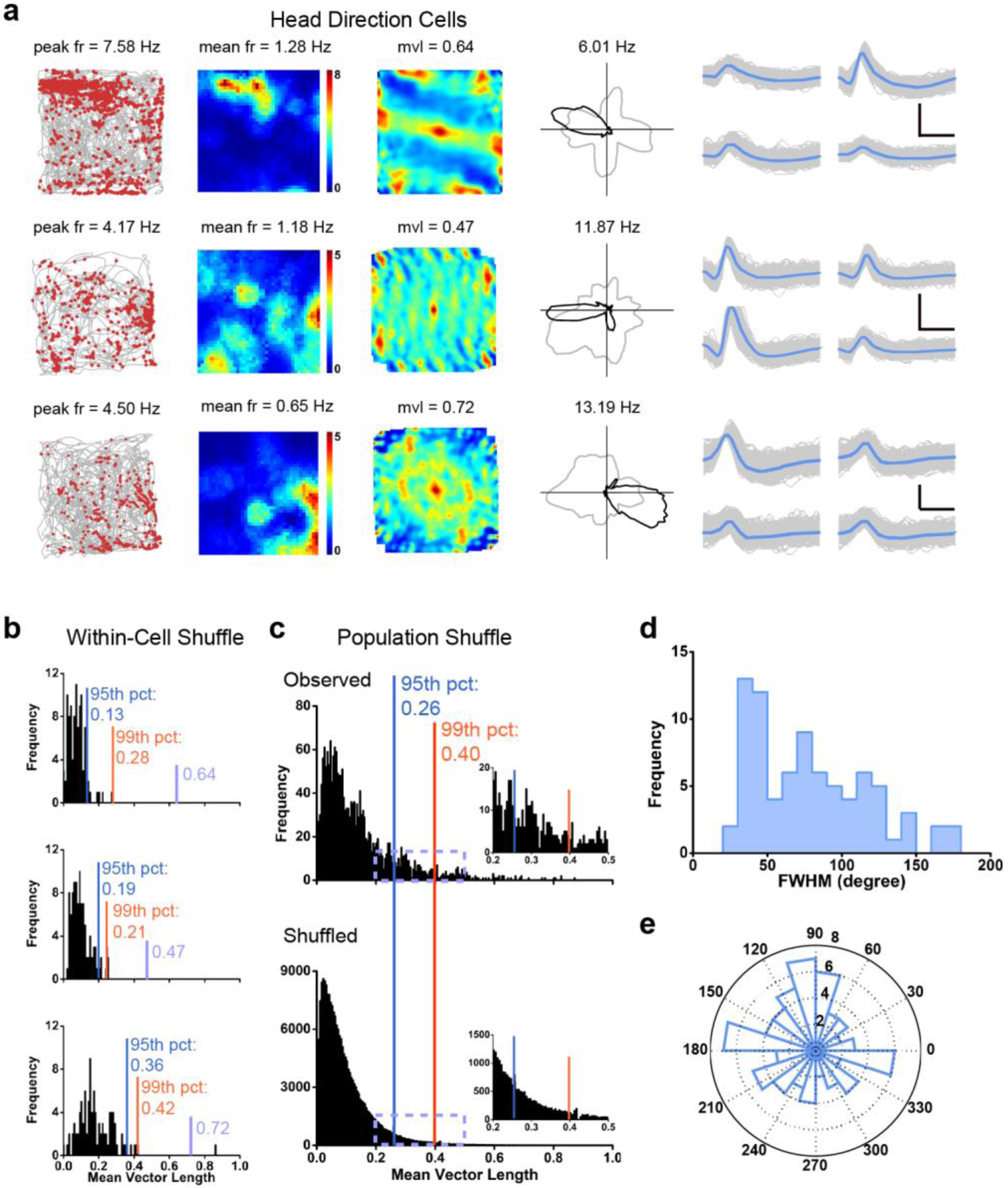
Head direction cells recorded from the somatosensory cortex. **(a)** Three examples of somatosensory head direction cells. Trajectory (grey line) with superimposed spike locations (red dots) (left column); spatial firing rate maps (middle left column), autocorrelation diagrams (middle right column) and head direction tuning curves (black) plotted against dwell-time polar plot (grey) (right column). Firing rate is color-coded with dark blue indicating minimal firing rate and dark red indicating maximal firing rate. The scale of the autocorrelation maps is twice that of the spatial firing rate maps. Peak firing rate (fr), mean firing rate (fr), mean vector length (mvl) and angular peak rate for each representative head direction cell are labelled at the top of the panels. The directional plots show strong head direction tuning. Spike waveforms on four electrodes are shown on the right column. Scale bar, 150 µV, 300 µs. (**b)** Distribution of within-cell shuffled mean vector length for three representative somatosensory head direction cells. The orange and blue stippled lines mark the 99^th^ and the 95^th^ percentile significance level of each randomly shuffled distribution, respectively. The purple line indicates the observed mean vector length. (**c**) Distribution of mean vector length for all identified somatosensory units. The top panel shows the distribution for observed values. The bottom panel shows the distribution for randomly shuffled data from the same sample. The orange and blue stippled lines mark the 99^th^ and the 95^th^ percentile significance level of each randomly shuffled distribution, respectively. A zoomed panel shows the magnification of the specified area marked by the purple dashed rectangle. (**d**) Distribution of head directional tuning width. (**e**) Preferred direction of recorded head direction cells from S1. The polar plot shows the distribution of the peak firing direction of all identified somatosensory head direction cells. The preferred head direction exhibits a uniform distribution (*P* = 0.17, Rayleigh’s test).

To measure the sharpness of the head directional tuning curve, we quantified the average full width at half maximum (FWHM) of somatosensory head direction cells. The average FWHM of the angular distributions was 77.58 ± 4.36 degrees (mean ± s.e.m.) (Fig. 2d). The distribution of head direction preferences was uniformly distributed across 360° (Rayleigh test for uniformity: *P* = 0.17, Fig. 2e). We next evaluated the intra-trial directional stability by correlating the distributed firing rate across all directional bins between the first and second halves of individual recording trials. We found head direction related firing to be stable within sessions (Figs. S17a-c), with the average angular correlation coefficient for S1 head direction cells being 0.58 ± 0.02 (Fig. S17d). Like hippocampal place cells, head direction cells also anchored to salient visual cues ^52^. We further tested the specificity and stability of S1 head direction cells by rotating a salient visual cue 90° from its original position in the enclosure and back (Fig. S18, as in S1 place cells described above). As with S1 place cells, S1 head direction cells also rotated with the movement of the salient visual cue (Figs. S18a-c).

To measure the difference of the animal’s angular velocity within and outside preferred firing directions for those identified head direction cells in S1, we calculated the average angular velocity in the “preferred firing directions” and “non-preferred firing directions” for 80 identified head direction cells in S1 (Fig. S19a). We found that there was no significant difference between the average angular velocity in the “preferred firing directions” and “non-preferred firing directions” for head direction cells in S1 (52.97 ± 1.15 degrees/s vs 55.52 ± 1.34 degrees/s, *n* = 80, *P* = 0.12, two-tailed paired *t*-test, Fig. S19b).

Although the previous study had shown that food chasing had little effect on head directional firing in the rat anterior thalamic nuclei ^53^, we evaluated the firing patterns of head direction cells in the S1 without food pellet dispersion. We recorded the same head direction cell in two consecutive secessions with (bottom panels in Fig. S20a) and without (upper panels in Fig. S20a) food pellets, respectively. No significant differences were observed to suggest that food chasing did not modulate preferred firing angles in S1 head direction cells. Similar results were observed for place cells in the S1 without throwing food pellets within the running arena (upper panels in Fig. S20b).

### Border/Boundary vector coding in the somatosensory cortex

Boundary vector cells and border cells discharge exclusively whenever an animal is physically close, at a specific distance and direction, to one or several environmental boundaries, for example, the enclosure walls of the recording arena ^4, 5, 54^. S1 cells were defined as border cells if border scores (see Methods) were larger than the 99^th^ percentile of population shuffled scores (Fig. 3c). A total of 86/2025 (4.25%) S1 cells were classified as border cells, and this percentage was significantly higher than expected by random selection from the entire shuffled population (Fig. 3c; *Z* = 14.68, *P* < 0.001; binomial test with expected *P_0_* of 0.01 among large samples). Similar results were observed by performing cell-specific shuffling for those three representative somatosensory border cells (Fig. 3b). Most border cells (*n* = 58, 67.44%) fired along a single boundary while others were active along with two (*n* = 15, 17.44%), three (*n* = 6, 6.98%) or even four (*n* = 7, 8.14%) boundaries within the enclosure (Fig. 3a and Fig. S22). The average firing rate and the border score were 1.00 ± 0.09 Hz (Fig. S8c) and 0.66 ± 0.01, respectively. The histograms of peak-to-peak amplitude and peak-to-trough spike width were summarized in Fig. S8g. The number of S1 border cells seemed to be higher in layer V than in layer IV or layer VI (Fig. S3b and Fig. S3e). At a lower 95^th^ percentile criterion, 234/2025 (11.56%) somatosensory cells were classified as border cells.

**Fig. 3.**
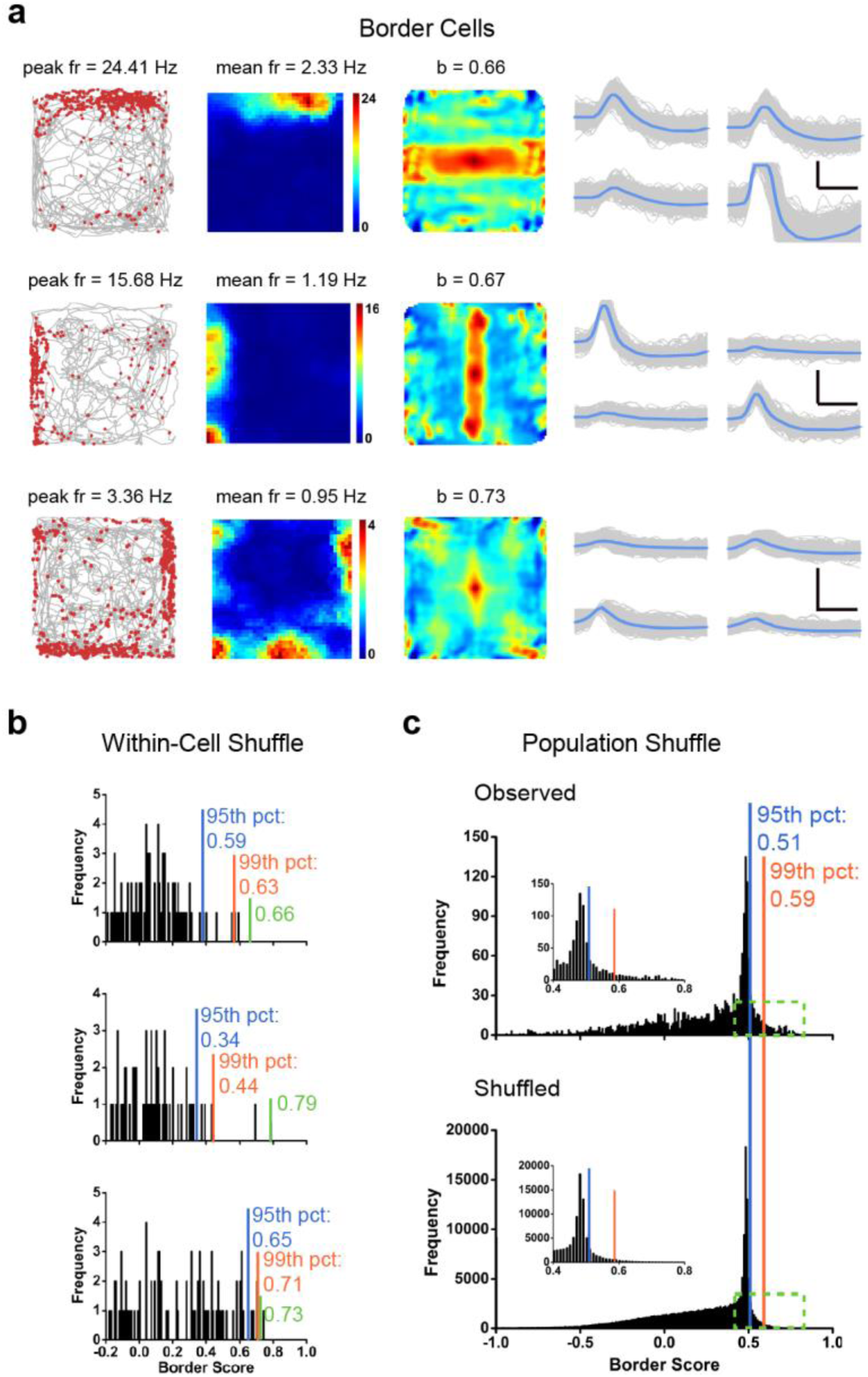
Border cells recorded from the somatosensory cortex. **(a)** Three representative examples of somatosensory border cells. Trajectory (grey line) with superimposed spike locations (red dots) (left column); heat maps of spatial firing rate (middle column) and autocorrelation diagrams (right column). Firing rate is color-coded with dark blue indicating minimal firing rate and dark red indicating maximal firing rate. The scale of the autocorrelation maps is twice that of the spatial firing rate maps. Peak firing rate (fr), mean firing rate (fr) and border score (b) for each representative border cell are labelled at the top of the panels. Spike waveforms on four electrodes are shown on the right column. Scale bar, 150 µV, 300 µs. (**b)** Distribution of within-cell shuffled border score for three representative somatosensory border cells. The orange and blue stippled lines mark the 99^th^ and the 95^th^ percentile significance level of each randomly shuffled distribution, respectively. The green line indicates the observed border score. (**c**) Distribution of border scores for pooled somatosensory cells. The top panel shows the distribution for observed values. The bottom panel shows the distribution for randomly shuffled versions from the same sample. The orange and blue stippled lines mark the 99^th^ and the 95^th^ percentile significance level of each randomly shuffled distribution, respectively. A zoomed panel shows the magnification of the specified area marked by the green dashed rectangle.

Qualitatively, S1 border cells maintained their border-specific firing fields between circular and square enclosures (Figs. S23a-c), comparable to boundary vector cells and border cells from the presubiculum, parasubiculum and the medial entorhinal cortex (MEC) ^4, 5, 17, 54^. S1 border cells also responded to a new wall insert to a familiar enclosure (Figs. S24a-c), as described in the MEC ^5, 54^. As observed in the MEC and the subiculum ^4, 5^, removal of walls from the enclosure on an elevated platform preserved similar “border” specificity to that in the walled arena ^54^. These results confirmed that somatosensory border cells continued to code for geometric boundaries instead of physical borders in the unwalled environments (Figs. S25a-c). Interestingly, we also detected a certain number of boundary vector cells ^4, 54^ which discharged further away from the edge of the enclosure (Fig. S26). Further quantitative characterization of somatosensory boundary vector cells combined with the previously described “perimeter” coding might reveal how somatosensory boundary vector cell closely resemble their counterparts in the subiculum ^1, 4, 54^.

### Grid cells in the somatosensory cortex

Hexagonal firing patterns identified in the MEC ^6^ can also be observed in the somatosensory cortex. To quantify the regularity of hexagonal firing patterns of the somatosensory cells (Fig. 4a and Fig. S28), we calculated grid scores based on rotated autocorrelograms of rate maps. A cell was categorized as a grid cell if its grid score was higher than the 99^th^ percentile of the shuffled data. 72/2025 cells (3.55%) met this criterion (Fig. 4d), and this percentage was significantly higher than expected by random selection from the entire shuffled population (Fig. 4d; *Z* = 12.90, *P* < 0.001; binomial test with expected *P_0_* of 0.01 among large samples). Similar results were observed by performing within-cell shuffling (Fig. 4b). To eliminate the possible influence of inhomogeneous sampling on the multiple firing fields of somatosensory grid cells and verify S1 grid cells through spike shuffling, we performed additional “field shuffling” ^22^ and found that these exampled grid cells also passed the gridness threshold for the “field shuffling’ procedure (Fig. 4c). The average firing rate and the grid score were 1.18 ± 0.12 Hz (Fig. S8d) and 0.55 ± 0.02 (Fig. S4d), respectively. The histograms of peak-to-peak amplitude and peak-to-trough spike width were presented in Fig. S8h.

**Fig. 4.**
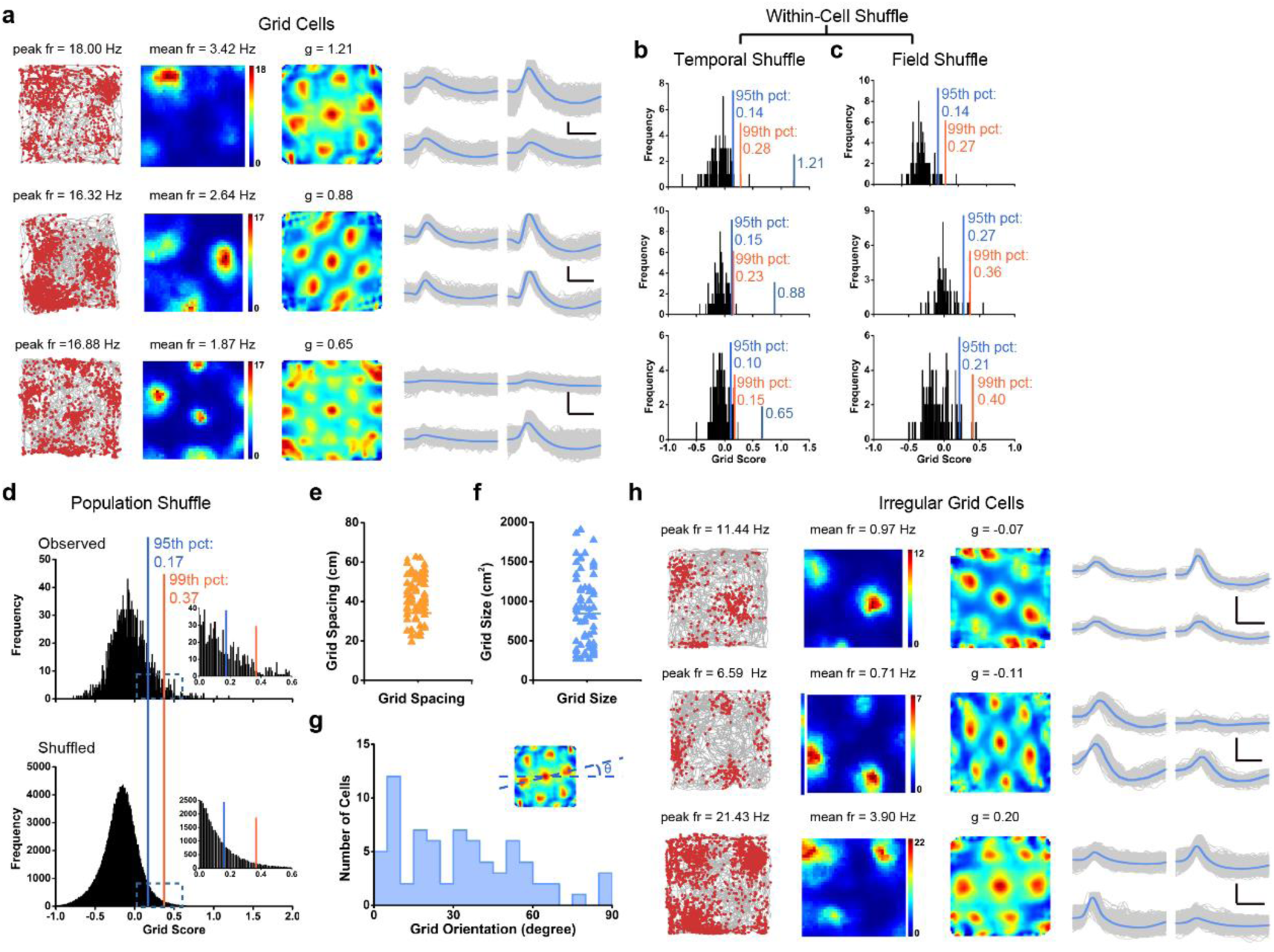
Grid cells in the somatosensory cortex. **(a)** Three representative examples of somatosensory grid cells. Trajectory (grey line) with superimposed spike locations (red dots) (left column); heat maps of spatial firing rate (middle column) and autocorrelation diagrams (right column). Firing rate is color-coded with dark blue indicating minimal firing rate and dark red indicating maximal firing rate. The scale of the autocorrelation maps is twice that of the spatial firing rate maps. Peak firing rate (fr), mean firing rate (fr) and grid score (g) for each representative grid cell are labelled at the top of the panels. A crystal-like hexagonal firing pattern was observed. Spike waveforms on four electrodes are shown on the right column. Scale bar, 100 µV, 300 µs. (**b)** Distribution of within-cell shuffled grid score for three representative somatosensory grid cells. The orange and blue stippled lines mark the 99^th^ and the 95^th^ percentile significance level of each randomly shuffled distribution, respectively. The cyan line indicates the observed grid score. (**c)** The same as (**b**) but for field shuffle. (**d**) Distribution of grid scores for somatosensory cells. The top panel shows the distribution for observed values. The bottom panel shows the distribution for randomly shuffled versions from the same sample. The orange and blue stippled lines mark the 99^th^ and the 95^th^ percentile significance level of each randomly shuffled distribution, respectively. A zoomed panel shows the magnification of the specified area marked by the cyan dashed rectangle. (**e-f**) The raster plots show the distribution of grid spacing and grid size of the classified somatosensory grid cells. (**g**) The histogram of grid orientation from categorized somatosensory grid cells. (**h**) Three representative examples of irregular somatosensory grid cells. Spike waveforms on four electrodes are shown on the right column. Scale bar, 150 µV, 300 µs.

Grid spacing in somatosensory grid cells (Fig. 4e) was 42.72 ± 1.30 cm, and the average grid size was 842.44 ± 54.22 cm^2^ (Fig. 4f). The fluctuation in somatosensory grid spacings is not a result of interindividual differences across different animals (Fig. S4d). Moreover, the orientation of the somatosensory grid cells varied and was on the average found to be 32.63 ± 2.64 degrees (Fig. 4g). Again, with a lower threshold at the 95^th^ percentile, 218/2025 S1 neurons (10.37%) were classified as grid cells, with the grid score threshold of 0.17 (blue line in Fig. 4d). When recorded in a larger environment (1.5 m x 1.5 m box), the discharging patterns of somatosensory grid cells showed similar multiple regular triangular structures in a larger enclosure to those within the smaller enclosure (1.0 m x 1.0 m) with a high grid score (Figs. S29a-c), confirming that those identified grid cells were not false-positive grid cells due to accidental triangular node structures. It appeared that somatosensory grid cells were less prevalent and a little noisier than place cells, head direction cells and grid cells identified in the S1. It remains to be determined whether multisensory inputs and intrinsic oscillatory dynamics within the somatosensory cortex might generate different rate-coded grid computations from those within the hippocampal-entorhinal cortex. Taken together, the local configuration of grid cells in the somatosensory cortex exhibited similar features as those in MEC. Furthermore, we detected irregular grid cells with negative grid scores in the S1 along with regular grid cells (Fig. 4h and Fig. S31).

### Conjunctive cells in the somatosensory cortex

To establish the existence of S1 neurons coding more than one spatial correlate, we assessed the strength of head direction tuning for all somatosensory place cells, border cells and grid cells ^7^. A certain number of somatosensory neurons were found to be conjunctive cells (Fig. 5 and Fig. S3a). Specifically, 25/195 place cells (Fig. 5a), 11/86 border cells (Fig. 5b) and 5/72 grid cells (Fig. 5c) were significantly tuned by head direction. Interestingly, we also detected irregular hexagonal grid cells modulated with head direction (Fig. 5d).

**Fig. 5.**
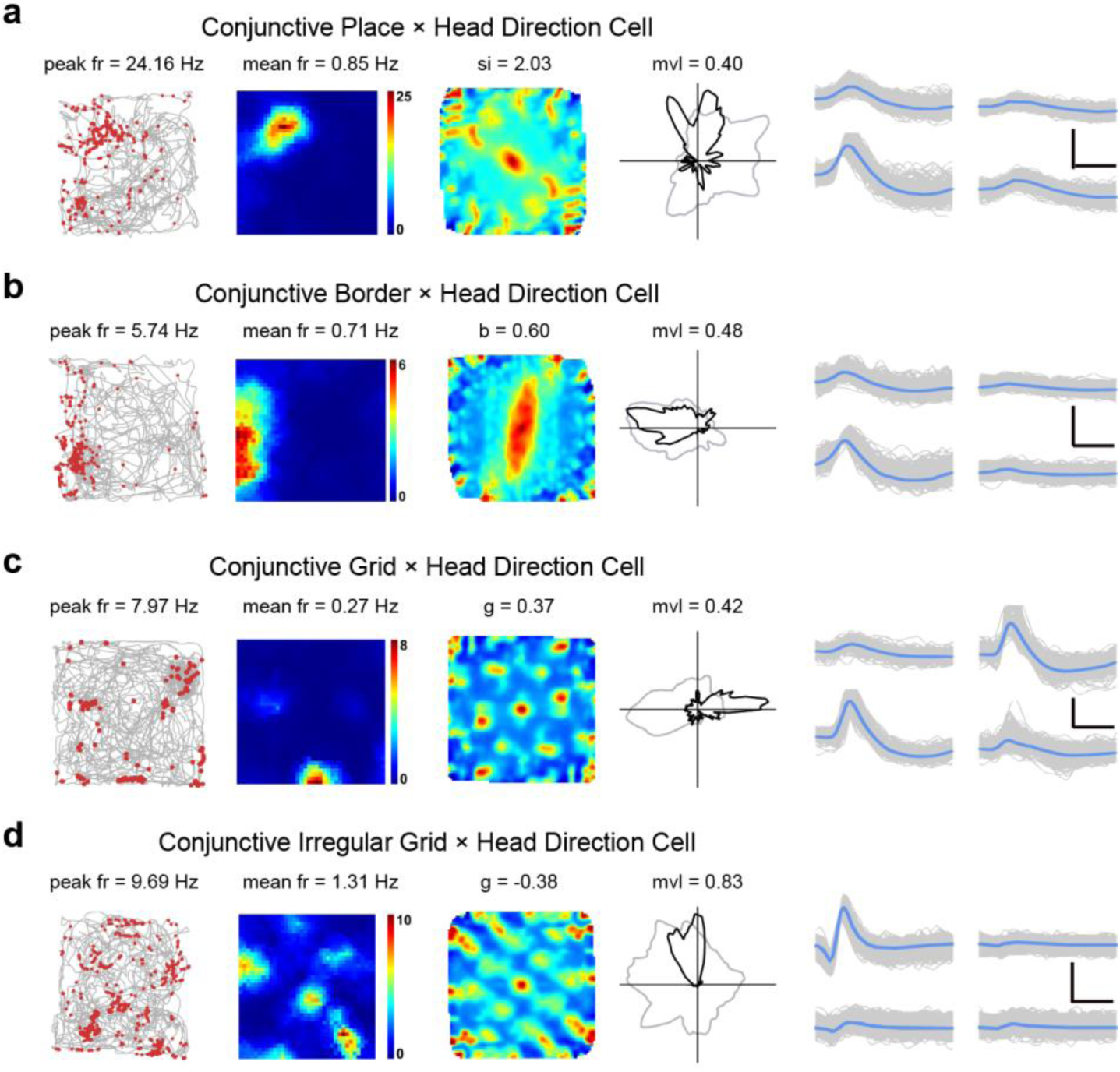
Four different types of conjunctive cells in the somatosensory cortex. (**a**) A representative conjunctive place x head direction cell. (**b**) A representative conjunctive border x head direction cell. (**c**) A representative conjunctive grid x head direction cell. (**d**) A representative conjunctive irregular grid x head direction cell. Trajectory (grey line) with superimposed spike locations (red dots) (left column); heat maps of spatial firing rate (middle left column), autocorrelation diagrams (middle right column) and head direction tuning curves (black) plotted against dwell-time polar plot (grey) (right column). Firing rate is color-coded with dark blue indicating minimal firing rate and dark red indicating maximal firing rate. The scale of the autocorrelation maps is twice that of the spatial firing rate maps. Peak firing rate (fr), mean firing rate (fr), spatial information (si), border score (b), grid score (g) and mean vector length (mvl) for each representative cell are labelled at the top of the panels. Spike waveforms on four electrodes are shown on the right column. Scale bar, 150 µV, 300 µs.

The degree of directionality in conjunctive cells showed no obvious difference from that of pure head direction cells (Watson’s U^2^ test, *P* = 0.09). These results suggested that the somatosensory cortex carried similar conjunctive spatial x directional, border x directional and gird x directional signal to those identified previously in the rat subiculum ^45^ and the rat MEC ^5, 7^.

### Impact of the whisker trimming on the somatosensory spatial activities

Since rodents use their whiskers to actively explore the external environment for whisker-based navigation ^55, 56^, we implanted two additional rats (Fig. S1b) and recorded neuronal activity before and after repeated whisker trimming (Fig. S32a and Fig. S32c) to gauge the involvement of the vibrissae. Two consecutive recording sessions from the same rat before and right after whisker trimming showed that the distribution of clusters and waveforms in the S1 was qualitatively similar (Fig. S32b and Fig. S32d).

To further quantify the change of well-isolated spike clusters before and after whisker trimming, we calculated the isolation distance and L-ratio ^42^. The median isolation distances were 122.35 and 132.9 before and after whisker trimming, respectively. There was no significant difference in cluster separation and L-ratio in the S1 before and right after whisker trimming (Fig. S32e and Fig. S32f, *n* = 78, *P* = 0.73 and 0.76, respectively, two-tailed paired *t*-test). The average drift of center of mass for the total cell sample before and after whisker trimming was 0.13 ± 0.01, indicating there was minimal change in spike clusters before and after whisker trimming. Besides, both peak and average firing rates of the same neurons before and after whisker trimming did not change significantly (Fig. S32g and Fig. S32h, *n* = 78, *P* = 0.75 and 0.47, respectively, two-tailed paired *t*-test).

To further assess whether whisker trimming affected the S1 spatial firing properties, we were able to continuously record the same S1 border cell before and after whisker trimming. The spatial firing patterns of the border cell persisted and responded to a new wall inserted into the square enclosure (Fig. S33a and Fig. S33b). Overall, both the spatial responses and firing properties in the S1 were not significantly affected by the whisker trimming.

When we pooled and analyzed a total of 461 cells recorded after whisker trimming, we were still able to identify all different spatial cell types including 43 place cells (Fig. 6a), 15 head direction cells (Fig. 6b), 29 border cells (Fig. 6c) and 17 grid cells (Fig. 6d). The percentage of all recorded spatially tuned cell types after whisker trimming were similar to those recorded from rats without whisker trimming. Furthermore, the cutoffs for defining place cells (Fig. 6e), head direction cells (Fig. 6f), border cells (Fig. 6g) and grid cells (Fig. 6h) after whisker trimming highly resembled those for different somatosensory spatial cell types without whisker trimming. Taken together, the persistence of recorded somatosensory spatial cells after whisker trimming demonstrated that our identified cells were not significantly influenced by the whisker trimming.

**Fig. 6.**
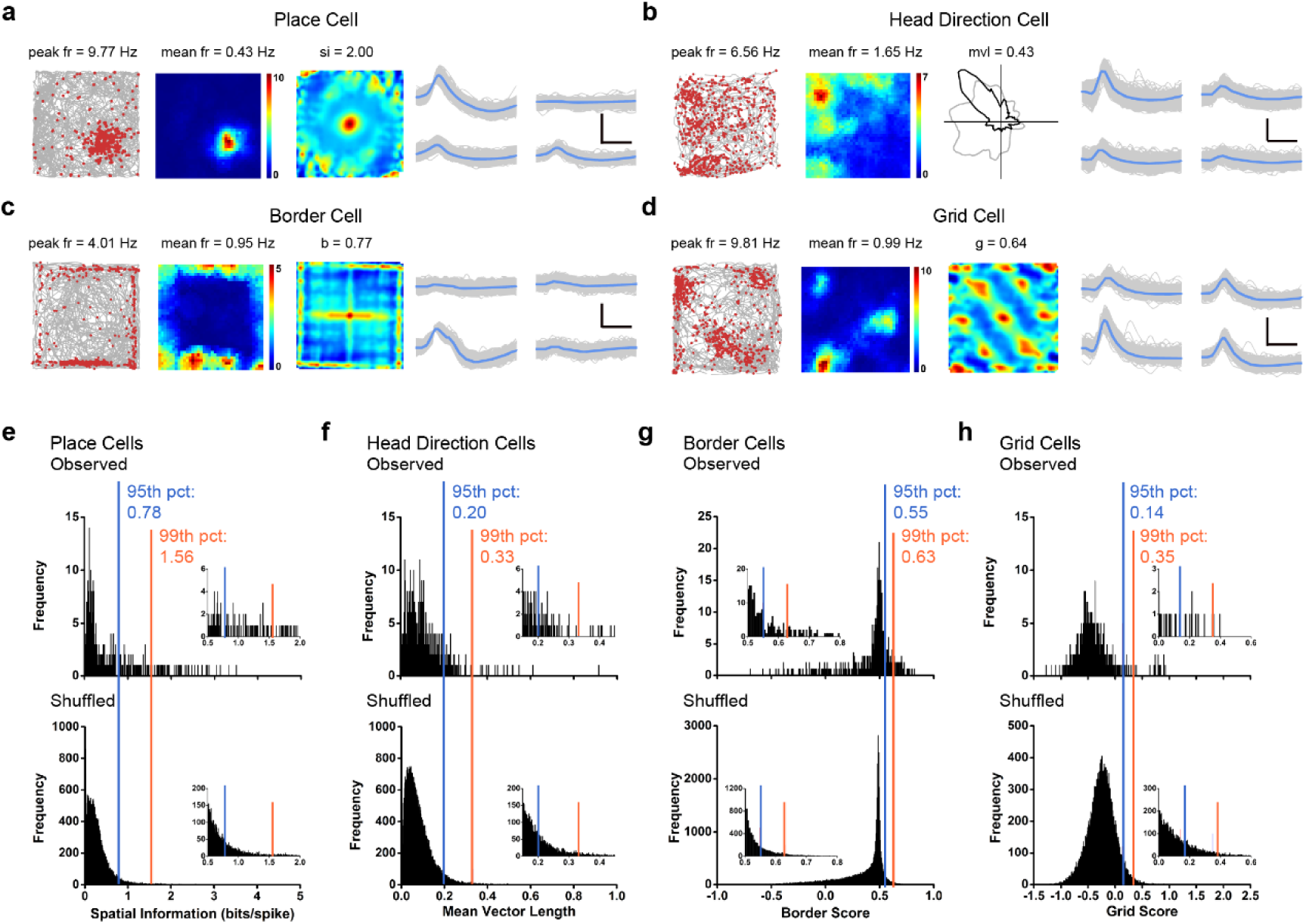
Somatosensory spatial response after the whisker trimming. **(a**-**d)** Four representative examples of somatosensory place cell (**a**), head direction cell (**b**), border cell (**c**) and grid cell (**d**) after whisker trimming. Trajectory (grey line) with superimposed spike locations (red dots) (left column); spatial firing rate maps (middle left column), autocorrelation diagrams (middle right column) and head direction tuning curves (black) plotted against dwell-time polar plot (grey) (right column). Firing rate is color-coded with dark blue indicating minimal firing rate and dark red indicating maximal firing rate. The scale of the autocorrelation maps is twice that of the spatial firing rate maps. Peak firing rate (fr), mean firing rate (fr), spatial information (si), mean vector length (mvl), border score (b) or grid score (g) and angular peak rate for each representative cell are labelled at the top of the panels. Spike waveforms on four electrodes are shown on the right column. Scale bar, 150 µV, 300 µs. (**e-h)** Distribution of spatial information (**e**), mean vector length (**f**), border score (**g**) or grid score (**h**) for somatosensory cells after whisker trimming. The top panel shows the distribution for observed values. The bottom panel shows the distribution for randomly shuffled data from the same sample. The orange and blue stippled lines mark the 99^th^ and the 95^th^ percentile significance level of each randomly shuffled distribution, respectively.

## Discussion

This study provides the first demonstration and characterization of spatially tuned neuronal discharges in the primary somatosensory cortex. We have shown place, grid, boundary vector/border, head direction and conjunctive cell activities, as reported in parahippocampal and associated areas ^39^, to co-exist in the single-domain primary somatosensory cortex. These spatially tuned activities encoded within the newly identified somatosensory spatial navigation system appear to be specific, stable, anchored to external visual cues and robust against non-spatial environmental perturbations.

Based on the assumption that proprioception makes a significant contribution to spatial representation in the parahippocampal cortices ^39, 40^, we predicted spatially selective activities should exist in the somatosensory cortex. To our surprise, we were able to record the full complement of spatially selective cells (place, grid, boundary vector/border, head direction and conjunctive cells) in the primary somatosensory cortex. Given the paucity of direct projections from the parahippocampal areas to the S1 region ^57, 58^, we assume somatosensory spatial activities do not arise from the hippocampal-entorhinal circuit. However, we cannot completely eliminate the possibility that the somatosensory spatial positioning system may be at least partially dependent on the hippocampal-entorhinal system. Whether the newly found somatosensory spatial navigation system is independent of, parallel with or convergent onto the classical one within the hippocampal-entorhinal microcircuit remains an interesting question and needs to be addressed in future studies ^1^. The availability of single-cell transcriptomics and connectomics for the somatosensory cortex might help to unravel the interaction or interdependence between these two genetically pre-configured spatial maps ^59, 60^. Since there is very little overlap between septal projections to the hippocampal-entorhinal areas ^61^, and that septal inactivation severely attenuates place and grid cell activities in both regions ^62–64^, how S1 responds to septal inactivation may provide the necessary insight for hippocampal-entorhinal dependency.

The laminar organization of the S1 in relation to connectivity and function has been well-characterized, largely arising from studies on the vibrissae barrel fields ^65, 66^. There is insufficient information on the specific connectivity patterns of S1 in the rat, but mouse data suggest similar cortico-cortical, thalamic-cortical, cortical-thalamic, cortico-striatal and corticofugal connection patterns between S1 and other parts of the S1 ^67^. Our current study is roughly limited to sampling layers IV through VI. There is a pattern for head direction cells to be absent in layer VI and place cells to predominate in deeper layers (**Figs. S3e-h**). From such distribution of spatial cell types across cortical depth, we could infer that head direction information is not present in cortico-thalamic communications, given the lack of head direction cells at depths corresponding to layer VI in the somatosensory cortex. There also appears to be a gap in the most superficial aspect of layer V (Va) where all spatial cell types appear to be under-represented. Interestingly, layer Va projections are mostly recurrent ^68^ and appear to serve as a trans-columnar and trans-laminar integration hub within the S1 ^69–71^. Layer Va may serve as an area that integrates spatial and proprioceptive information in the S1, hence salient spatial activity is not readily observable in this layer. If true, this would also suggest that the major extra-cortical output from S1 combines, but do not contain conventional spatial information as characterized here. A full sampling of cortical layers in S1 using high-density linear probes ^72^ is necessary to address how spatial information is generated, integrated and distributed elsewhere (e.g. thalamic, striatal and brainstem targets) in the brain.

As mentioned above, there is a lack of direct connections between S1 and areas that are traditionally associated with spatial processing. Where do the spatial signals come from? Are they more universal in cortical processing than previously thought? What is the functional significance of spatial activities in the S1, or S1 in general? While extensive intra and inter columnar/laminar exist in the S1 and may support *de novo* generation of place cells (from integrating local grid cell activities) and conjunctive cells, it is unlikely the full complement of known spatial representations are generated independently within the S1. We have demonstrated that S1 place fields and preferred head direction rotate with the varied positioning of salient visual cue as cells found in the hippocampal-entorhinal circuit. This observation suggests the spatial activities cannot solely be driven by proprioception arising from lemniscal/paralemniscal pathways. While a sparse, direct and functional connection exists between the S1 and the visual cortex ^73^, it is unlikely to account for visually anchored spatial tuning in the somatosensory cortex. Instead, S1 spatial activities are more likely to be an efference copy from elsewhere in the brain.

S1 functions and activities are intricately linked to motor structures, especially the motor cortex. Functional coupling between S1 spindle activity and spontaneous muscle twitches can be observed as early as two days after birth ^74^. It is known that both primary and secondary motor cortex are reciprocally connected with S1 ^75, 76^, and the preparation, execution and dynamic changes in movements can all be predicted from M1/S1 activities ^77^. In fact, S1 has recently been shown to have an active role in directly controlling motor output ^78^. Position-dependent neuronal discharges (i.e. place field-like representations) relating to hand movement through space are well-established in the primate motor cortex ^79^. In fact, considering movement of the hand or forelimb in general: speed ^79, 80^, (hand) direction ^79^ and conjunctive ^79, 80^ coding in the motor cortices can be seen as homologous to their spatial navigation counterparts in the limbic system. Essentially, it appears the motor cortex contains spatially tuned activities for specific body parts that are analogous to limbic activities that appear to represent whole body/head movement. Therefore, whole body/head and limb movements through space may require similar neurocomputation, of which is also present in the S1 in addition to motor and limbic cortices. This proposal is also consistent with the idea that S1 simulates the body itself in addition to its representation, which requires spatial information ^37^. From this synthesis, we speculate that we would be able to record spatially tuned activities in the limb region of the motor cortex, as well as other sensorimotor cortices (e.g. the forelimb and trunk regions). Particularly, inactivation of the reciprocally connected motor cortex (i.e. limb region of the motor cortex) can test if S1 spatial activities are efference copy from the motor cortex. Recent reports of place cell-like activities in the primate sensorimotor cortices ^38^ during quasi-active movements through space support the notion that spatial representation of body movement may exist throughout the cortex.

Our discussion so far favors somatosensory spatial activity may be an efferent copy from the motor cortex. If true, then the question remains as to how an area such as the motor cortex without direct connectivity with spatially tuned limbic structures (like the S1) can acquire spatial activities. To the best of our knowledge, no grid- or boundary vector-/border-like spatial representation has been reported in the motor cortex in conventional reaching studies, although this discrepancy may be related to that motor tasks in such studies are often limited to a single defined trajectory (insufficient spatial coverage to detect grid cells) in open task space (without physical borders). Likewise, the “body simulation” model ^37^ predicts the existence of body part location information (which we interpret to be place cell-like representations) to exist in layer IV, but our data indicate the full complement of spatial activities can be detected from at least layers IV through VI. Our data is not fully compatible with currently available experimental and theoretical work in explaining why all currently recognized spatial cell activities can be found in S1 and how they are generated. Regardless, we propose it is likely spatial information is integral to the “body simulation” function in S1 as put forward by Brecht ^37^. High-resolution kinematic studies ^81^ of body part (e.g. limb) movements through space would be required to test if spatial activities in the S1 do in fact represent body parts in space.

In this report, we have demonstrated place, grid, boundary vector/border, head direction and conjunctive cell activities in the primary somatosensory cortex. These activities are comparable to their counterparts in the parahippocampal cortices and are stable, specific, robust to non-spatial manipulations and anchored to salient spatial cues. We propose spatial tuning in the primary somatosensory cortex may be crucial for body representation/simulation in space, and the newly identified somatosensory spatial representation system may generalize to other sensorimotor cortices pertinent to behavioral coordination.

## Materials and Methods

### Subjects

Ten male Long-Evans rats (2-4 months old, 250-450 grams at the time of the surgery) were used for this study. All animals were singly housed in transparent cages (35 cm x 45 cm x 45 cm, W x L x H) and maintained on a 12-hour reversed light-dark cycle (lights on at 9 p.m. and off at 9 a.m.). Experiments were performed during the dark phase. Rats were maintained in a vivarium with controlled temperature (19-22°C), humidity (55-65%). and were kept at about 85-90% of free-feeding body weight. Food restriction was imposed 8-24 hours before each training and recording trial. Water was available *ad libitum*. All animal experiments were performed in accordance with the National Animal Welfare Act under a protocol approved by the Animal Care and Use Committee from both Army Medical University and Xinqiao Hospital.

### Surgery and tetrode placement

Rats were anesthetized with isoflurane. Microdrives loaded with four tetrodes were implanted to target the hindlimb (S1HL), forelimb (S1FL) and shoulder (S1Sh) regions of the primary somatosensory cortex (anterior-posterior (AP): 0.2-2.2 mm posterior to bregma; medial-lateral (ML): 2.2-3.4 mm lateral to midline, dorsal-ventral (DV): 0.4/0.6-3 mm below the dura.), secured with dental cement with 8-10 anchor screws. A screw served as the ground electrode. Tetrodes were assembled with four 17 µm Platinum/Iridium wires (#100167, California Fine Wire Company). Tetrodes had impedances between 150 and 300 kΩ at 1 kHz through electroplating (nanoZ; White Matter LLC). 8-10 jeweler screws were attached into the rat skull, and individual microdrives were anchored to screws with several rounds of application of the dental cement.

### Training and data collection

Behavioral training, tetrode turning and data recording started one-week post-surgery. Rats were trained to run around in a 1 m x 1 m square box with a white cue card (297 mm x 210 mm) mounted on one side of the wall. Food pellets were scattered into the arena intermittently to encourage exploration.

Each recording session lasted between 10 and 30 min to facilitate full coverage of the testing arena. Tetrodes were lowered in steps of 25 or 50 µm daily until well-separated single units can be identified. Data were acquired by an Axona system (Axona Ltd., St. Albans, U.K.) at 48 kHz, band-passed between .8-6.7 kHz and a gain of x5-18k. Spikes were digitized with 50 8-bit sample windows. Local field potentials were recorded from one of the electrodes with a low-pass filter (500 Hz).

### Spike sorting, cell classification and rate map

Spike sorting was manually performed offline with TINT (Axona Ltd, St. Albans, U.K.), and the clustering was primarily based on features of the spike waveform (peak-to-trough amplitude and spike width), together with additional autocorrelations and cross-correlations ^44, 82^. During our manual cluster cutting, we always counted neurons with similar or identical waveform shapes only once whenever similar or identical individual neurons were recorded and tracked across two consecutive recording sessions. To confirm the quality of cluster separation, we calculated L-ratio as well as isolation distance between clusters.

Two small light-emitting diodes (LEDs) were mounted to the headstage to track the rats’ speed, position and orientation via an overhead video camera. Only spikes with instantaneous running speeds > 2.5 cm/s were chosen for further analysis in order to exclude confounding behaviors such as immobility, grooming and rearing.

To classify firing fields and firing rate distributions, the position data were divided into 2.5-cm x 2.5-cm bins, and the path was smoothed with a 21-sample boxcar window filter (400 ms; 10 samples on each side) ^7, 17, 43^. Cells with > 100 spikes per session and with a coverage of >80% were included for further analyses. Maps for spike numbers and spike times were smoothed with a quasi-Gaussian kernel over the neighboring 5 x 5 bins ^7, 17, 43^. Spatial firing rates were calculated by dividing the smoothed map of spike numbers with spike times. The peak firing rate was defined as the highest rate in the corresponding bin in the spatial firing rate map. Mean firing rates were calculated from the whole session data.

### Analysis of place cells

Spatial information is a quantification of the extent to which a neuron’s firing pattern can predict the position of freely moving animals and is expressed in the unit of bits per spike. The spatial information was calculated as:

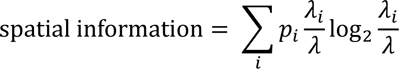

where *λ*_*i*_ is the mean firing rate of the cell in the *i-*th bin, *λ* is the overall mean firing rate of the cell in the trial, and *p*_*i*_ is the probability for the animal being at the location of the *i-*th bin.

Adaptive smoothing ^44^ was applied to optimize the trade-off between spatial resolution and sampling error before the calculation of spatial information. The data were first divided into 2.5-cm x 2.5-cm bins, and then the firing rate within each bin was calculated by expanding a circle centered on the bin until

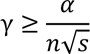

where γ is the circle’s radius in bins, *n* is the number of occupancy of samples within the circle, *s* is the total number of spikes fired within the circle, and α is a constant parameter set to 10,000. With a position sampling frequency at 50 Hz, the firing rate assigned to that bin was then set to 50 · *s*/*n*.

A place cell was classified as a cell with the spatial information above chance level, which was computed by a random permutation process using all recorded cells. For each round of the shuffling process, the entire sequence of spike trains from each cell was time-shifted along the animal’s trajectory by a random period between 20 s and the trial duration minus 20 s, with the end wrapped to the beginning of the trial. A spatial firing rate map was then constructed, and spatial information was calculated. This shuffling process was repeated 100 times for each cell, generating a total of 202,500 permutations for the 2025 somatosensory neurons. This shuffling procedure preserved the temporal firing characteristics in the unshuffled data while disrupting the spatial structure at the same time.

Spatial information score was then measured for each shuffled rate map. The distribution of spatial information values across all 100 permutations of all cells was computed and finally, the 99^th^ percentile of the significant level was determined. The threshold values for categorizing cells into place cells were defined as the spatial information scores above the 99^th^ percentile of the distribution from shuffled populations. In addition to the population shuffling, the within-cell shuffling was also performed within the entire sequence of spike trains from each individual neuron, and the shuffling process was also repeated 100 times for each single cell.

Spatial sparsity was used to measure how compact and selective the place field of each place cell is relative to the recording enclosure. The spatial sparsity was calculated using the formula as follows ^44^:

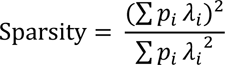

Where *p*_*i*_ is the occupancy probability for the animal being at the location of the *i*-th bin in the map and *λ*_*i*_ is the mean firing rate of the cell in bin *i*.

Spatial coherence was estimated by calculating the mean correlation between the firing rate of each bin in the map and the aggregate firing rate of the eight nearest bins ^34^. We used unsmoothed firing rate maps for computing the spatial coherence.

The spatial correlation across trials from the same recording arena was computed for each cell by correlating the firing rates in corresponding paired bins of two smoothed rate maps. The spatial stability within trials was estimated by calculating spatial (2D) correlations between firing rate maps generated from the first and second halves of the same trial. Place cells with spatial stability lower than 0.3 were excluded for further analysis.

### Analysis of grid cells

Spatial autocorrelation was calculated with smoothed rate maps ^7, 17, 43^. Autocorrelograms were derived from Pearson’s product-moment correlation coefficient correcting for edge effects and unvisited locations.

With *λ*(*x*, *y*) representing the average firing rate of a cell at coordinate (*x*, *y*), the autocorrelation between the spatial firing field itself and the spatial firing field with lags of *τ*_*x*_ and *τ*_*y*_ was calculated as:

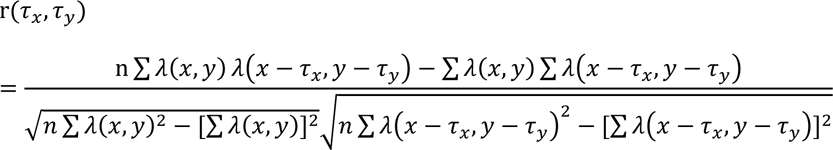

where the summation is over whole n pixels in *λ*(*x*, *y*) for which firing rate was calculated for both *λ*(*x*, *y*) and *λ*(*x* − *τ*_*x*_, *y* − *τ*_*y*_). Autocorrelations were not calculated for spatial lags of *τ*_*x*_, *τ*_*y*_ where *n* < 20.

The degree of spatial regularity (“gridness” or “grid score”) was calculated for each unit by using a circular sample centered on the central peak of the autocorrelogram but excluding the central peak itself, and by comparing rotated versions of this circular sample ^5, 7^. The Pearson’s correlations between this circular sample and its rotated versions were calculated, with the angles of rotation of 60° and 120° in the first group, and 30°, 90° and 150° in the second group. Gridness or the neuron’s grid score was defined as the minimal difference between any of the coefficients in the first group and any of the coefficients in the second group. Shuffling was performed in the same procedure used for defining place cells. Grid cells were categorized as cells with the rotational symmetry-based grid scores exceeding the 99^th^ percentile of the distribution of grid scores for shuffled data from the entire population of identified somatosensory cells.

Grid spacing was computed as the median distance from the grid center to the closest peak among six neighboring firing fields in the autocorrelogram of the spatial firing map. Since such analysis is sensitive to noise in the grid autocorrelogram, grid spacing was computed only when the median distance to the six neighboring peaks for the analyzed cell was comparable to the radius of the circle centered on the gird autocorrelogram with the highest grid score. The radius of this circle around the center of the autocorrelogram was also referred to as the grid field size.

Grid orientation was calculated by first computing vectors from the center of the autocorrelogram to each of the three adjacent peaks among six neighboring firing fields in the autocorrelogram of the spatial firing map in the counterclockwise direction, beginning with a camera-based reference line of zero degree. The angle between the minimal orientation of those three vectors and the camera-based reference line was defined as the grid orientation.

### Analysis of head direction cells

The rat’s head direction was estimated by the relative position of the LEDs differentiated through their sizes ^7, 17, 43^. The directional tuning curve for each recorded cell was drawn by plotting the firing rate as a function of the rat’s head angle, which is divided into bins of 3 degrees and then smoothed with a 14.5-degree mean window filter (2 bins on each side). To avoid bias, data were only used if all head angle bins contain data.

The strength of directionality was calculated by computing the mean vector length from circular distributed firing rates. The chance values were determined by a shuffling process simulated in the same way as for place cells, with the entire sequence of spike trains time-shifted between 20 s and the whole trail length minus 20 s along the animal’s trajectory. Cells were defined as head direction cells if the mean vector lengths of the recorded cells were larger than the 99^th^ percentile of the mean vector lengths in the shuffled distribution. Angular stability was computed by calculating the correlation of firing rates across directional bins generated from the first and second halves of the same trial. Head direction cells with angular stability lower than 0.3 were excluded for further analysis.

### Analysis of border cells

Border or boundary vector cells were identified by calculating, for each recorded cell, the difference between the maximal length of any of the four walls touching on any single spatial firing field of the cell and the average distance of the firing field to the nearest wall, divided by the sum of those two values ^5, 17, 43^. Border scores ranged from −1 for cells with perfect central firing fields to +1 for cells with firing fields that exactly line up with at least one entire wall. Firing fields were defined as summation of neighboring pixels with total firing rates higher than 0.3 times the cell’s maximum firing rate that covered a total area of at least 200 cm^2^.

Border cell classification was verified in the same way as for place cells, head direction cells and grid cells. For each permutation trial, the whole sequence of spike trains was time-shifted along the animal’s trajectory by a random period between 20 s and 20 s less than the length of the entire trial, with the end wrapped to the start of the trial. A spatial firing rate map was then obtained, and a border score was estimated.

The distribution of border scores was calculated for the entire set of permutation trials from all recorded cells, and the 99^th^ percentile was then determined. Cells were defined as border cells if the border score from the observed data was higher than the 99^th^ percentile for border scores in the entire distribution generated from the permutated data.

### Whisker trimming

To test how the removal of another salient somatosensation mediate by vibrissae input impacts on FL/HL S1 spatial cells, we carefully trimmed all whiskers of each recoded rat bilaterally using blunt surgical scissors to within 1-2 mm above the skin surface just right before each recording session every day. Since bilateral whisker-trimmed rats did not exhibit abnormal behaviors, we continuously recorded somatosensory units from 2 whiskers-trimmed rats.

### Environmental manipulations

For visual landmark rotation, we first recorded neuronal activity in the standard session followed by a 90° cue-card rotation in the clockwise or counterclockwise direction. Then another standard session was performed with the cue-card rotated back to the original position. For recording in the elevated platform without walls, we first recorded the somatosensory spatial cells in the square box, followed by the recording in the elevated platform without walls. Finally, the animals were returned to the original square box for another recording session. For the recording of border cells in the presence of inserted wall, we first identified the somatosensory border cells in the square box. Then the recording session was followed by the insertion of a wall along the center of the external wall. Another recording session was performed after removing the inserted wall. For food/no food comparisons, we recorded the somatosensory spatial cells in two consecutive recording sessions without and with throwing food pellets into the running enclosure.

### Histology and reconstruction of recording positions

At the end of the experiment, rats were euthanized with an overdose of sodium pentobarbital and perfused transcardially with phosphate-buffered saline (PBS) followed by 4% paraformaldehyde (PFA). Brains were removed and stored in 4% PFA overnight. The brain was then placed in 10, 20 and 30% sucrose/PFA solution sequentially across 72 hours before sectioning using a cyrotome. Thirty-micron sections were obtained through the implant region. Sections were mounted on glass slides and stained with cresyl violet (Sigma-Aldrich). The final recording positions were determined from digitized images of the Nissl-stained sections. Positions of each individual recordings were estimated from the deepest tetrode track, notes on tetrode advancement with tissue shrinkage correction by dividing the distance between the brain surface and electrode tips by the last advanced depth of the recording electrodes. Electrode traces were confirmed to be located within the hindlimb region (S1HL) from eight implanted rats, the forelimb region (S1FL) from one rat and the shoulder region (S1Sh) from another rat but all electrode tracking paths from 10 implanted rats were verified to be away from the barrel field (S1BF) of the primary somatosensory cortex according to the Rat Brain Atlas ^83^.

## Acknowledgments

We would like to thank five anonymous reviewers for their insightful comments and constructive suggestions during the peer reviews. We are indebted to H. Wu, W. Tang, H. Chen, S. Lv, H. Yang and J. Ni for their encouragement and generous help. We would like to acknowledge J. Cai, B. Deng, L. Hu and S. Wang for their technical assistance. We would like to express our gratitude to Neil Burgess and Caswell Barry for sharing their Matlab codes with us. We would like to appreciate Calvin Young for his critical comments on the manuscript. X.L. is supported by the Chongqing Municipality postdoctoral fellowship (Grant# cstc2019jcyj-bshX0035). We are grateful for two medical research funds from Xinqiao Hospital (Grant# 2017A034 and Grant# 2019XQY16) and a startup fund from Army Medical University (Grant# 2017R028) to S.-J.Z. This work was also supported by the National Natural Science Foundation of China through the Project Grant NSFC-31872775 to S.-J.Z. Original results published in this manuscript were previously uploaded on the preprint server for biology (https://www.biorxiv.org/content/10.1101/473090v1) at *bioRxiv* on November 19, 2018 ^41^.

## Data Availability

Recording dataset will be prepared available in a forthcoming public domain, and inquiries into acquiring the recording dataset beforehand should be directed to the corresponding author.

## Author Contributions

S.-J.Z. conceived and designed the study. X.L. and S.-J.Z. performed the experiments, collected the data and performed the analyses. X.L. made the figures. S.-J.Z. wrote the manuscript.

## Additional Information

### Supplementary information

(supplementary Figs. S1-S33 and Table S1) accompanies this paper.

---

**Fig. S1.**
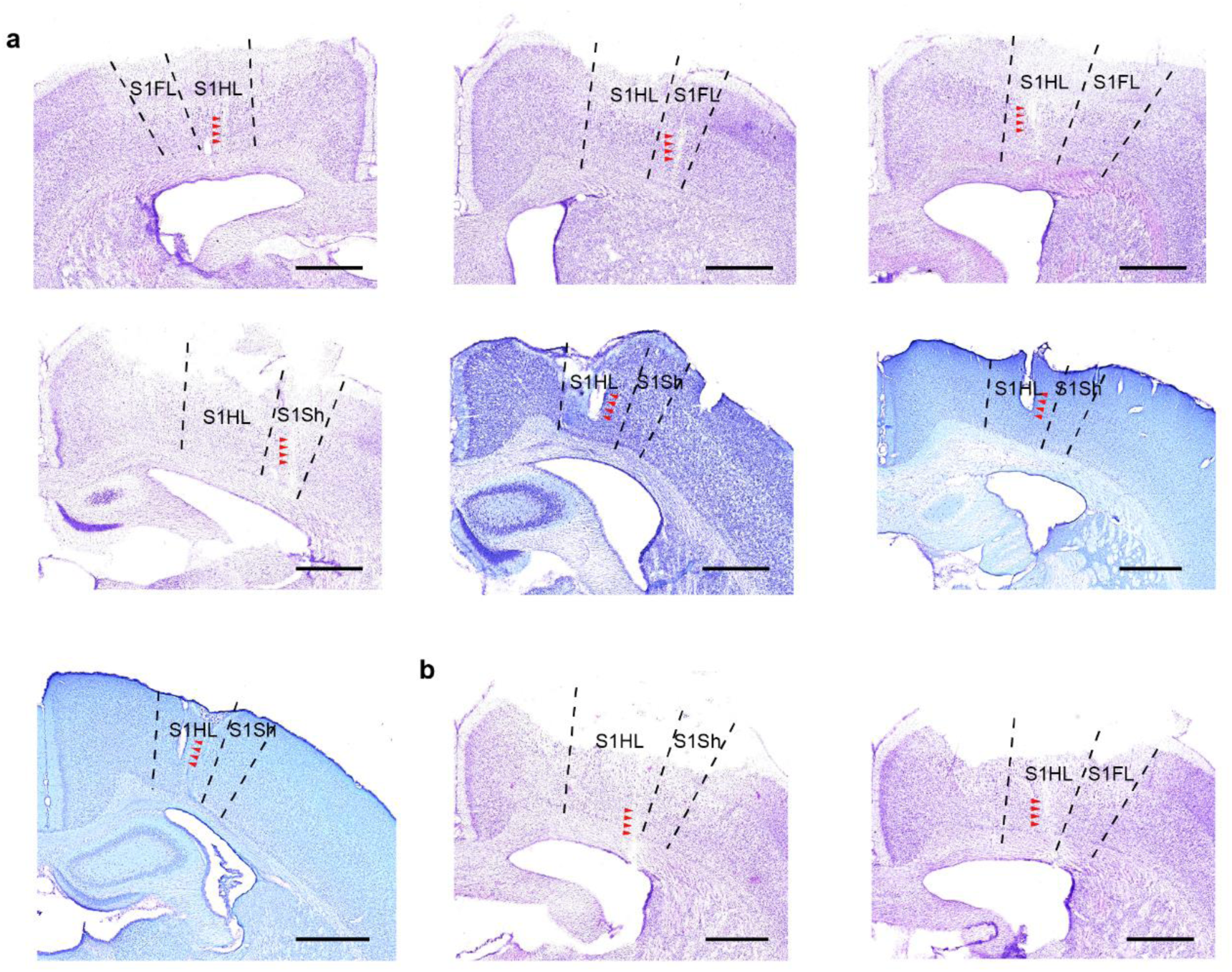
Electrode track and recording locations in the primary somatosensory cortex. (**a**) Cresyl violet-stained coronal brain sections show recording electrode tracks (arrowheads) and final recording positions in seven rats with tetrodes implanted in the rat primary somatosensory cortex. (**b**) Cresyl violet-stained coronal brain sections showing representative recording locations (arrowheads) for two additionally implanted rats in the whisker trimming experiment. Broken dashes depict the boundaries of the hindlimb region (S1HL), shoulder region (S1Sh) and forelimb (S1FL) region of the primary somatosensory cortex according to the rat brain atlas of Paxinos and Watson (2007). Scale bar, 1 mm.

**Fig. S2.**
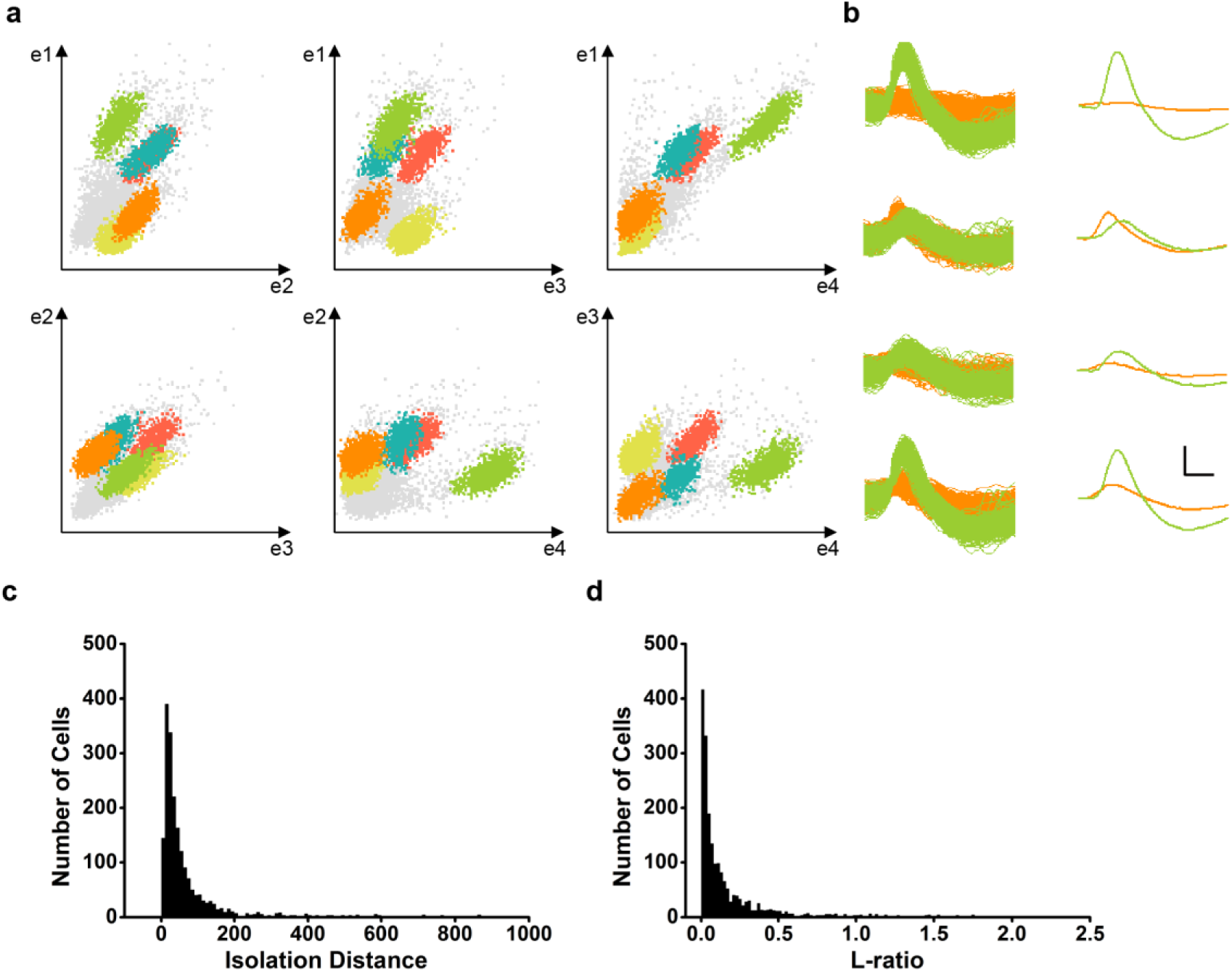
Cluster diagrams, waveforms and isolation quality of spike clusters recorded from the somatosensory cortex. (**a**) Scatterplots show the relationship between peak-to-trough amplitudes for all spikes for each wire (e1-e4) in a tetrode. (**b**) Overlaid and mean waveforms from two separated green and orange clusters in the scatterplots are shown for four electrodes from the same tetrode. Scale bar, 150 µV, 200 µs. (**c**) The distribution of isolation distance for identified somatosensory units. (**d**) Same as (**c**) for the L-ratio.

**Fig. S3.**
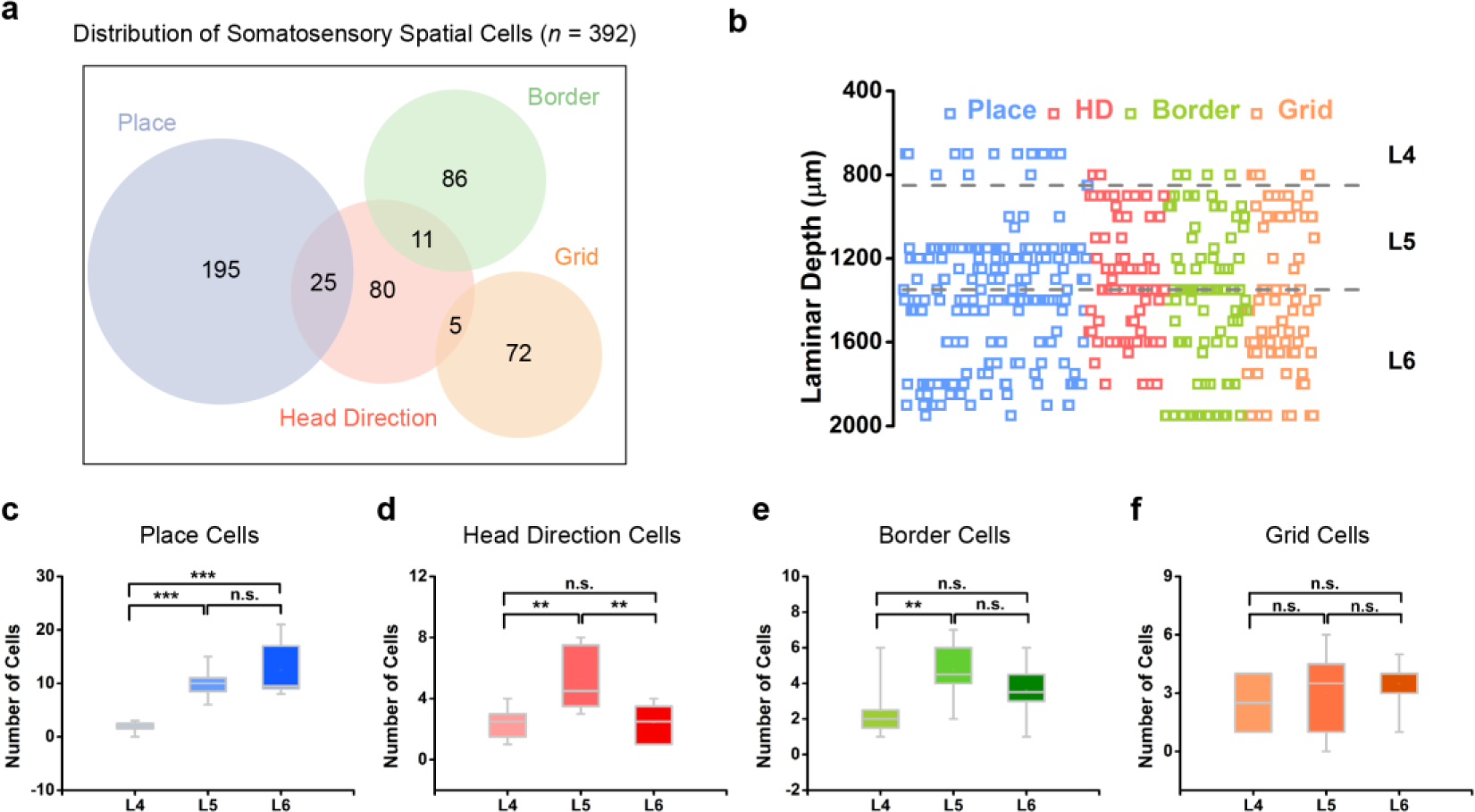
Layer distribution of four different somatosensory spatial cell types. (**a**) Venn diagram displaying the distribution of four functionally distinct somatosensory spatial cell types. (**b**) The approximate distributed laminar locations of all recorded functionally distinct somatosensory spatial cells. (**c-f**) The layer distribution of average number of identified somatosensory place cells (**c**), head direction cells (**d**), border cells (**e**) and grid cells (**f**) across different implanted rats.

**Fig. S4.**
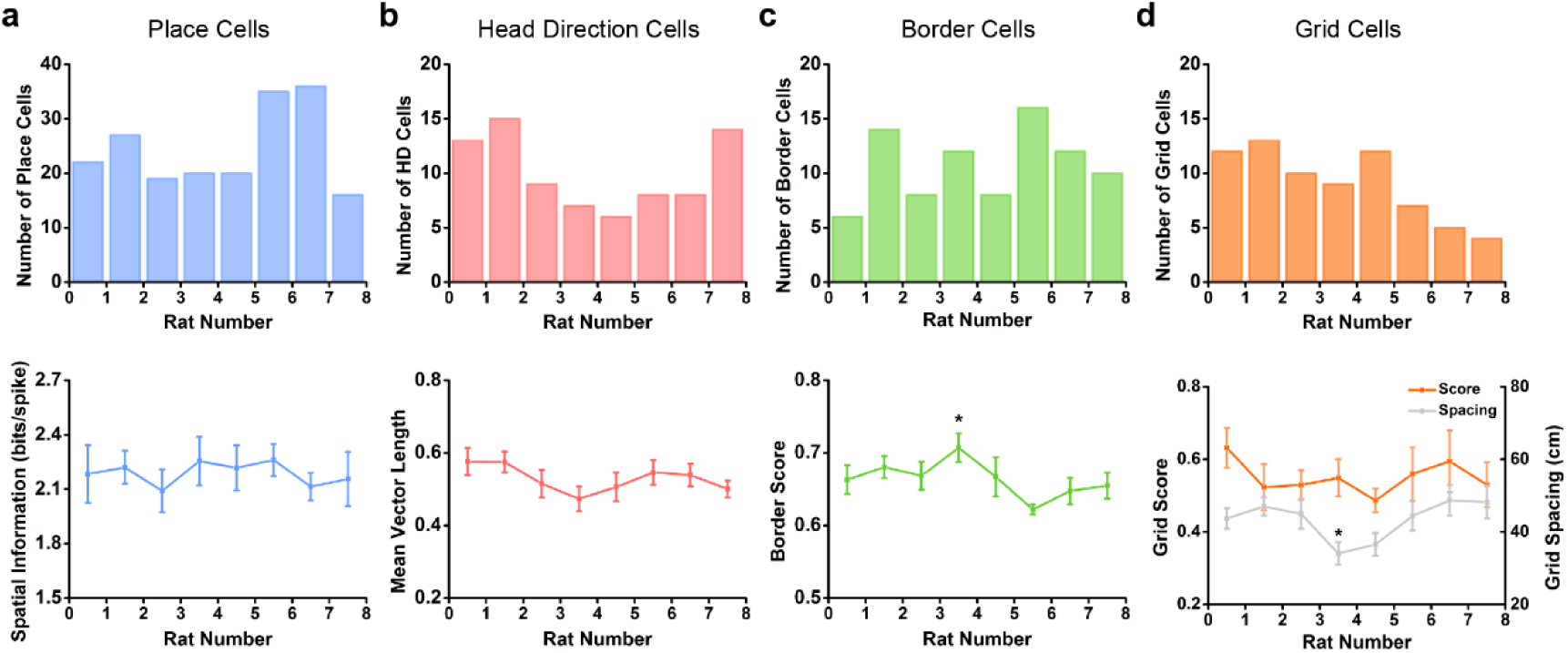
Distribution of number and characteristics of four different somatosensory spatial cell types. (**a-d**) Distribution of number (upper panels) and spatial characteristics (bottom panels) of identified somatosensory place cells (**a**), head direction cells (**b**), border cells (**c**) and grid cells (**d**) across different animals.

**Fig. S5.**
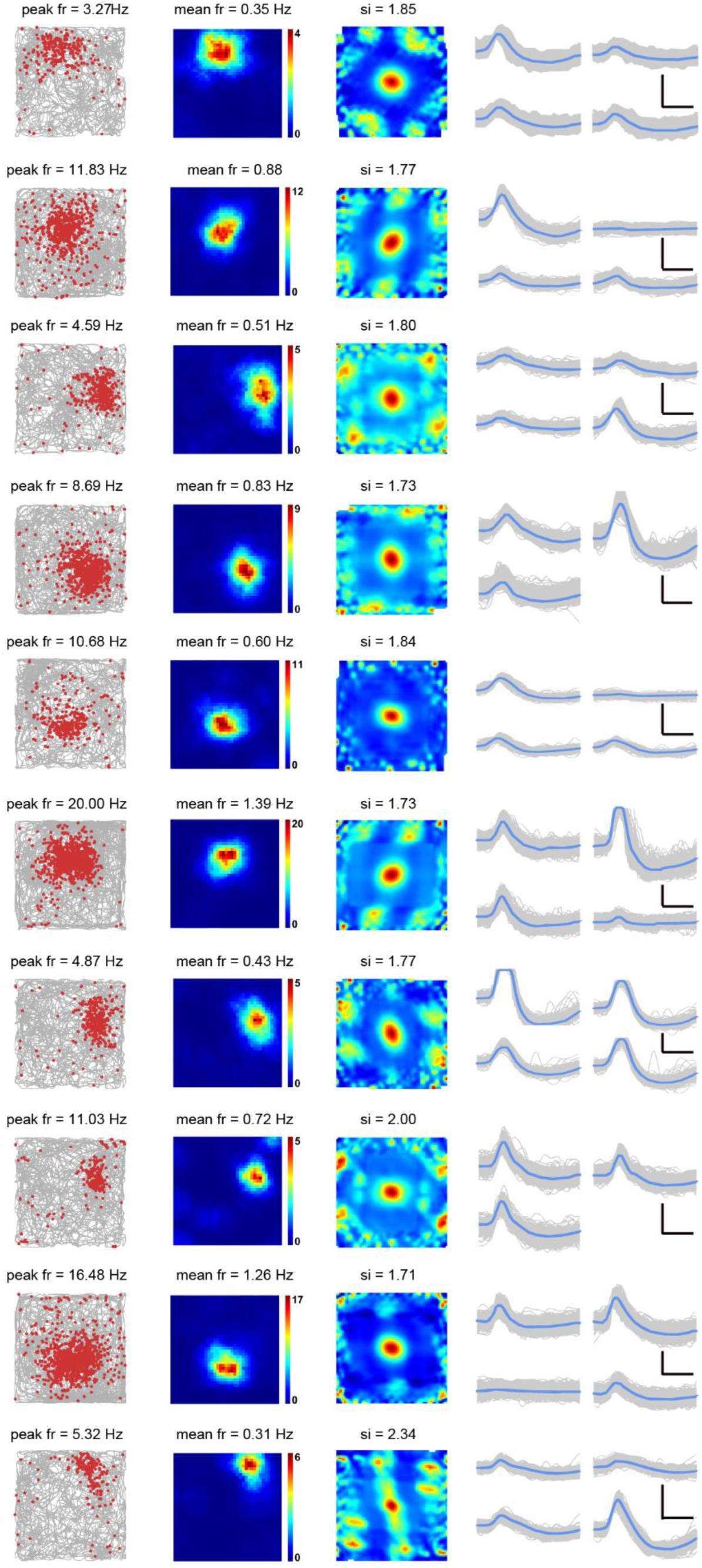
More examples of somatosensory place cells recorded from the somatosensory cortex. (**a-c**) Representative somatosensory place cells with bin coverage over 90%. Trajectory (grey line) with superimposed spike locations (red dots) (left column); heat maps of spatial firing rate (middle column) and autocorrelation (right column) are color-coded with dark blue indicating minimal firing rate and dark red indicating maximal firing rate. The scale of the autocorrelation maps is twice that of the spatial firing rate maps. Peak firing rate (fr), mean firing rate (fr) and spatial information (si) for each representative cell are labelled at the top of the panels. Spike waveforms on four electrodes are shown on the right column. Scale bar, 150 µV, 300 µs.

**Fig. S6.**
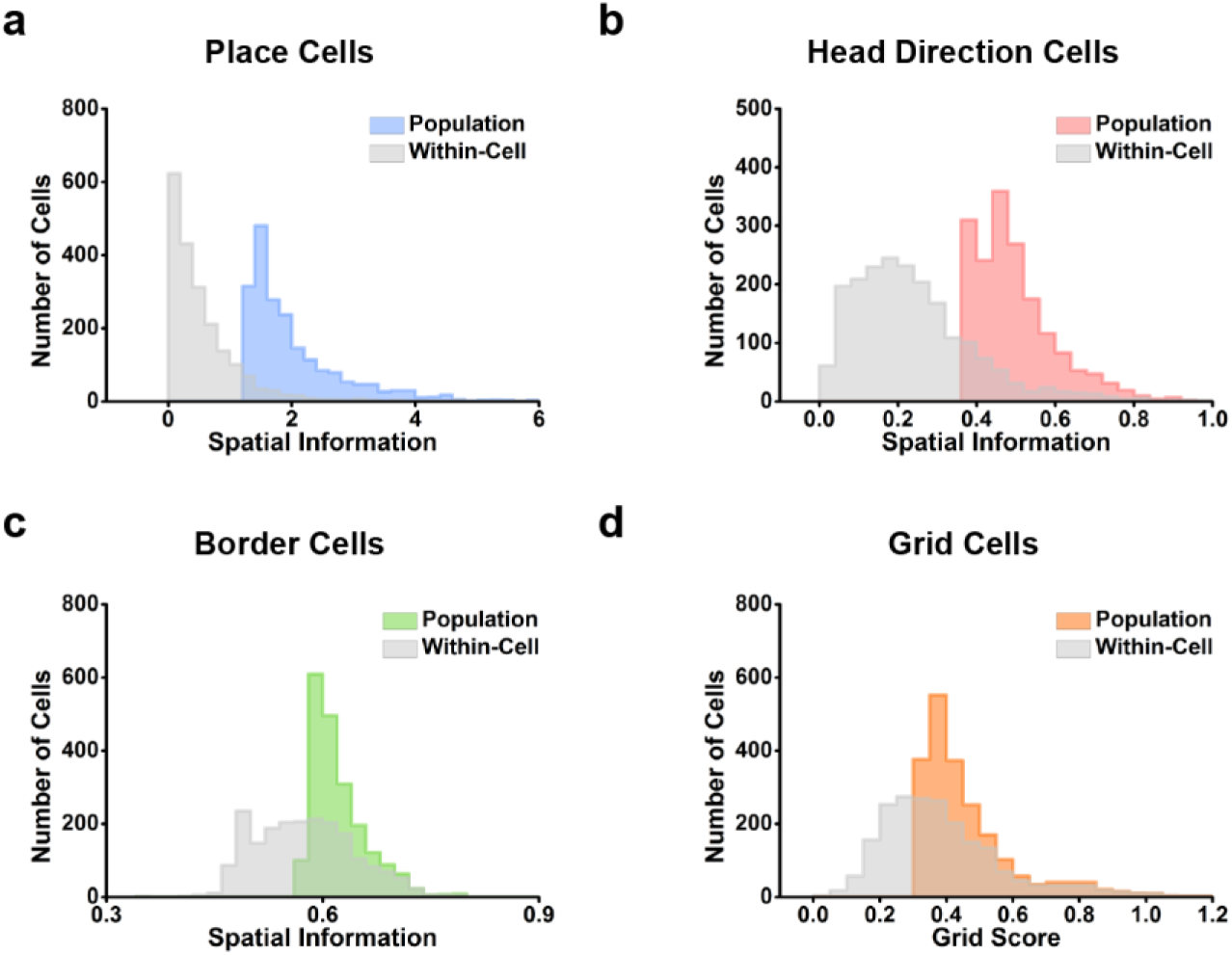
Comparison of spatial threshold defined by population shuffling and within-cell shuffling for four different somatosensory spatial cell types. (**a-d**) Histograms showing the 99^th^ percentile significance level of each randomly shuffled distribution for population shuffling and within-cell shuffling of all 2025 identified single units for identified somatosensory place cells (**a**), head direction cells (**b**), border cells (**c**) and grid cells (**d**).

**Fig. S7.**
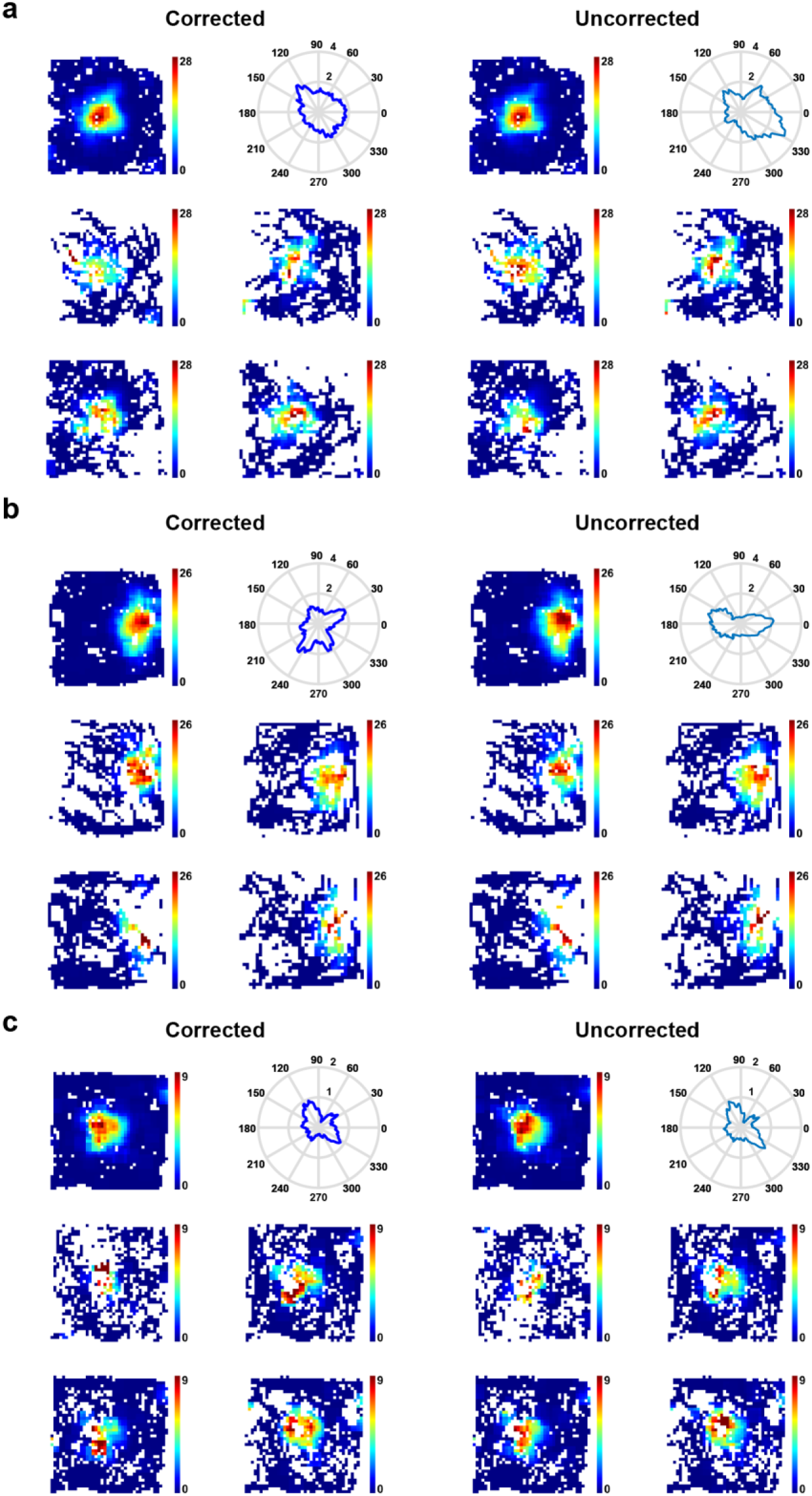
Quantification of spatial responses of somatosensory place cells using the maximum likelihood factorial model. (**a-c**) Firing rate maps of three representative place cells from Fig. 1b. Left column shows the corrected locational and directional firing rate maps using the maximum-likelihood correction approach; right column shows the corresponding uncorrected firing rate maps. Spatial firing rate maps in four directions are shown in the lower panels. Note the similar locational responses in four different directions.

**Fig. S8.**
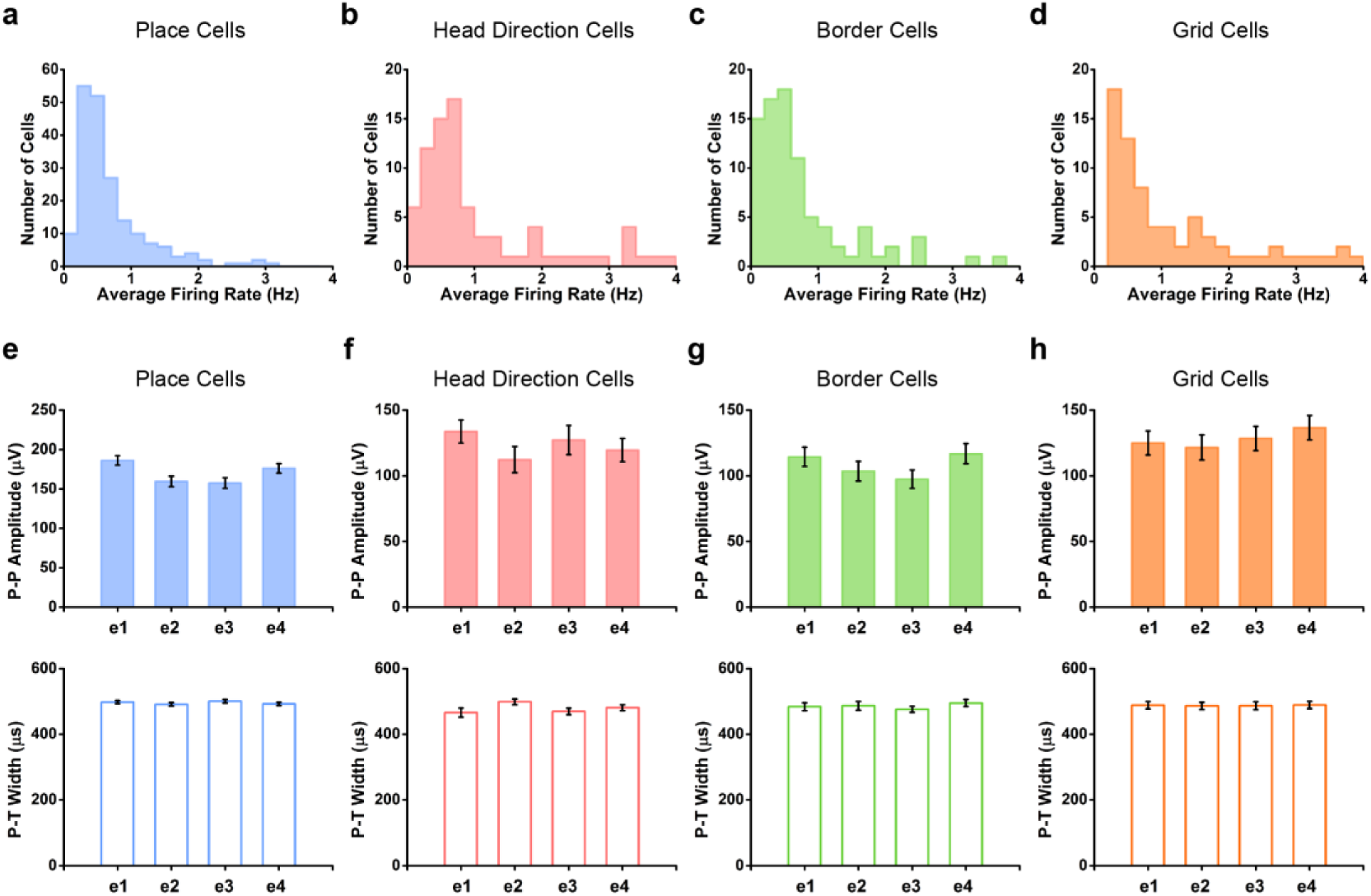
Distribution of the average firing rate and summary of spike waveform for four different somatosensory spatial cell types. (**a-d**) Histograms showing the average firing rate of identified somatosensory place cells (**a**), head direction cells (**b**), border cells (**c**) and grid cells (**d**). (**e-h**) Histograms showing the peak-to-peak amplitudes (upper panels) and peak to trough width (bottom panels) of spike waveforms on four electrodes of identified somatosensory place cells (**e**), head direction cells (**f**), border cells (**g**) and grid cells (**h**).

**Fig. S9.**
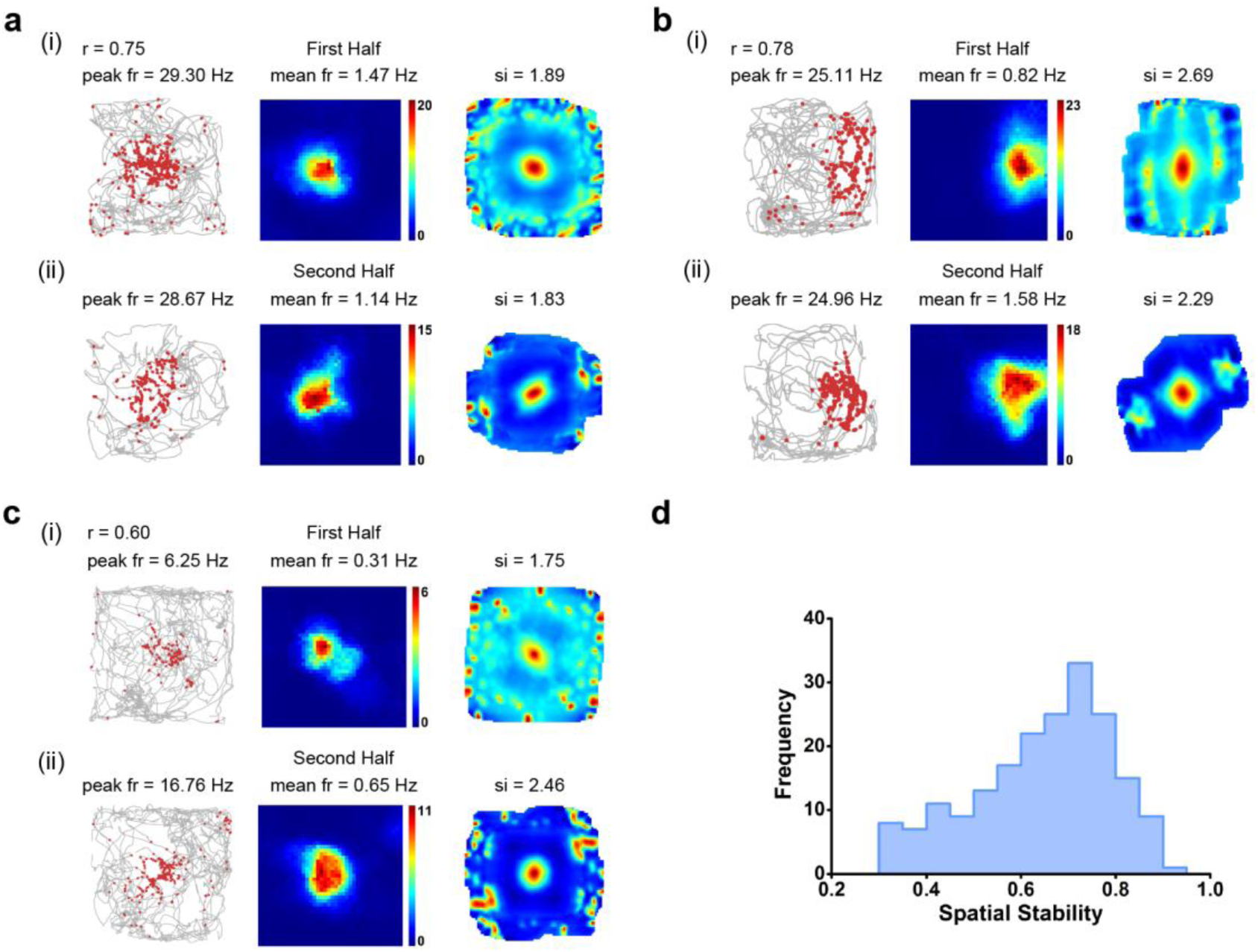
Spatial stability of somatosensory place cells. (**a-c**) Intra-trial spatial stability between the first and second halves of three representative place cells from Fig. 1b. Trajectory (grey line) with superimposed spike locations (red dots) (left column); rate maps (middle column) and autocorrelation maps (right column) for the first half (**i**) and the second half (**ii**) of the trials. Firing rate is color-coded with blue indicating minimum firing rate and red indicating maximum firing rate. The scale of the autocorrelation maps is twice that of the spatial firing rate maps. Peak firing rate (fr), mean firing rate (fr) and spatial information (si) are labelled at the top of the plots. Pearson’s correlation coefficients of firing rate maps between the first and second halves are indicated with *r* at the top-left corner. (**d)** Distribution of spatial stability of firing rate maps between the first and second halves of all identified somatosensory place cells.

**Fig. S10.**
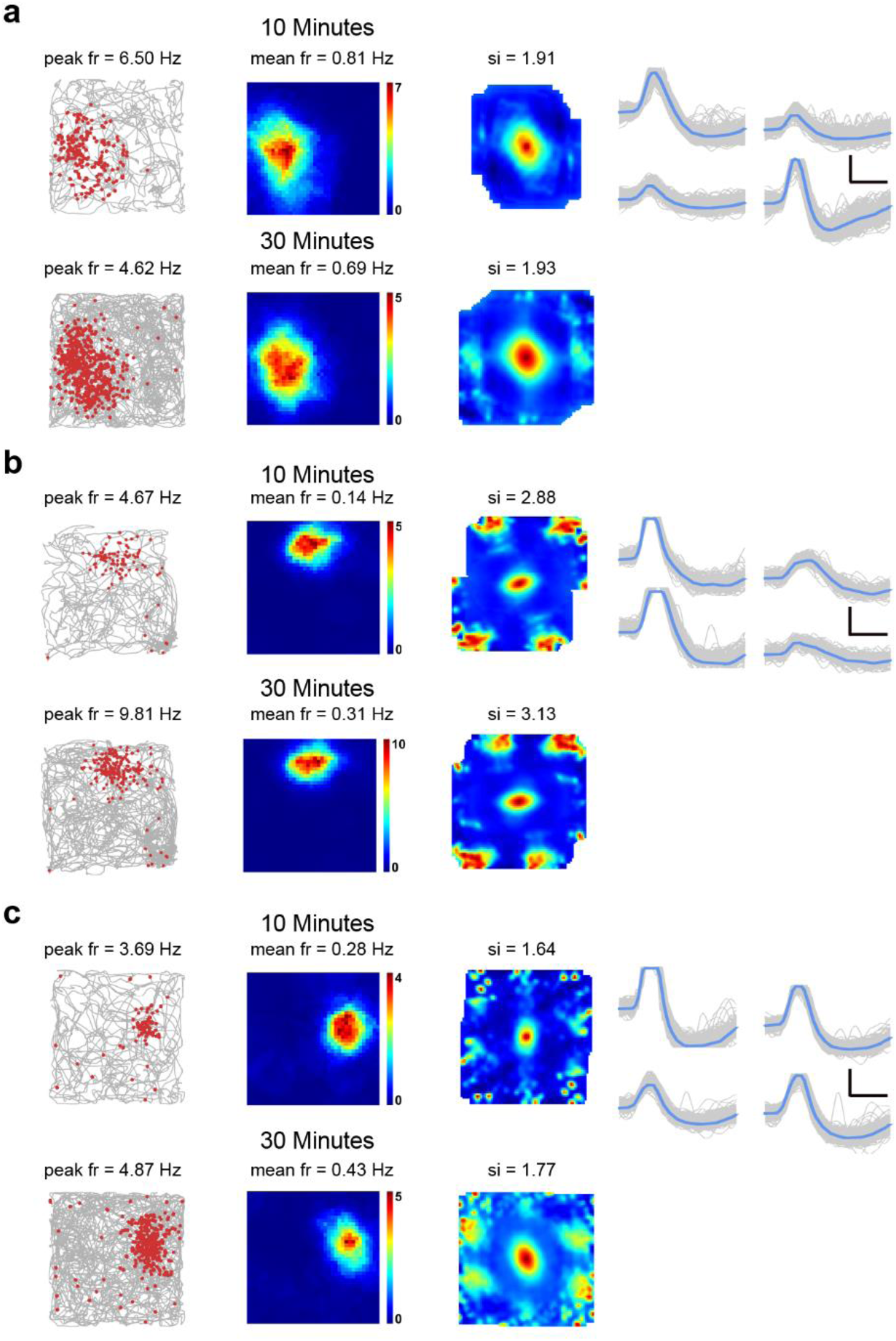
Persistence of somatosensory place fields from longer recording sessions. (**a**-**c**) Spatial stability of three representative somatosensory place cells between the short (top panels) and longer (bottom panels) recording sessions. Trajectory (grey line) with superimposed spike locations (red dots) (left column); spatial firing rate maps (middle column) and autocorrelation diagrams (right column). Firing rate is color-coded with blue indicating minimum firing rate and red indicating maximum firing rate. The scale of the autocorrelation maps is twice that of the spatial firing rate maps. Peak firing rate (fr), mean firing rate (fr) and spatial information (si) for each recording session are labelled at the top of the panels. Spike waveforms on four electrodes are shown on the right column. Scale bar, 150 µV, 300 µs.

**Fig. S11.**
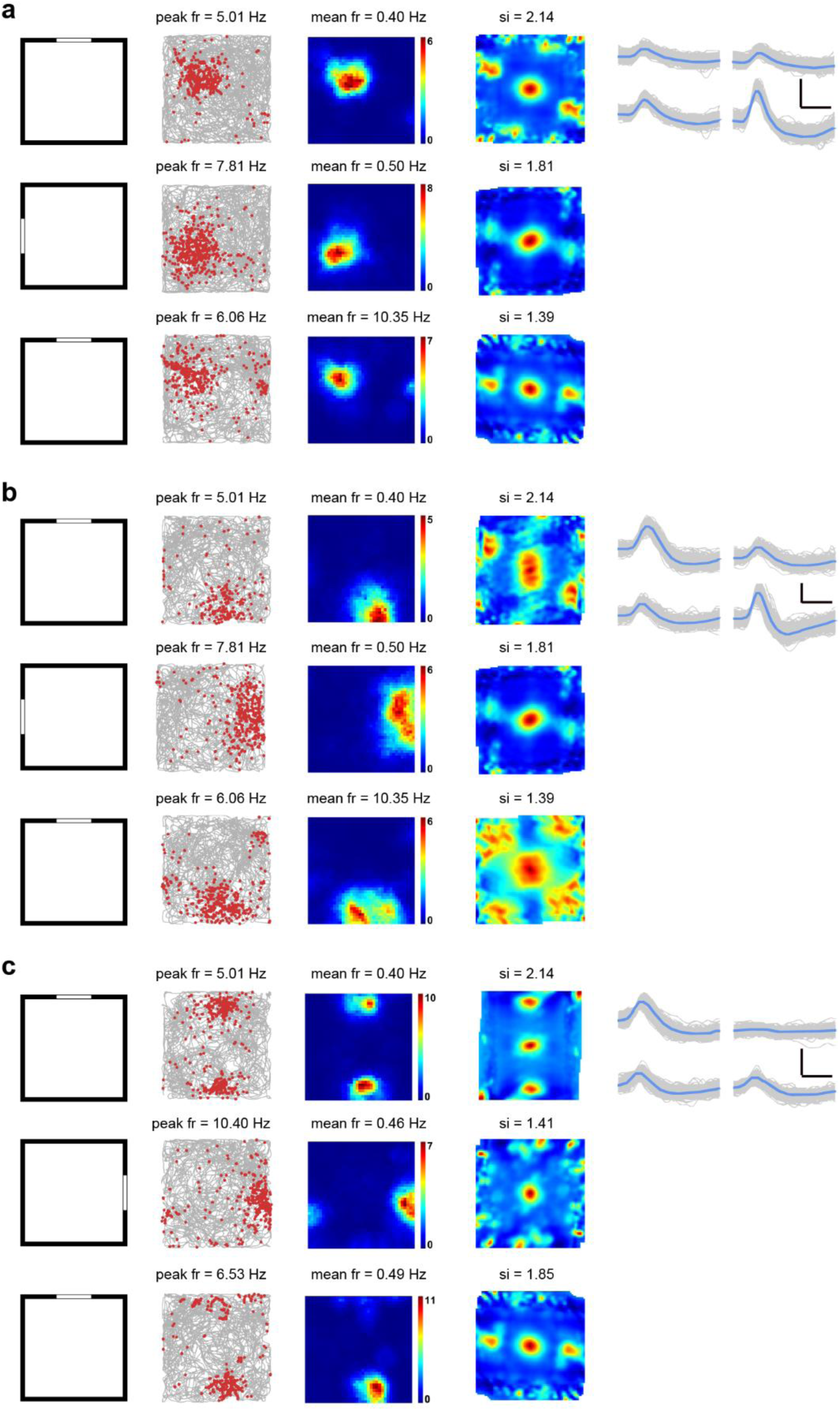
Cue-rotation control of somatosensory place cell coding. (**a-c**) Each panel shows the response of the same somatosensory place cell during three sessions of the cue-rotation condition. Top panels, before the cue-rotation; middle panels, counterclockwise or clockwise 90° of cue-rotation; bottom panels, cue-rotation back to the original condition. The cue card is represented by a white arc in each panel. The experimental diagram (left column); trajectory (grey line) with superimposed spike locations (red dots) (middle left column); spatial firing rate maps (middle right column) and autocorrelation diagrams (right column). Firing rate is color-coded with blue indicating minimum firing rate and red indicating maximum firing rate. The scale of the autocorrelation maps is twice that of the spatial firing rate maps. Peak firing rate (fr), mean firing rate (fr) and spatial information (si) for each recording session are labelled at the top of the panels. Spike waveforms on four electrodes are shown on the right column. Scale bar, 150 µV, 300 µs.

**Fig. S12.**
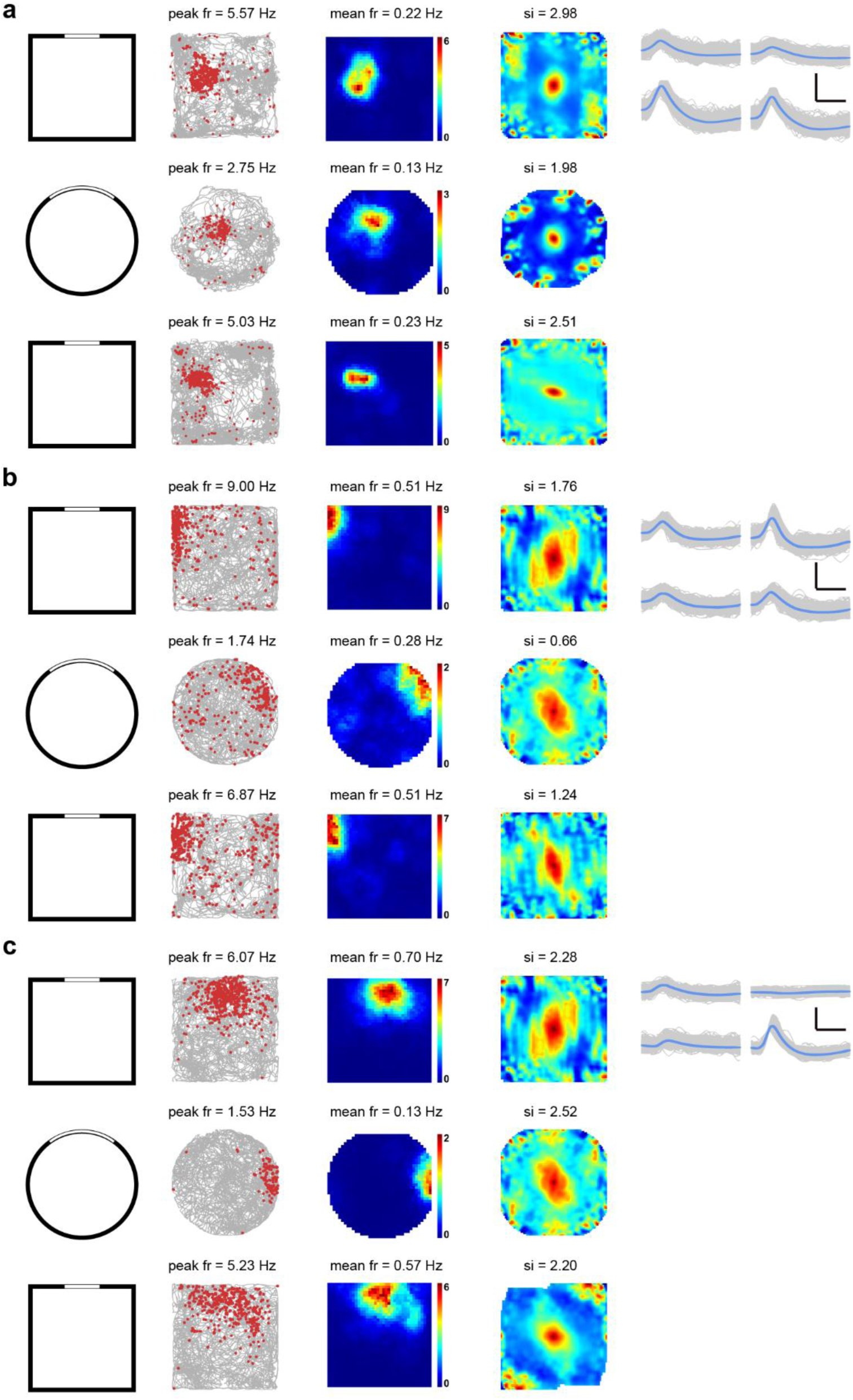
Remapping of somatosensory place cells in different environments. (**a**-**c**) Spatial responses of three representative somatosensory place cells in different boxes in a constant location. The experimental diagram (left column); trajectory (grey line) with superimposed spike locations (red dots) (middle left column); spatial firing rate maps (middle right column) and autocorrelation diagrams (right column). Firing rate is color-coded with blue indicating minimum firing rate and red indicating maximum firing rate. The scale of the autocorrelation maps is twice that of the spatial firing rate maps. Peak firing rate (fr), mean firing rate (fr) and spatial information (si) for each recording session are labelled at the top of the panels. Spike waveforms on four electrodes are shown on the right column. Scale bar, 150 µV, 300 µs.

**Fig. S13.**
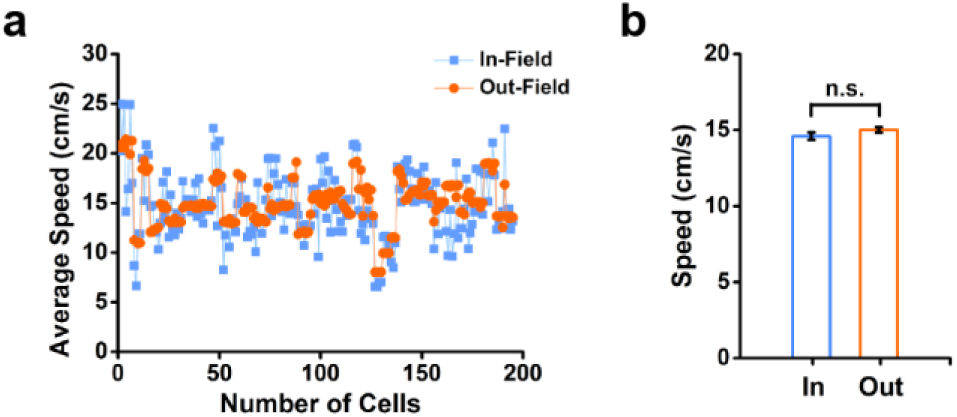
Distribution of the average in-field and out-field running speed for place cells in the somatosensory cortex. (**a**) The distribution of the average running speed within and outside the firing fields of all identified place cells in the somatosensory cortex. (**b**) The comparison of the average in-field and out-field running speed. *n* = 195, *P* = 0.10, two-tailed paired *t-*test, n.s., not significant.

**Fig. S14.**
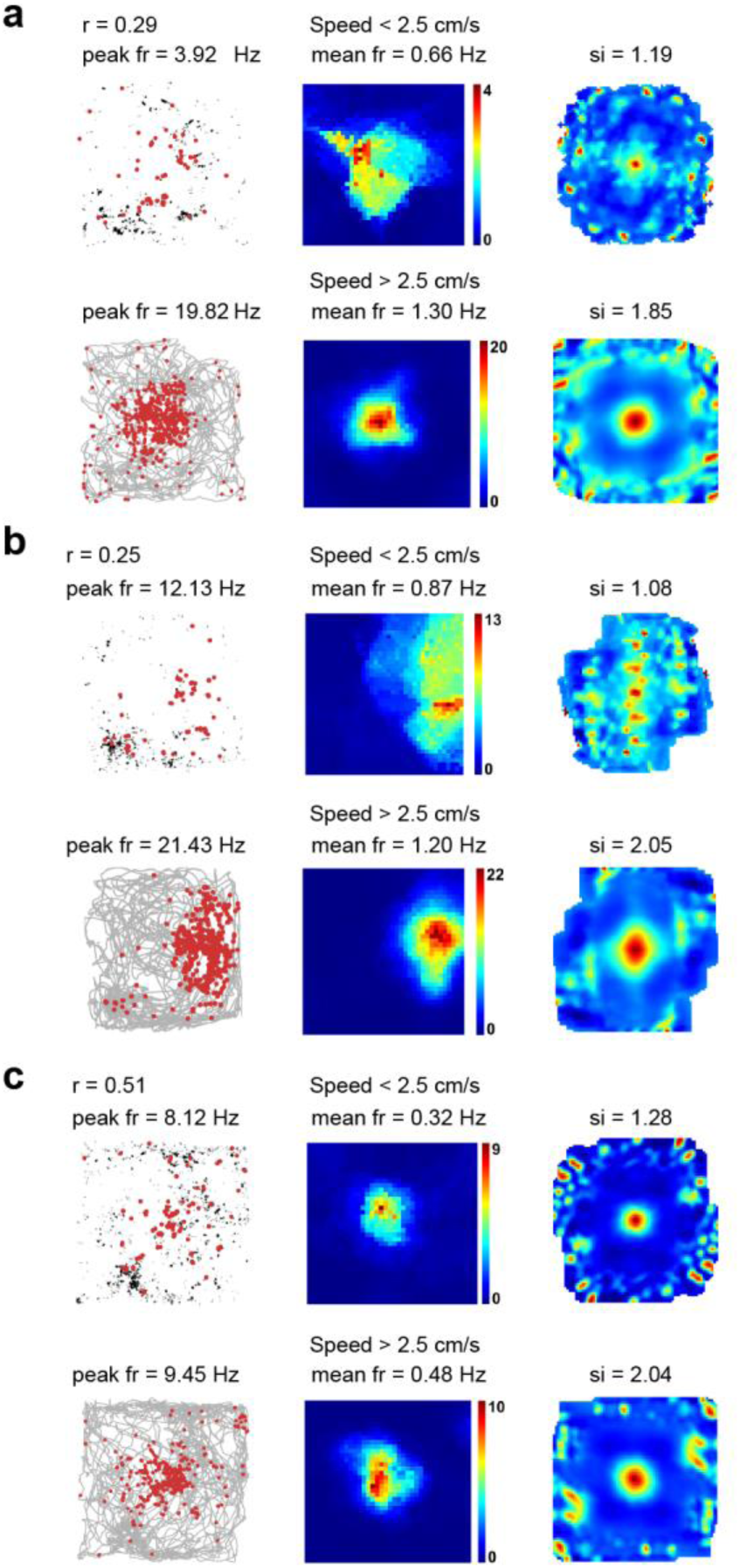
Comparison of the spatial response of somatosensory place cells between active running and slow mobility. (**a-c**) Comparison of spatial response of representative place cells from Fig. 1b with instantaneous running speeds <2.5 cm/s and instantaneous running speeds > 2.5 cm/s, respectively. Trajectory (grey line) with superimposed spike locations (red dots) (left column); rate maps (middle column) and autocorrelation maps (right column) for instantaneous running speeds <2.5 cm/s and instantaneous running speeds > 2.5 cm/s, respectively. Firing rate is color-coded with blue indicating minimum firing rate and red indicating maximum firing rate. The scale of the autocorrelation maps is twice that of the spatial firing rate maps. Peak firing rate (fr), mean firing rate (fr) and spatial information (si) are labelled at the top of the plots. Pearson’s correlation coefficients between two firing rate maps when the instantaneous running speeds <2.5 cm/s and instantaneous running speeds > 2.5 cm/s are indicated with *r* at the top-left corner.

**Fig. S15.**
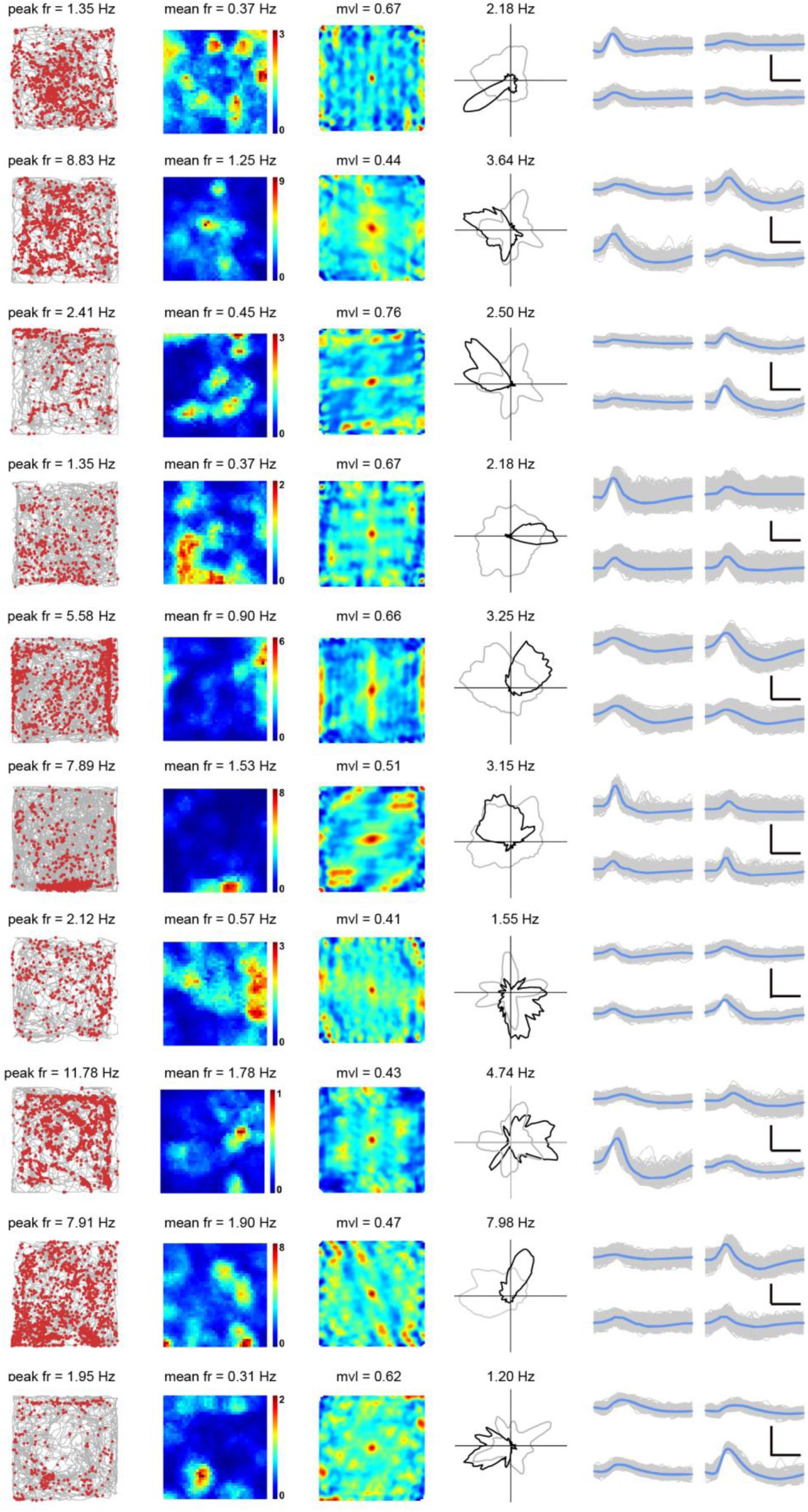
More examples of somatosensory head directions cells recorded from the somatosensory cortex. (**a-c**) Representative somatosensory head direction cells with bin coverage over 90%. Trajectory (grey line) with superimposed spike locations (red dots) (left column); spatial firing rate maps (middle left column), autocorrelation diagrams (middle right column) and head direction tuning curves (black) plotted against dwell-time polar plot (grey) (right column). Firing rate is color-coded with dark blue indicating minimal firing rate and dark red indicating maximal firing rate. The scale of the autocorrelation maps is twice that of the spatial firing rate maps. Peak firing rate (fr), mean firing rate (fr), mean vector length (mvl) and angular peak rate for each representative head direction cell are labelled at the top of the panels. The directional plots show strong head direction tuning. Spike waveforms on four electrodes are shown on the right column. Scale bar, 150 µV, 300 µs.

**Fig. S16.**
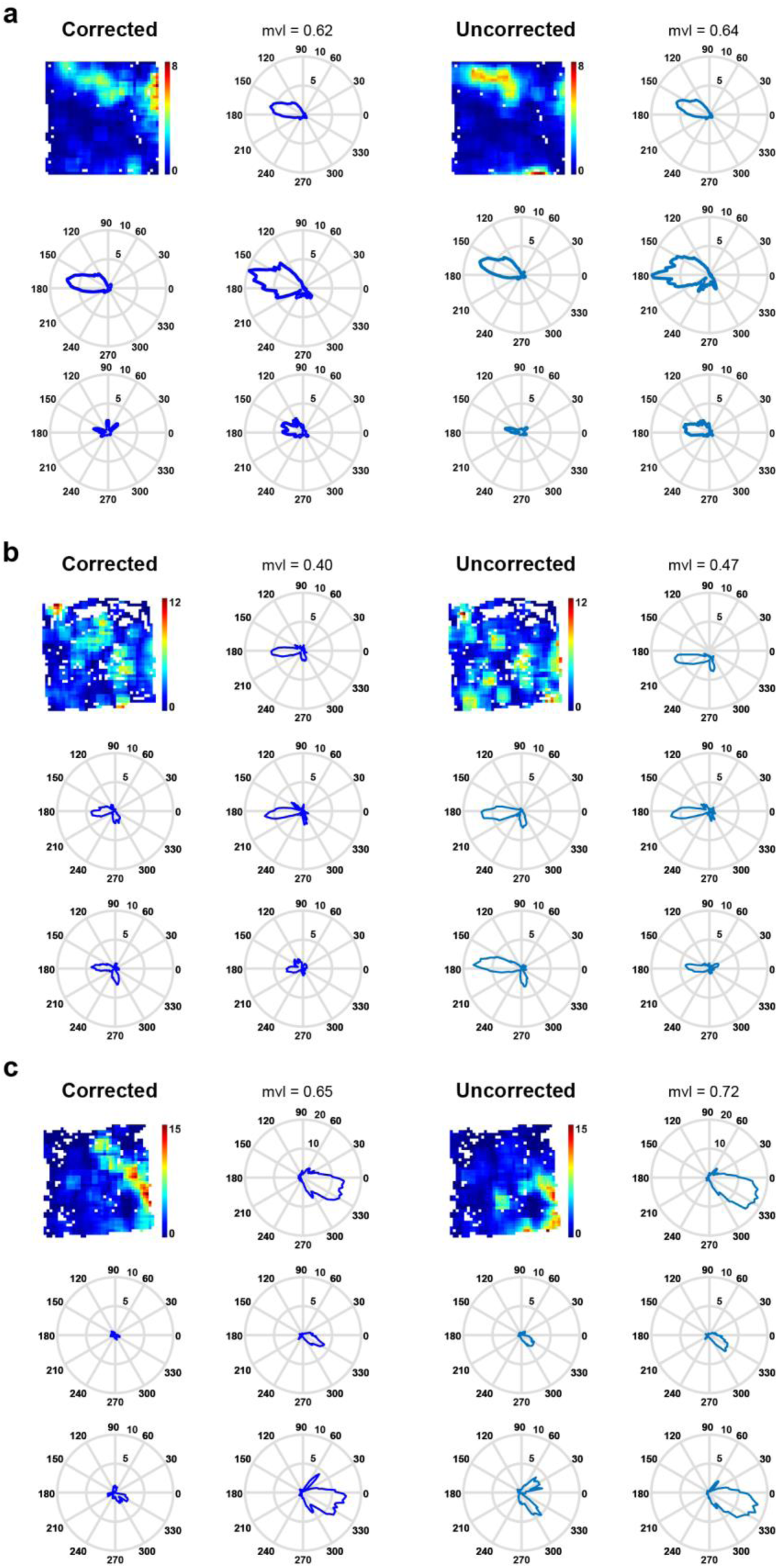
Quantification of head direction selectivity of somatosensory head direction cells using the maximum likelihood factorial model. (**a-c**) Comparison of head directionality of three representative head direction cells from Fig. 2a using a maximum-likelihood approach. Top left two panels show the corrected rate maps and polar plots under the maximum likelihood factorial model. Top right two panels show the uncorrected histograms. Mean vector length (mvl) is indicated at the top right corner of the polar plot panel. The head direction responses in each quadrant of the running box are shown in the four panels below. Somatosensory HD cells preserve the same sharp head direction selectivity in different parts of the running box regardless of possible inhomogeneous sampling of the animal’s locations and orientations.

**Fig. S17.**
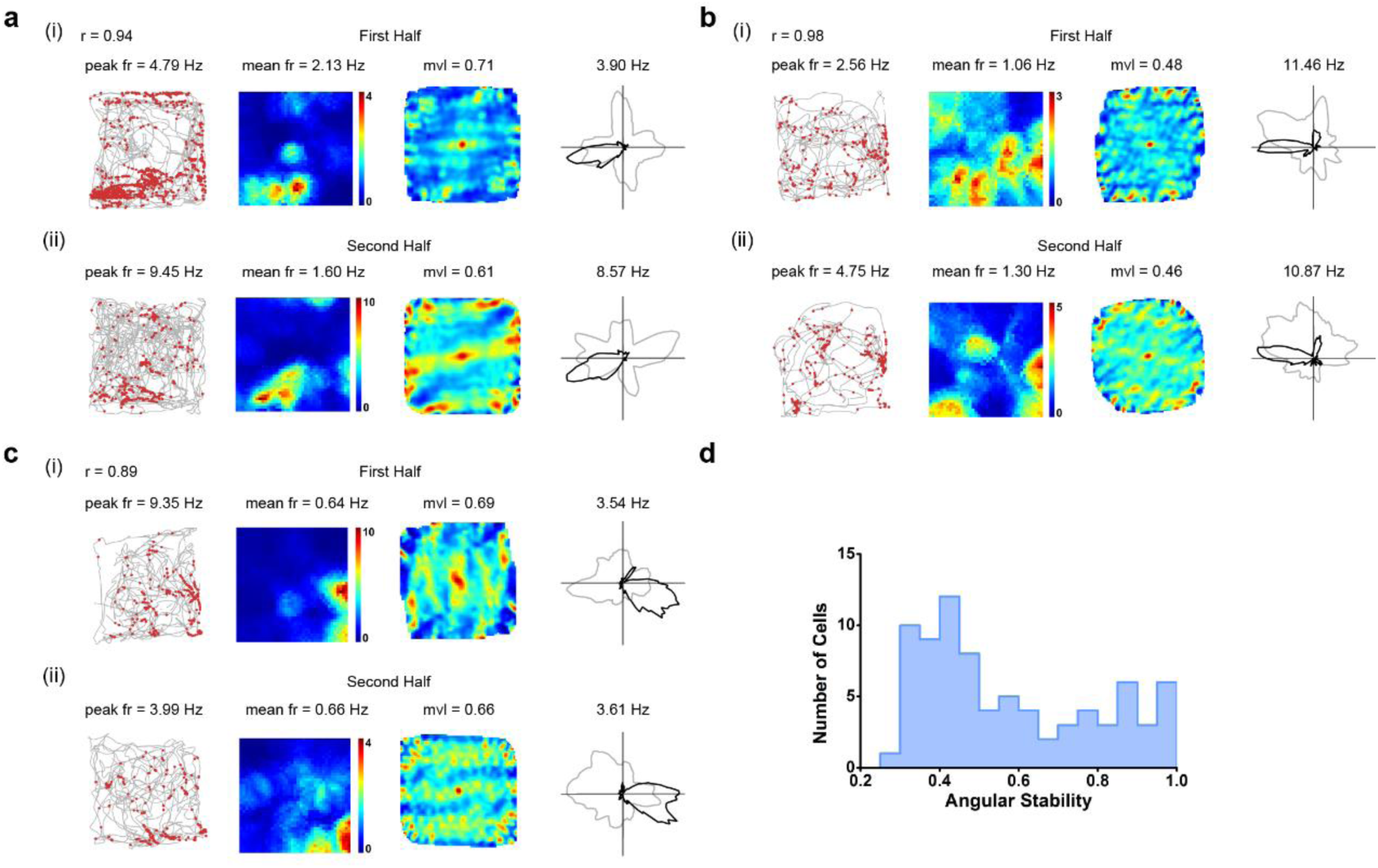
Angular stability of somatosensory head direction cells. (**a-c**) Intra-trial angular stability between the first and second halves of three representative head direction cells from Fig. 2a. Trajectory (grey line) with superimposed spike locations (red dots) (left column); spatial firing rate maps (middle left column), autocorrelation diagrams (middle right column) and head direction tuning curves (black) plotted against dwell-time polar plot (grey) (right column) for the first half (**i**) and the second half (**ii**) of the trials. Firing rate is color-coded with blue indicating minimum firing rate and red indicating maximum firing rate. The scale of the autocorrelation maps is twice that of the spatial firing rate maps. Peak firing rate (fr), mean firing rate (fr), mean vector length (mvl) and angular peak rate for each representative head direction cell are labelled at the top of the panels. Correlation coefficients of the distributed firing rate across all directional bins between the first and second halves of individual recording trials are indicated with *r*. (**d)** Population histogram of angular stability of the distributed firing rate across all directional bins between the first and second halves of all identified somatosensory head direction cells.

**Fig. S18.**
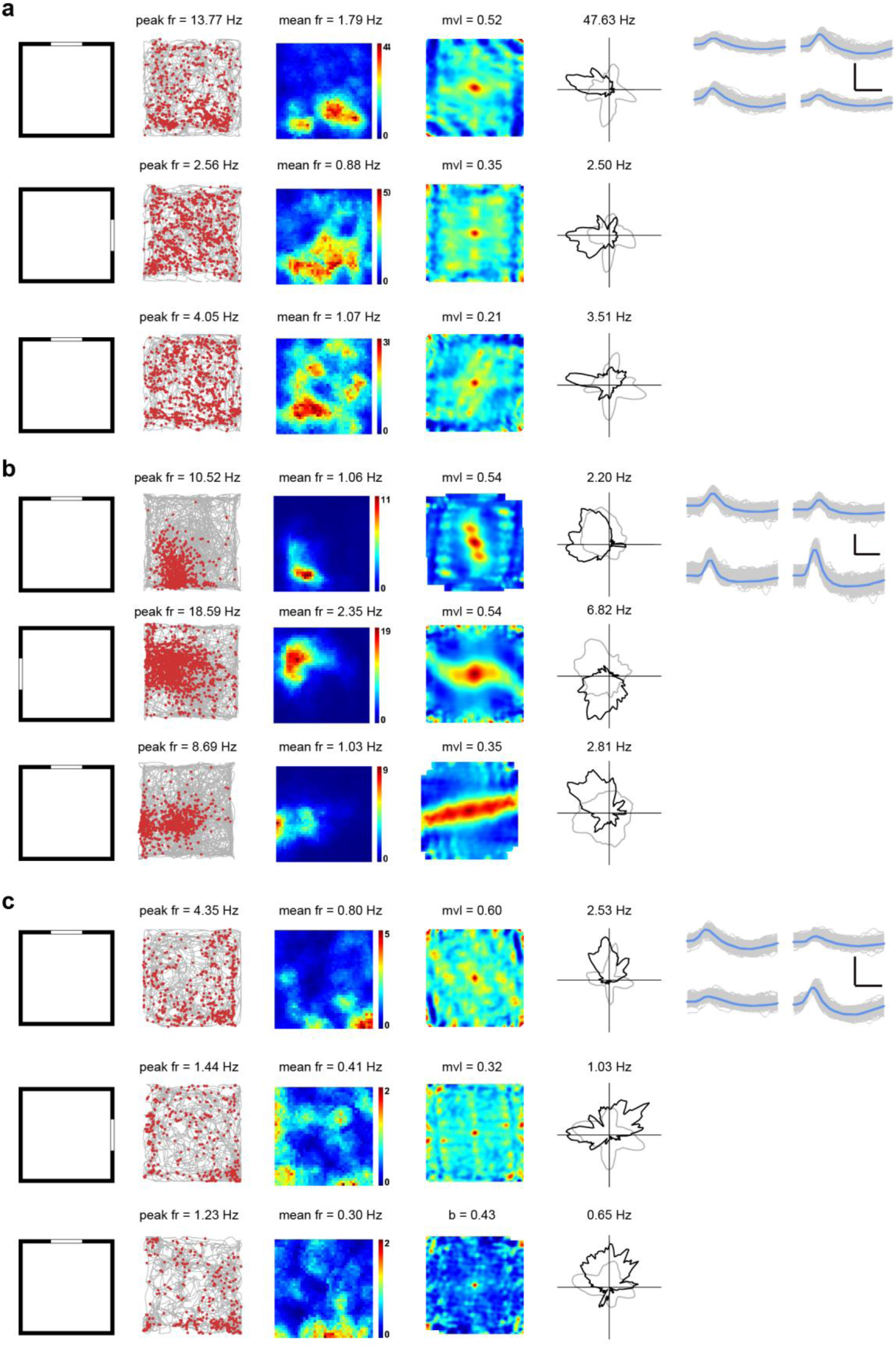
Cue-rotation control of somatosensory head directional responses. (**a-c**) Spatial responses of three representative somatosensory head direction cells during cue-rotation. Each panel shows the response of the same S1 head direction cell during three recording trails during the cue-rotation condition. Top panels, before the cue-rotation (**a**); middle panels, clockwise 90° of cue-rotation (**b**); bottom panels, cue-rotation back to the original condition (**c**). The cue card is represented by a white line in each panel. The experimental diagram (left column); trajectory (grey line) with superimposed spike locations (red dots) (middle left column); spatial firing rate maps (middle column), autocorrelation diagrams (middle right column) and head direction tuning curves (black) plotted against dwell-time polar plot (grey) (right column) for each recording trial. Firing rate is color-coded with blue indicating minimum firing rate and red indicating maximum firing rate. The scale of the autocorrelation maps is twice that of the spatial firing rate maps. Peak firing rate (fr), mean firing rate (fr), mean vector length (mvl) and angular peak rate for each representative head direction cell are labelled at the top of the panels. Spike waveforms on four electrodes are shown on the right column. Scale bar, 150 µV, 300 µs.

**Fig. S19.**
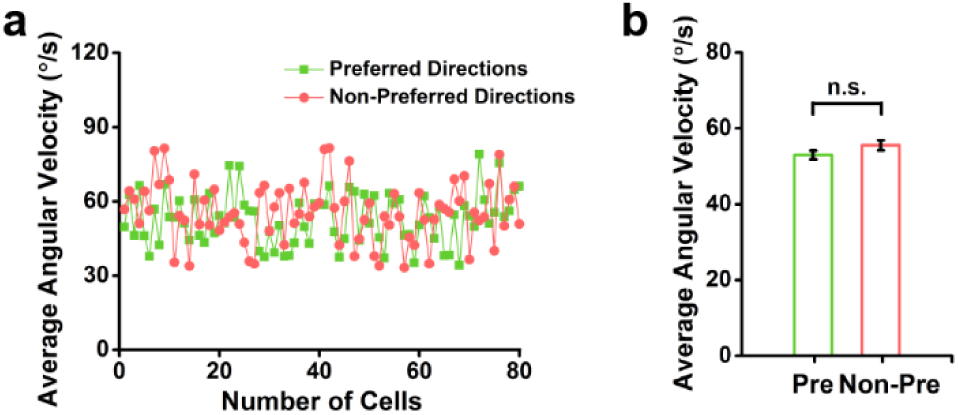
Distribution of the average angular velocity in the preferred and non-preferred firing directions for head direction cells in the somatosensory cortex. (**a**) The distribution of the average angular velocity in the “preferred firing directions” and “non-preferred firing directions” for all identified head direction cells in the somatosensory cortex. (**b**) The comparison of the angular velocity in the “preferred firing directions” and “non-preferred firing directions”. *n* = 80, *P* = 0.12, two-tailed paired *t-*test, n.s., not significant.

**Fig. S20.**
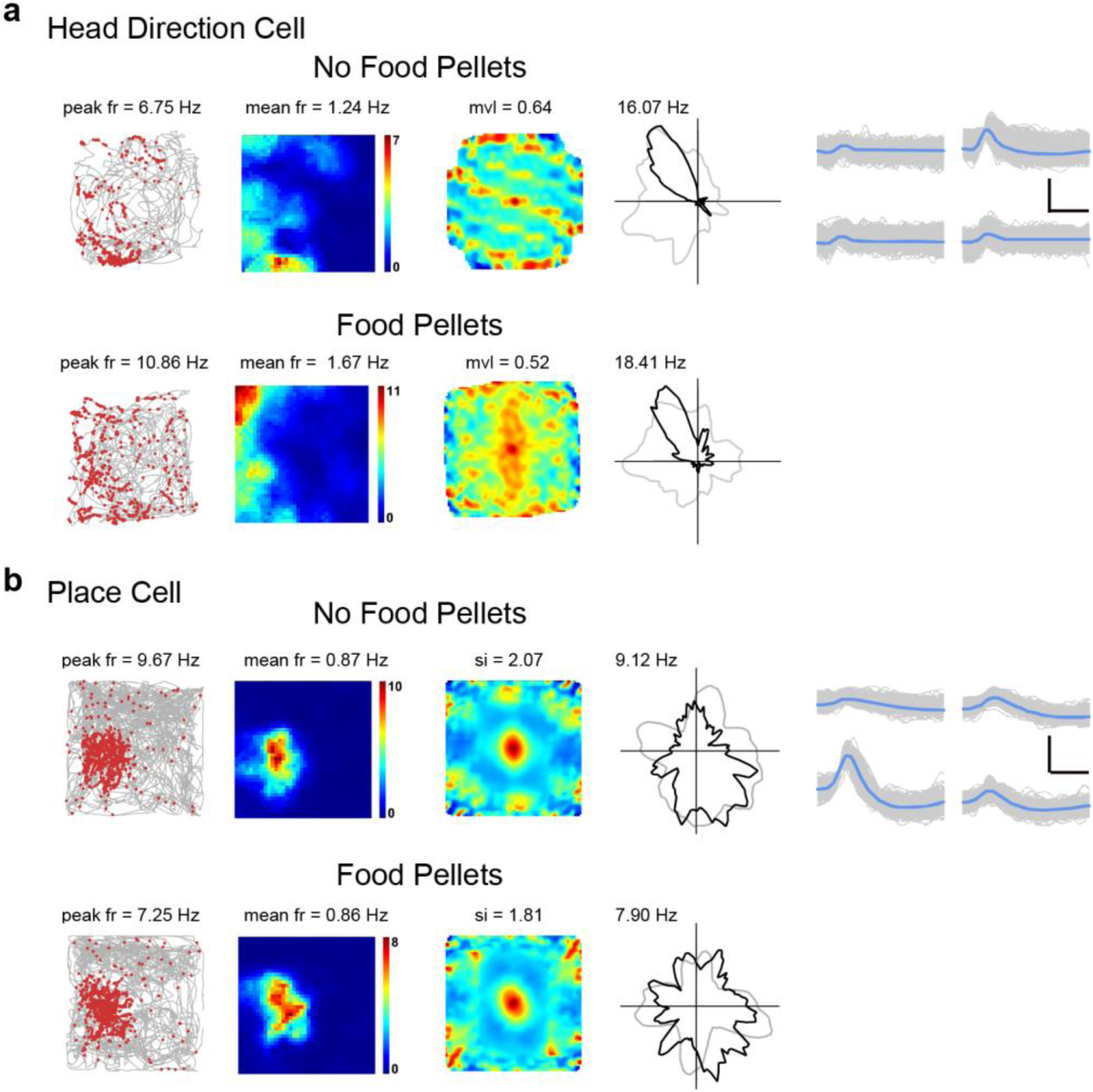
Somatosensory place and head direction cells in the presence or absence of food pellets. (**a**) The directional tuning of the same S1 head direction cell recorded with (top panels) and without (bottom panels) food pellets. (**b**) The firing fields of the same S1 place cell recorded with (top panels) and without (bottom panels) food pellets. Trajectory (grey line) with superimposed spike locations (red dots) (left column); spatial firing rate maps (middle left column), autocorrelation diagrams (middle right column) and head direction tuning curves (black) plotted against dwell-time polar plot (grey) (right column). Firing rate is color-coded with blue indicating minimum firing rate and red indicating maximum firing rate. The scale of the autocorrelation maps is twice that of the spatial firing rate maps. Peak firing rate (fr), mean firing rate (fr), mean vector length (mvl) or spatial information (si) and angular peak rate for each representative head direction cell are labelled at the top of the panels. Spike waveforms on four electrodes are shown on the right column. Scale bar, 150 µV, 300 µs.

**Fig. S21.**
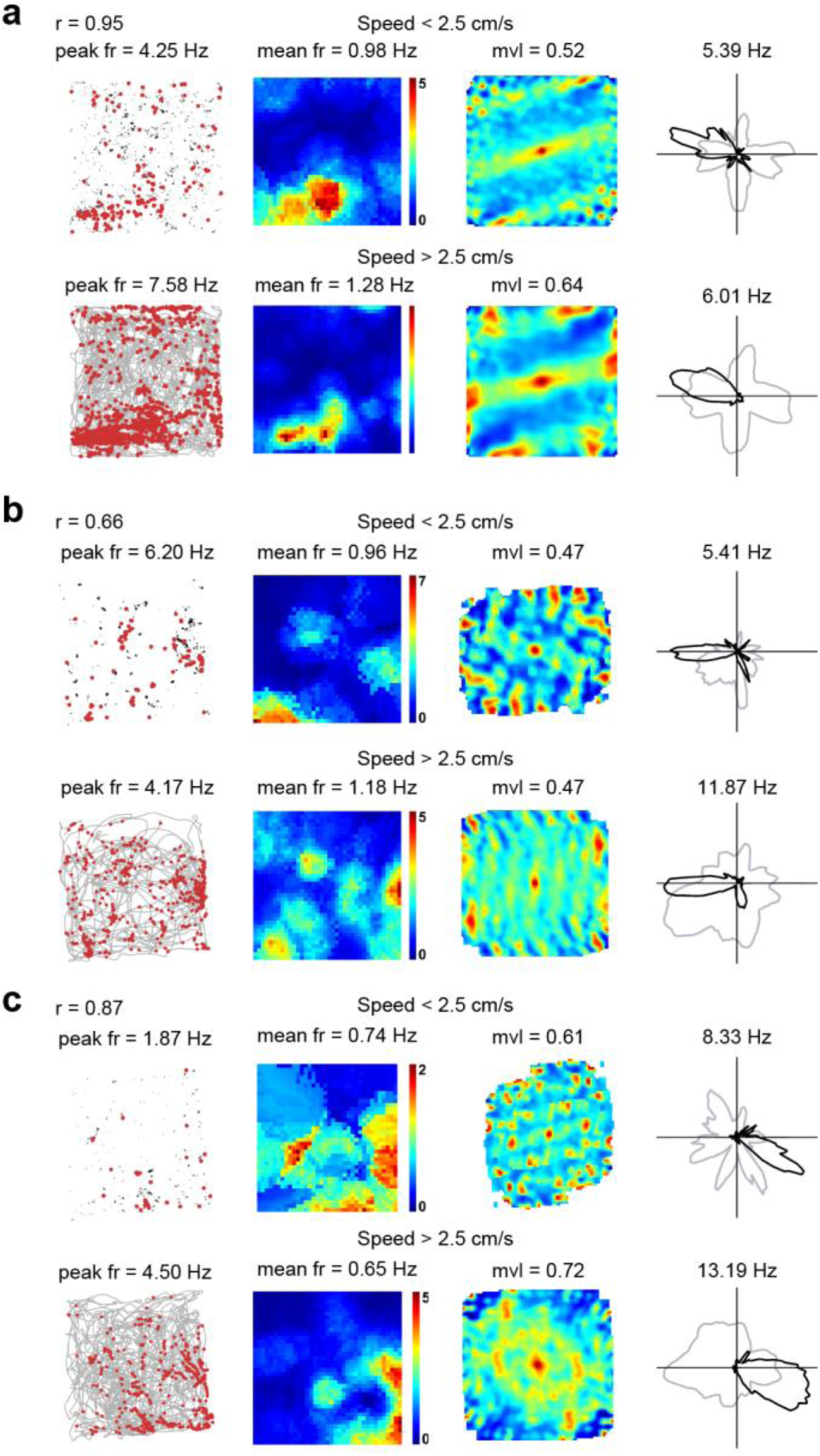
Comparison of the spatial response of somatosensory head direction cells between active running and slow immobility. (**a-c**) Comparison of the spatial response of representative head direction cells from Fig. 2a with instantaneous running speeds <2.5 cm/s and instantaneous running speeds > 2.5 cm/s, respectively. Trajectory (grey line) with superimposed spike locations (red dots) (left column); rate maps (middle left column), autocorrelation maps (middle right column) and head direction tuning curves (black) plotted against dwell-time polar plot (grey) (right column) for instantaneous running speeds <2.5 cm/s and instantaneous running speeds > 2.5 cm/s, respectively. Firing rate is color-coded with blue indicating minimum firing rate and red indicating maximum firing rate. The scale of the autocorrelation maps is twice that of the spatial firing rate maps. Peak firing rate (fr), mean firing rate (fr) and mean vector length (mvl) are labelled at the top of the plots. Correlation coefficients of the distributed firing rate across all directional bins when the instantaneous running speeds <2.5 cm/s and instantaneous running speeds > 2.5 cm/s are indicated with *r* at the top-left corner.

**Fig. S22.**
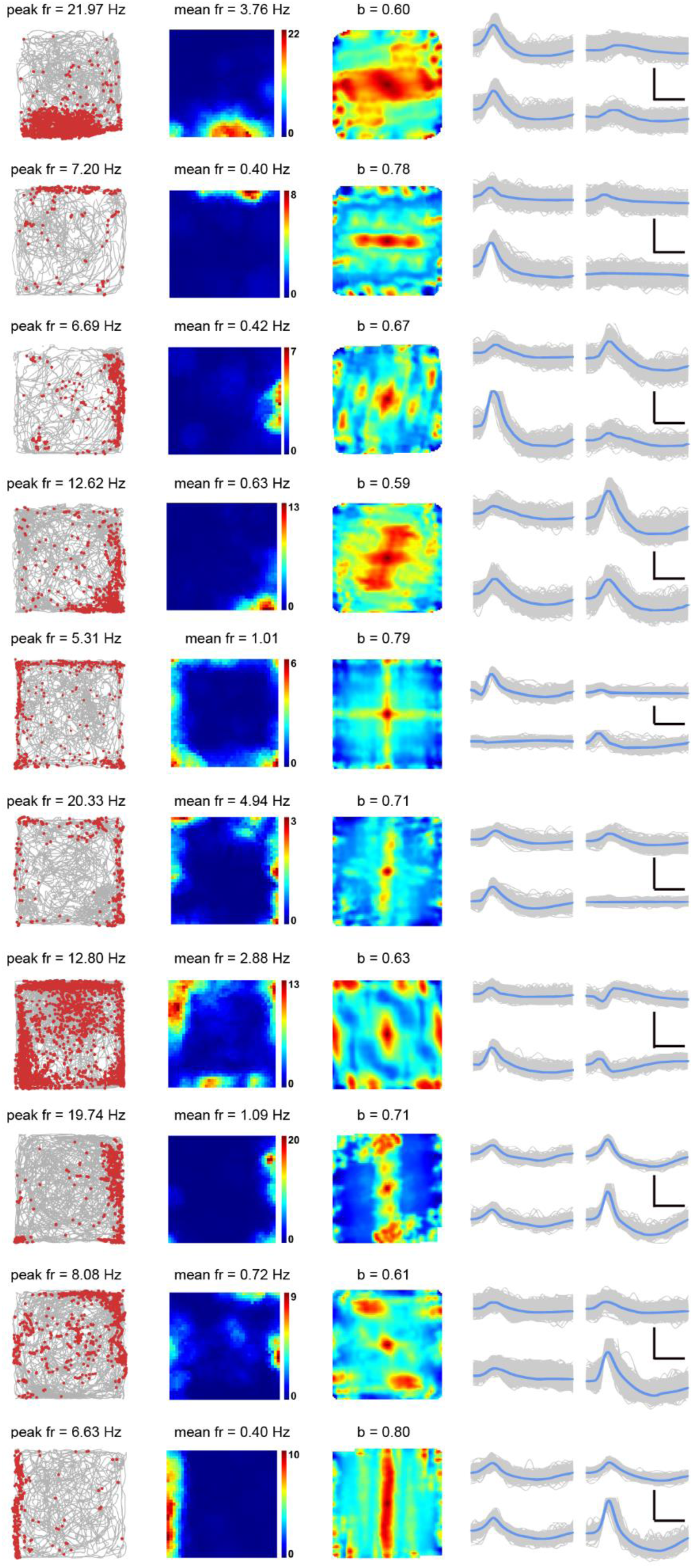
More examples of somatosensory border cells recorded from the somatosensory cortex. (**a-c**) Representative somatosensory border cells with bin coverage over 90%. Trajectory (grey line) with superimposed spike locations (red dots) (left column); spatial firing rate maps (middle left column), autocorrelation diagrams (middle right column) and head direction tuning curves (black) plotted against dwell-time polar plot (grey) (right column). Firing rate is color-coded with dark blue indicating minimal firing rate and dark red indicating maximal firing rate. The scale of the autocorrelation maps is twice that of the spatial firing rate maps. Peak firing rate (fr), mean firing rate (fr), mean vector length (mvl) and angular peak rate for each representative head direction cell are labelled at the top of the panels. The directional plots show strong head direction tuning. Spike waveforms on four electrodes are shown on the right column. Scale bar, 150 µV, 300 µs.

**Fig. S23.**
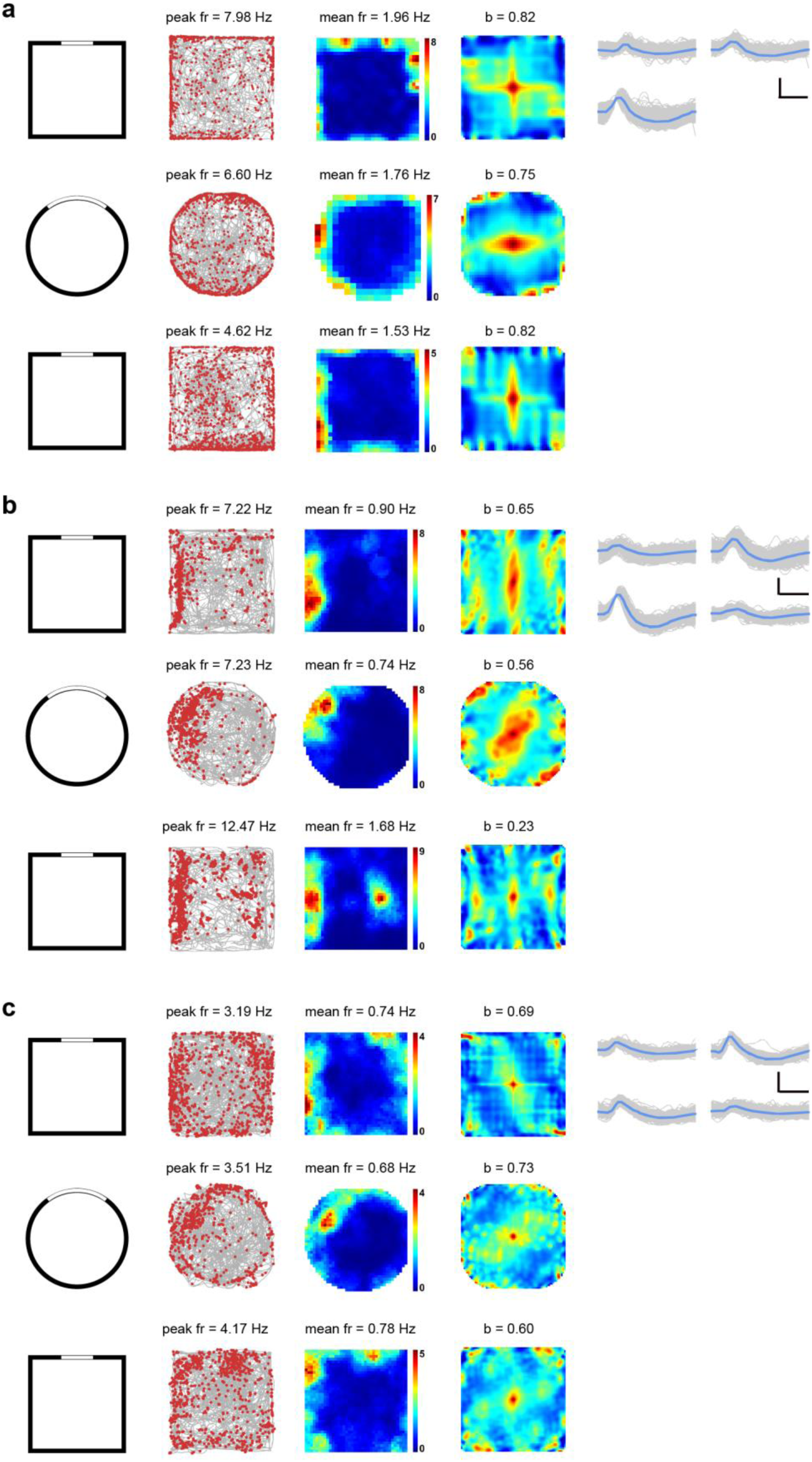
Preserved firing across different geometric shapes of somatosensory border cells. (**a-c**) Three representative S1 border cells preserve the firing patterns in the square and cylindrical box. Top panels, original square running box; middle panels, cylindrical box; bottom panels, back to the original square box. The experimental diagram (left column); trajectory (grey line) with superimposed spike locations (red dots) (middle left column); rate maps (middle right column) and autocorrelation diagrams (right column) for each recording trail. Firing rate is color-coded with blue indicating minimum firing rate and red indicating maximum firing rate. The scale of the autocorrelation maps is twice that of the spatial firing rate maps. Peak firing rate (fr), mean firing rate (fr) and border score for each recording session are labelled at the top of the panels. Spike waveforms on four electrodes are shown on the right column. Scale bar, 200 µV, 300 µs.

**Fig. S24.**
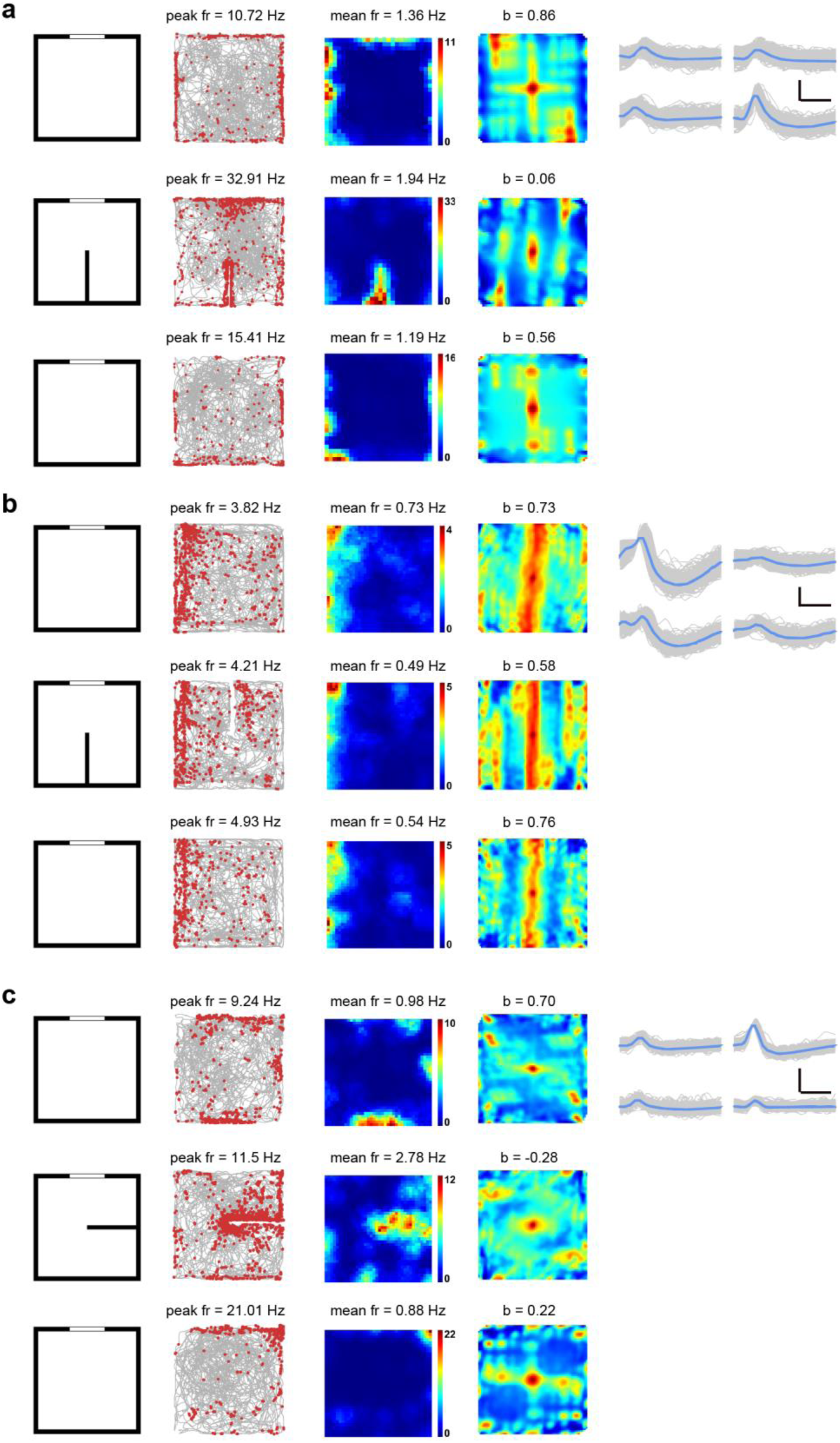
Spatial responses to the external inserts of somatosensory border cells. (**a-c**) Three representative somatosensory border cells recorded in the square enclosure with or without one external insert. Top panels, the square enclosure without insert; middle panels, the same square enclosure with an internal insert; bottom panels, the same square enclosure after removing the internal insert. The experimental diagram (left column); trajectory (grey line) with superimposed spike locations (red dots) (middle left column); rate maps (middle right column) and autocorrelation diagrams (right column) for each recording trail. Firing rate is color-coded with blue indicating minimum firing rate and red indicating maximum firing rate. The scale of the autocorrelation maps is twice that of the spatial firing rate maps. Peak firing rate (fr), mean firing rate (fr) and border score (b) for each recording session are labelled at the top of the panels. Spike waveforms on four electrodes are shown on the right column. Scale bar, 150 µV, 300 µs.

**Fig. S25.**
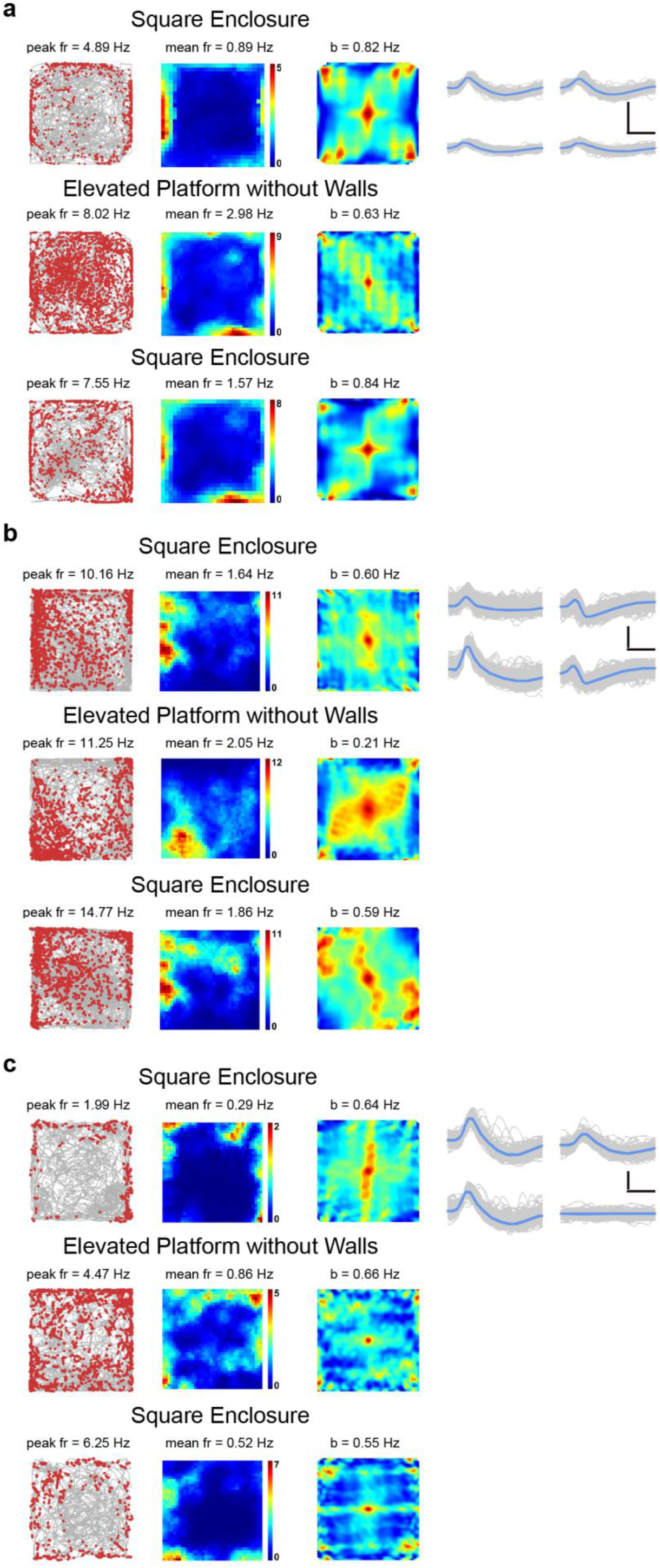
Somatosensory border cells recorded from the elevated platform without walls. (**a-c**) Spatial responses of three representative somatosensory border cells recorded in the square box, in the elevated platform without walls and back to the square box. Trajectory (grey line) with superimposed spike locations (red dots) (left column); heat maps of firing rate (middle column) and autocorrelation maps (right column). Firing rate is color-coded with blue indicating minimum firing rate and red indicating maximum firing rate. The scale of the autocorrelation maps is twice that of the spatial firing rate maps. Peak firing rate (fr), mean firing rate (fr) and border score (b) for each representative border cell are labelled at the top of the panels. Spike waveforms on four electrodes are shown on the right column. Scale bar, 150 µV, 300 µs.

**Fig. S26.**
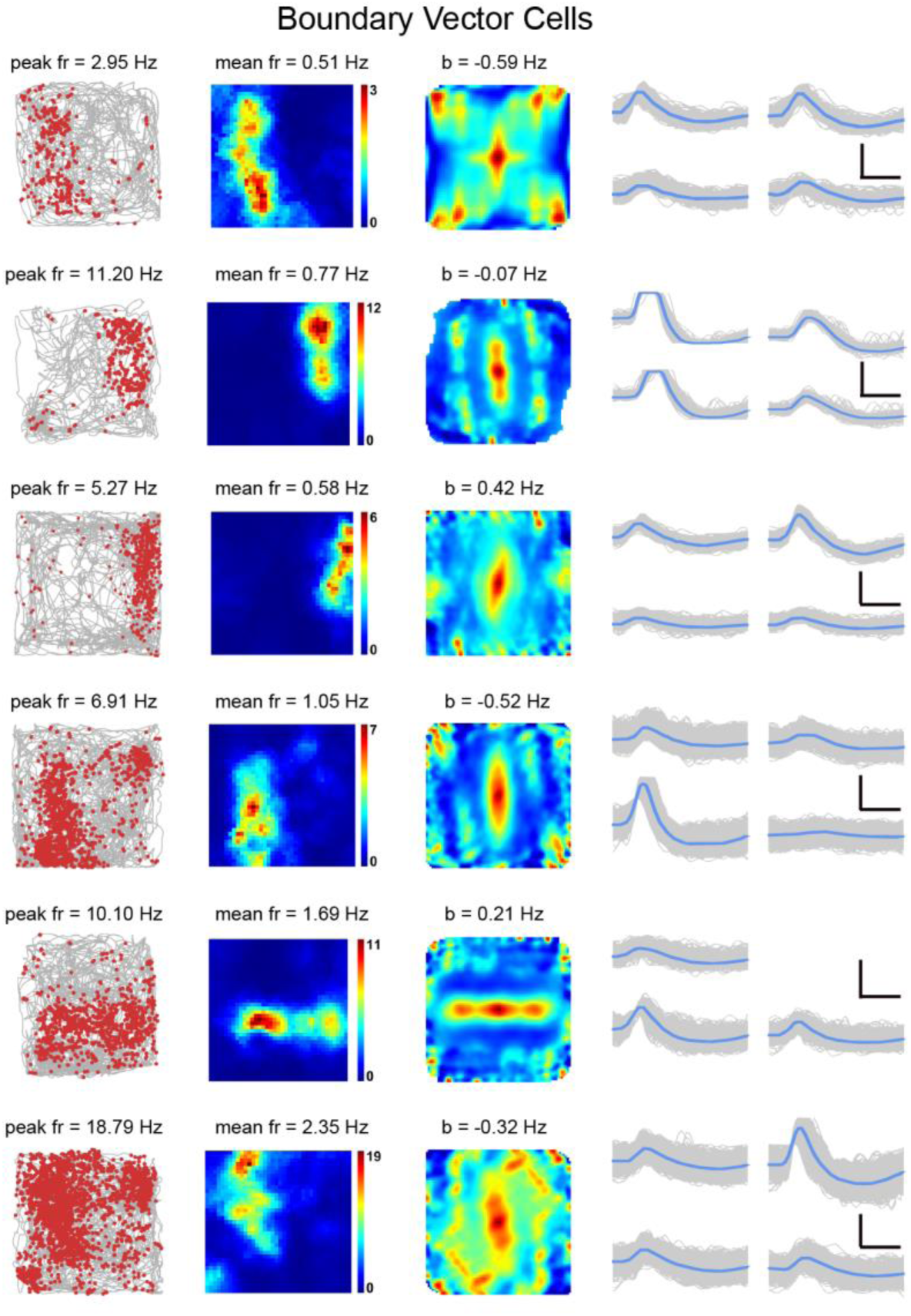
Somatosensory boundary vector cells recorded from the somatosensory cortex. Examples of somatosensory boundary vector cells. Trajectory (grey line) with superimposed spike locations (red dots) (left column); heat maps of firing rate (middle column) and autocorrelation diagrams (right column). Firing rate is color-coded with blue indicating minimum firing rate and red indicating maximum firing rate. The scale of the autocorrelation maps is twice that of the spatial firing rate maps. Peak firing rate (fr), mean firing rate (fr) and border score (b) for each representative boundary vector cell are labelled at the top of the panels. Spike waveforms on four electrodes are shown on the right column. Scale bar, 150 µV, 300 µs.

**Fig. S27.**
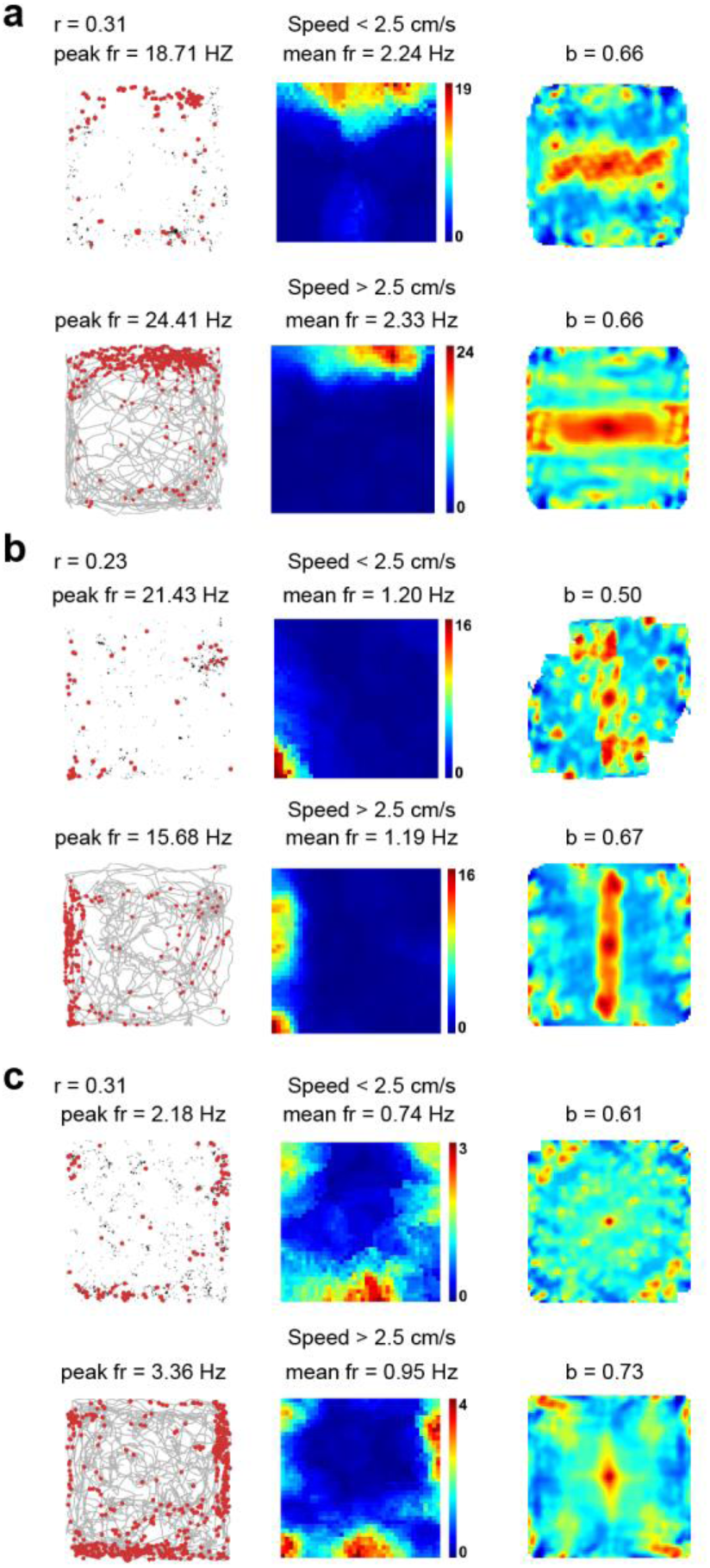
Comparison of the spatial response of somatosensory border cells between active running and slow immobility. (**a-c**) Comparison of the spatial response of representative border cells from Fig. 3a with instantaneous running speeds <2.5 cm/s and instantaneous running speeds > 2.5 cm/s, respectively. Trajectory (grey line) with superimposed spike locations (red dots) (left column); rate maps (middle column) and autocorrelation maps (right column) for instantaneous running speeds <2.5 cm/s and instantaneous running speeds > 2.5 cm/s, respectively. Firing rate is color-coded with blue indicating minimum firing rate and red indicating maximum firing rate. The scale of the autocorrelation maps is twice that of the spatial firing rate maps. Peak firing rate (fr), mean firing rate (fr) and border score (b) are labelled at the top of the plots. Pearson’s correlation coefficients between two firing rate maps when the instantaneous running speeds <2.5 cm/s and instantaneous running speeds > 2.5 cm/s are indicated with *r* at the top-left corner.

**Fig. S28.**
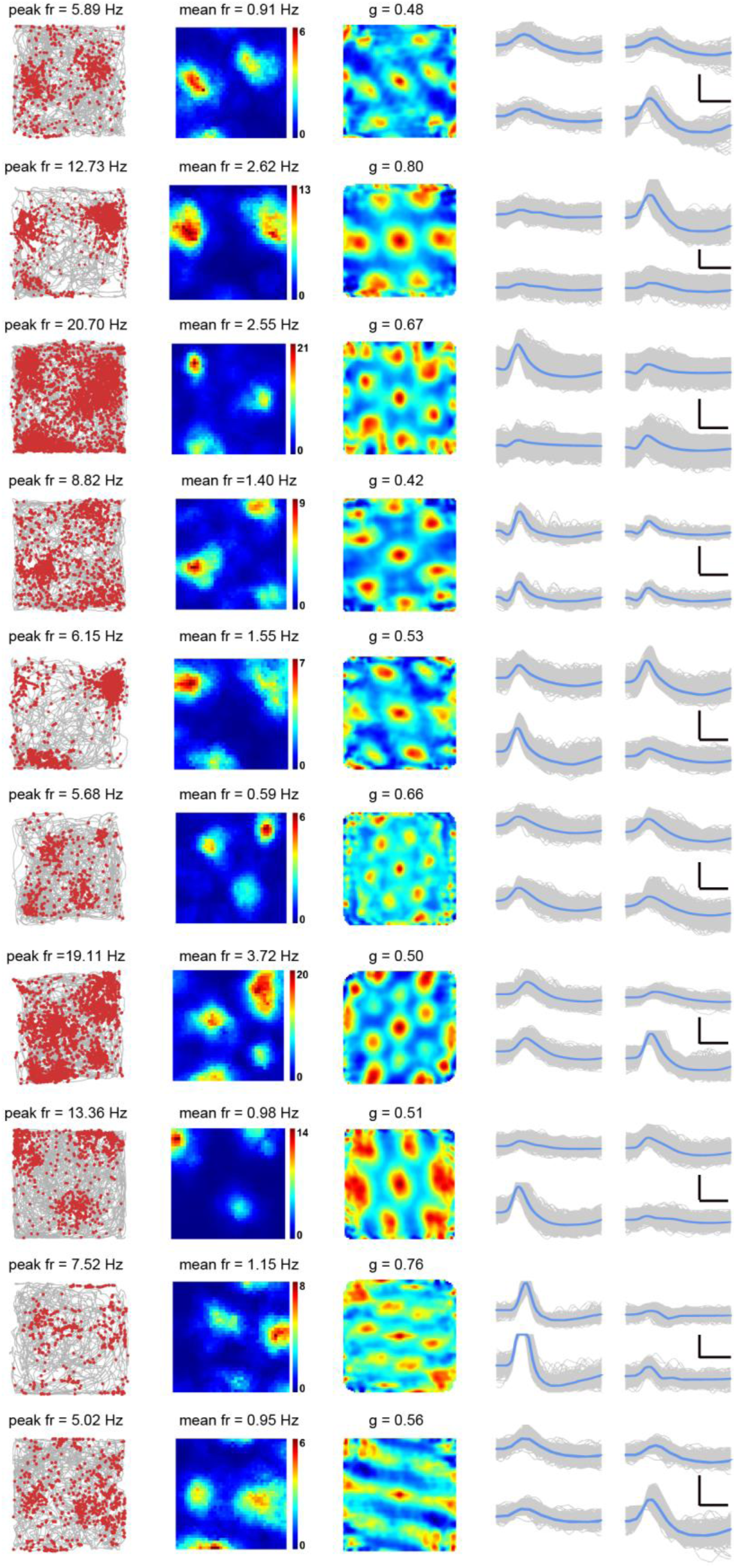
More examples of somatosensory grid cells recorded from the somatosensory cortex. Representative somatosensory grid cells with bin coverage over 90%. Trajectory (grey line) with superimposed spike locations (red dots) (left column); spatial firing rate maps (middle column) and autocorrelation diagrams (right column). Firing rate is color-coded with dark blue indicating minimal firing rate and dark red indicating maximal firing rate. The scale of the autocorrelation maps is twice that of the spatial firing rate maps. Peak firing rate (fr), mean firing rate (fr) and grid score (g) for each representative head direction cell are labelled at the top of the panels. The directional plots show strong head direction tuning. Spike waveforms on four electrodes are shown on the right column. Scale bar, 150 µV, 300 µs.

**Fig. S29.**
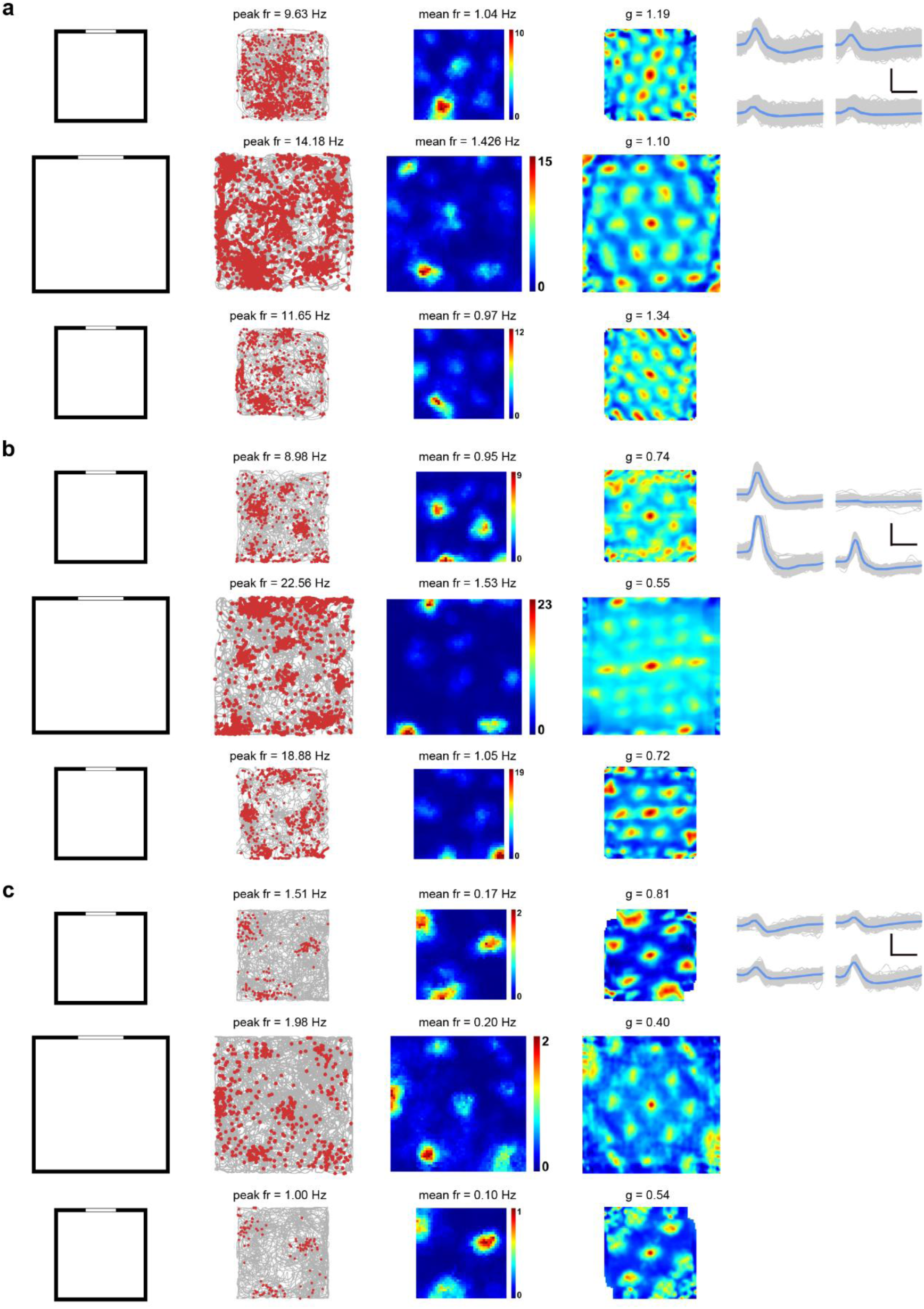
Somatosensory grid cells recorded in the larger environment. (**a-c**) Three examples of somatosensory grid cells recorded in 1 m x 1m, 1.5 m x 1.5 m and 1 m x 1m square box. The experimental diagram (left column); trajectory (grey line) with superimposed spike locations (red dots) (middle left column); spatial firing rate maps (middle right column) and autocorrelation diagrams (right column). Firing rate is color-coded with dark blue indicating minimal firing rate and dark red indicating maximal firing rate. The scale of the autocorrelation maps is twice that of the spatial firing rate maps. Peak firing rate (fr), mean firing rate (fr) and grid score (g) for each representative head direction cell are labelled at the top of the panels. Spike waveforms on four electrodes are shown on the right column. Scale bar, 150 µV, 300 µs.

**Fig. S30.**
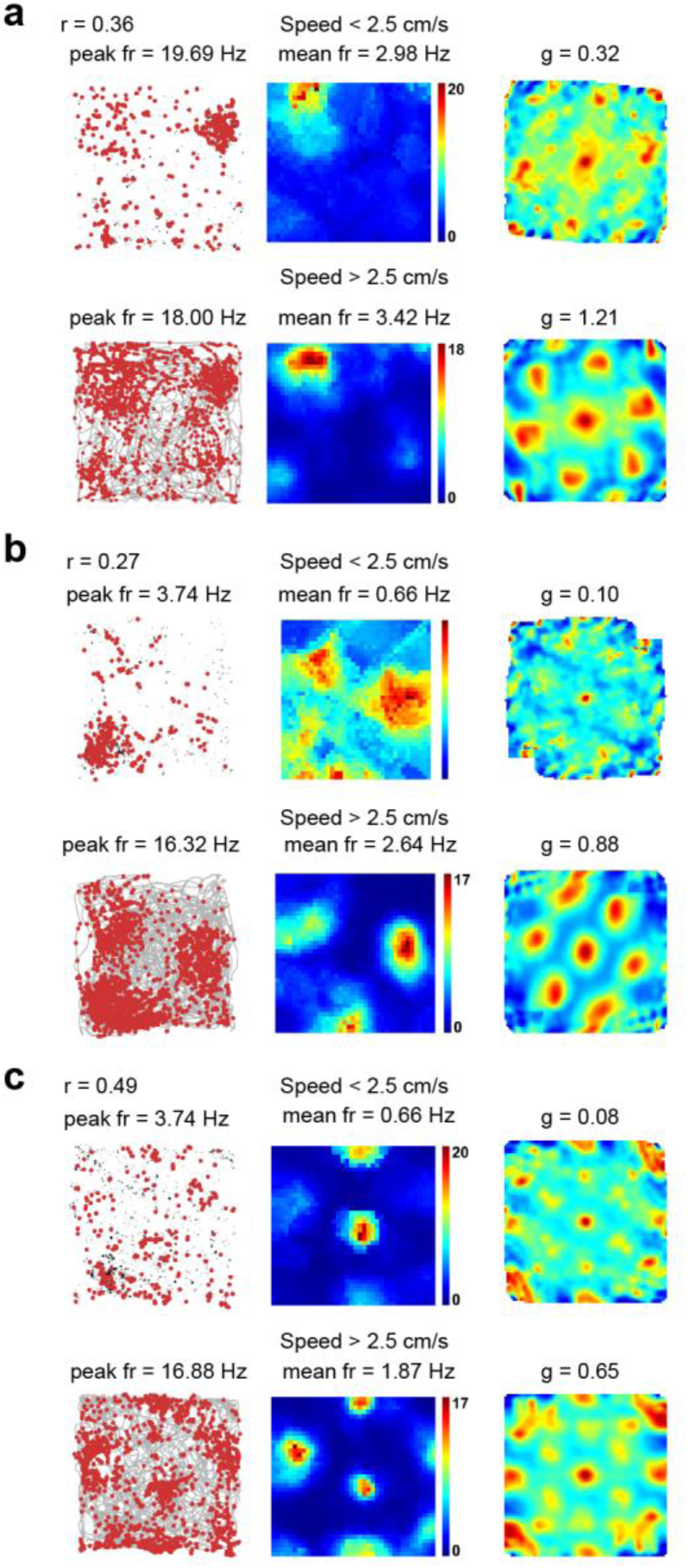
Comparison of the spatial response of somatosensory grid cells between active running and slow immobility. (**a-c**) Comparison of the spatial response of representative grid cells from Fig. 4a with instantaneous running speeds <2.5 cm/s and instantaneous running speeds > 2.5 cm/s, respectively. Trajectory (grey line) with superimposed spike locations (red dots) (left column); rate maps (middle column) and autocorrelation maps (right column) for instantaneous running speeds <2.5 cm/s and instantaneous running speeds > 2.5 cm/s, respectively. Firing rate is color-coded with blue indicating minimum firing rate and red indicating maximum firing rate. The scale of the autocorrelation maps is twice that of the spatial firing rate maps. Peak firing rate (fr), mean firing rate (fr) and grid score (b) are labelled at the top of the plots. Pearson’s correlation coefficients between two firing rate maps when the instantaneous running speeds <2.5 cm/s and instantaneous running speeds > 2.5 cm/s are indicated with *r* at the top-left corner.

**Fig. S31.**
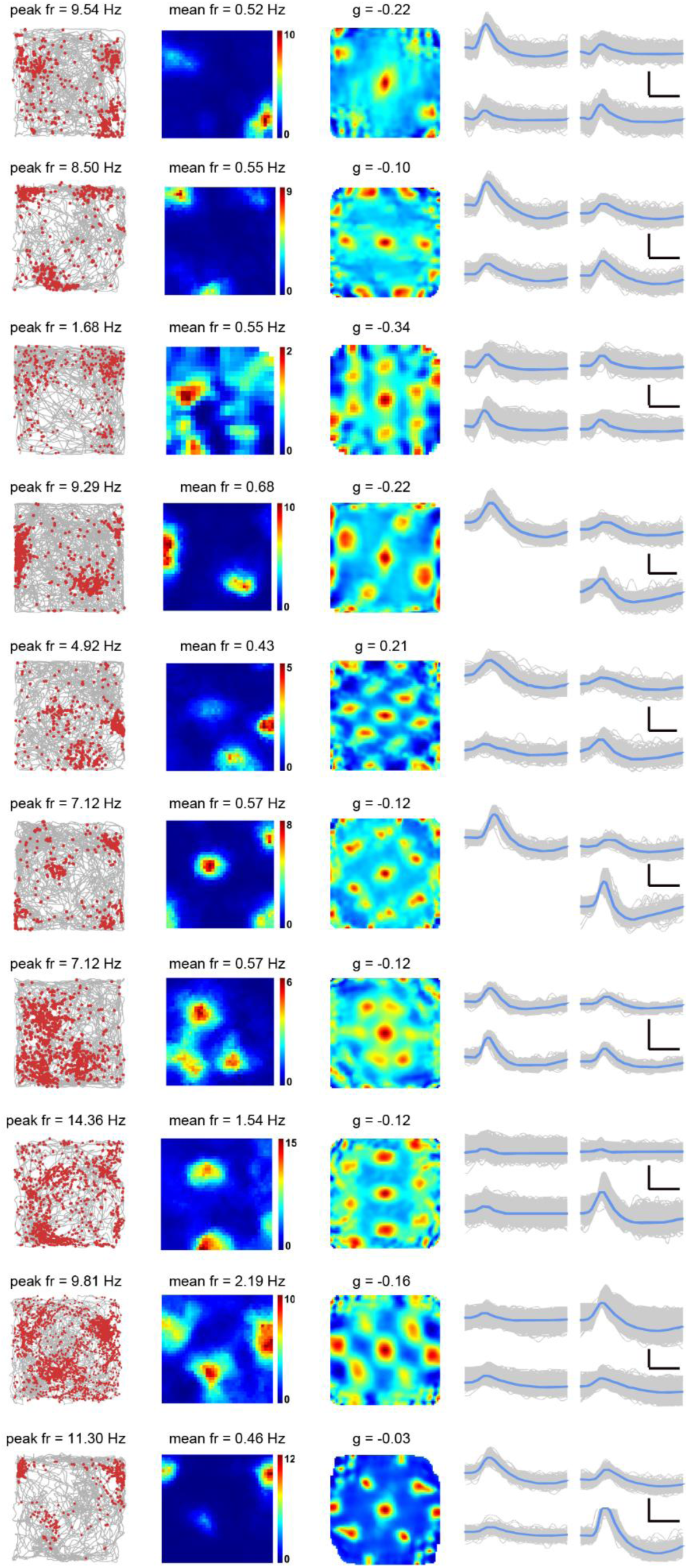
More examples of irregular grid cells recorded from the somatosensory cortex. Representative somatosensory irregular grid cells with bin coverage over 90%. Trajectory (grey line) with superimposed spike locations (red dots) (left column); spatial firing rate maps (middle column) and autocorrelation diagrams (right column). Firing rate is color-coded with dark blue indicating minimal firing rate and dark red indicating maximal firing rate. The scale of the autocorrelation maps is twice that of the spatial firing rate maps. Peak firing rate (fr), mean firing rate (fr) and grid score (g) for each representative head direction cell are labelled at the top of the panels. The directional plots show strong head direction tuning. Spike waveforms on four electrodes are shown on the right column. Scale bar, 150 µV, 300 µs.

**Fig. S32.**
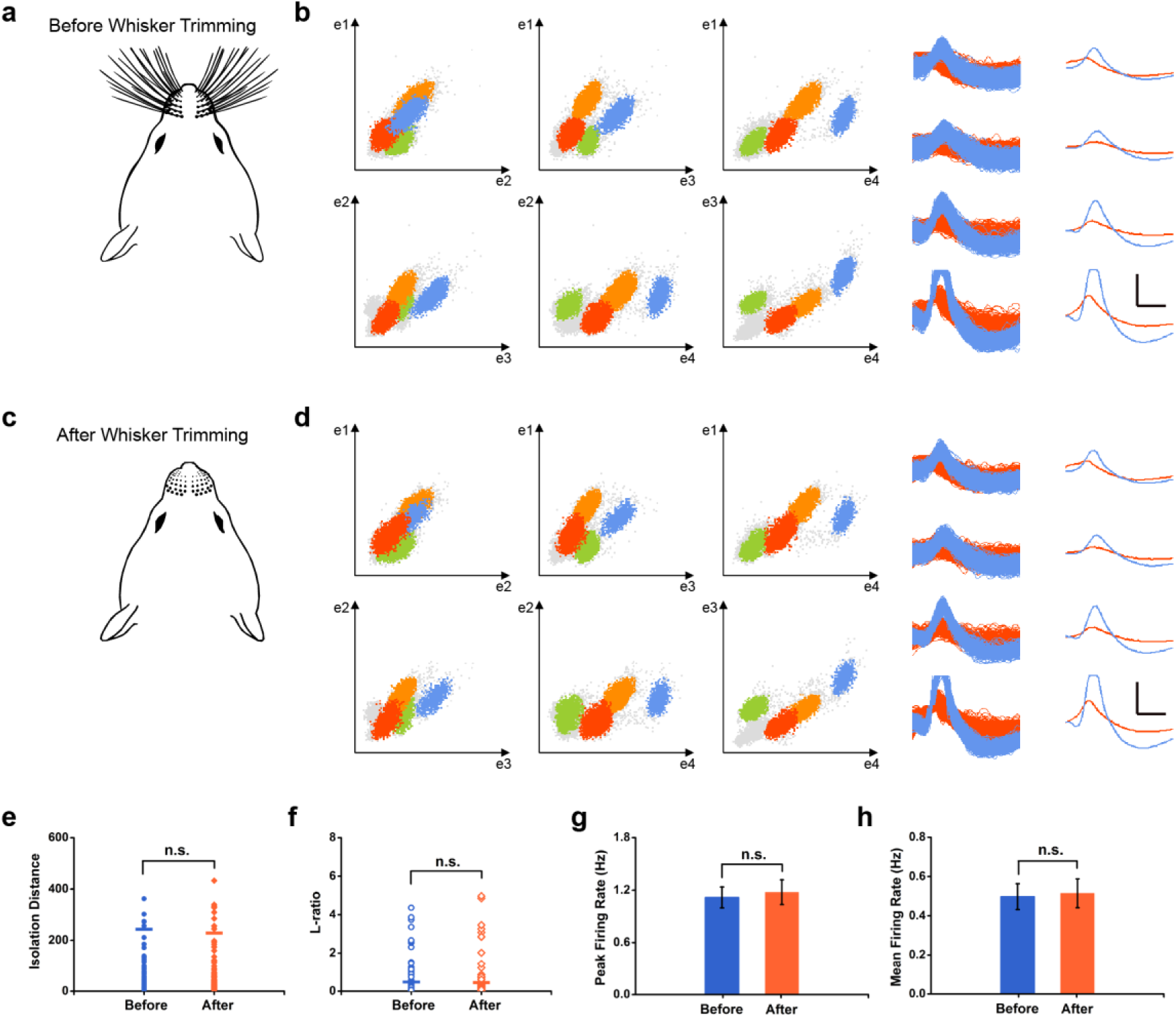
Preserved firing properties of the isolated somatosensory units before and after whisker trimming. (**a** and **c**) The diagram showing the rat before (**a**) and after (**c**) whisker trimming. (**b** and **d**) Scatterplots show preserved cluster diagrams before (**b**) and after (**d**) whisker trimming of the same tetrode from the same recording rat. Each individual dot represents a single recorded spike. Waveforms from two separated blue and red clusters in the scatterplots are shown for four electrodes from the same tetrode. The waveforms remain unchanged after whisker trimming. Scale bar, 150 µV, 300 µs. (**e**) The comparison of isolation distance for identified S1 units right before and after whisker trimming. (**f**) Same as **e** for the L-ratio. (**g**) Same as in (**f)** for the mean firing rate. (**h**) Same as in (**g)** for the peak firing rate. *n* = 78, two-tailed paired *t-*test, n.s., not significant.

**Fig. S33.**
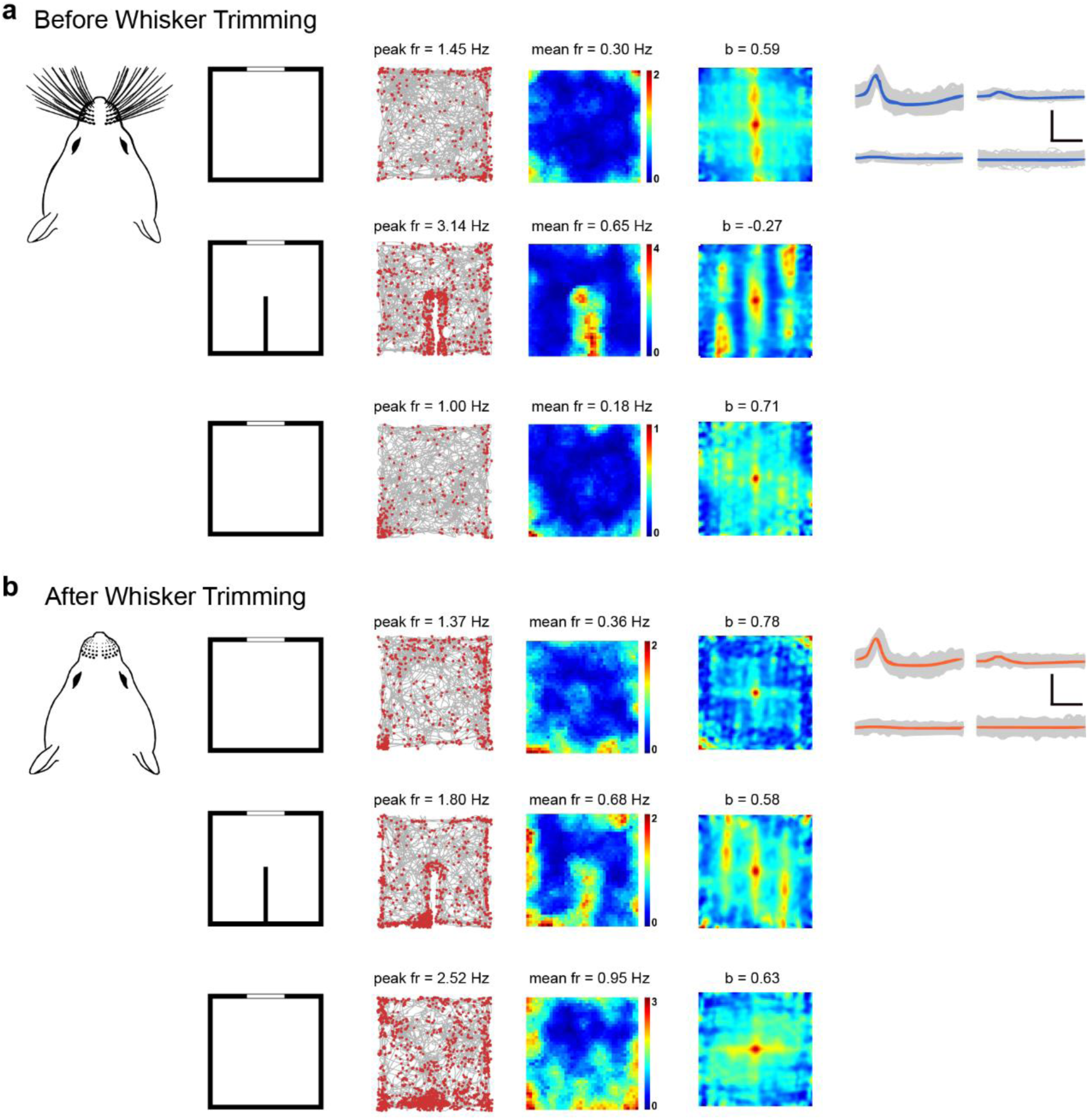
Preserved spatial firing properties of the same somatosensory border cell before and after whisker trimming. (**a** and **b**) The diagram showing the rat before (**a**) and after (**b**) whisker trimming. A representative somatosensory border cell preserves the firing patterns in the square box with (top and bottom panels) and without (middle panels) the internal insert before and after whisker trimming. The experimental diagram (left column); trajectory (grey line) with superimposed spike locations (red dots) (middle left column); rate maps (middle right column) and autocorrelation maps (right column) for each recording trial from the same S1 border cell. Firing rate is color-coded with blue indicating minimum firing rate and red indicating maximum firing rate. The scale of the autocorrelation maps is twice that of the spatial firing rate maps. Peak firing rate (fr), mean firing rate (fr) and border score (b) for each recording session are labelled at the top of the panels. Spike waveforms on four electrodes are shown on the right column. Scale bar, 100 µV, 300 µs.

**Table S1.** Summary of somatosensory spatial cell types recorded in each rat.

|  | Somatosensory Spatial Cell Types |  |  |  |  |
| --- | --- | --- | --- | --- | --- |
| Rat Number | Place Cells | Head Direction Cells | Border Cells | Grid Cells | Total Cells |
| 1 (S1HL) | 22 | 13 | 6 | 12 | 182 |
| 2 (S1HL) | 27 | 15 | 14 | 13 | 312 |
| 3 (S1FL) | 19 | 9 | 8 | 10 | 353 |
| 4 (S1HL) | 20 | 7 | 12 | 9 | 275 |
| 5 (S1Sh) | 20 | 6 | 8 | 12 | 263 |
| 6 (S1HL) | 35 | 8 | 16 | 7 | 197 |
| 7 (S1HL) | 36 | 8 | 12 | 5 | 301 |
| 8 (S1HL) | 16 | 14 | 10 | 4 | 142 |
| **Total** | 195 | 80 | 86 | 72 | 2025 |

